# Adaptation of bacteria to glyphosate: a microevolutionary perspective of the enzyme 5-enolpyruvylshikimate-3-phosphate (EPSP) synthase

**DOI:** 10.1101/2020.06.16.154005

**Authors:** Miia J. Rainio, Suvi Ruuskanen, Marjo Helander, Kari Saikkonen, Irma Saloniemi, Pere Puigbò

**Affiliations:** Department of Biology, University of Turku, Turku, Finland; Biodiversity Unit, University of Turku, Finland; Nutrition and Health Unit, Eurecat Technology Centre of Catalonia, Reus, Catalonia, Spain

**Keywords:** bacteria, antibiotic resistance, microevolution, glyphosate, epsps, shikimate pathway

## Abstract

Glyphosate is the leading herbicide worldwide, but it also affects prokaryotes because it targets the central enzyme (EPSPS) of the shikimate pathway in the synthesis of the three essential aromatic amino acids in autotrophs. Our results reveal that bacteria easily become resistant to glyphosate through changes in the EPSPS active site. This indicates the importance of examining how glyphosate affects microbe-mediated ecosystem functions and human microbiomes.

---

Evolutionarily conserved 5-enolpyruvylshikimate-3-phosphate synthase (EPSPS) is the key enzyme of the shikimate pathway in the synthesis of three essential aromatic amino acids (phenylalanine, tyrosine and tryptophan) across taxa, including bacteria, fungi and plants ^1,2^. Because the shikimate pathway does not occur in vertebrates, the use of glyphosate, targeting the EPSPS enzyme by competing for the binding site with the second substrate of the EPSPS (phosphoenol pyruvate, PEP) ^3^, is regarded safe for use in food production. Due to its affordable price, effectiveness and broad-spectrum ability to kill weeds, glyphosate has become the most commonly used herbicide worldwide ^4–6^. Although glyphosate antibiotic properties are known ^7,8^, its possible effects on microbiomes ^9–11^ have largely been neglected until recently ^10,12^. As microbes have driven eco-evolutionary processes since the origin of life, we note the importance of thoroughly understanding the possible undesirable effects of glyphosate on ecosystem structures, functions and services ^13^. Here, we propose that the wide use of glyphosate may potentially affect microbial communities by (i) eradicating susceptible microbes and (ii) leading to evolution of resistance. We analyzed 663 microbial genomes distributed across 32 alignable tight genomic clusters (ATGCs) ^14^, with the *EPSPSClass* server ^15^ to identify changes in sensitivity to glyphosate. The algorithm classifies EPSPS proteins based on the presence and absence of amino acid markers in the active site ^15^ (see methods).

A microevolutionary analysis of the EPSPS enzyme shows that phylogenetics best explains bacterial sensitivity to glyphosate (figure 1 and supplementary tables 1-2). In general, Firmicutes are significantly more resistant to glyphosate than Proteobacteria, whereas Actinobacteria is the most sensitive group to glyphosate. Furthermore, bacterial lifestyle is also associated with sensitivity, i.e., facultative host-associated (FHA) bacteria are more sensitive to glyphosate than free-living (FL) bacteria (supplementary tables 3-4 and supplementary figure 1). Intracellular parasites seem to be the most sensitive group; however, due to the limited amount of data, we cannot determine statistical significance for this group’s sensitivity. We hypothesize that sensitivity mirrors the strength of selective force of glyphosate on bacteria. FL bacteria are directly exposed to glyphosate spray at high concentrations, whereas host-associated microbes can be exposed to lower concentrations when the glyphosate is moving through the plant from leaves to roots, and intracellular parasites might be protected by the cell membrane.. Genomes with %GC below 40% tend to be more resistant to glyphosate (supplementary figure 2). Spearman correlation coefficients between sensitivity and genome parameters from the original ATGC dataset ^14^ were not statistically significant (see methods and supplementary table 5)

**Figure 1.**
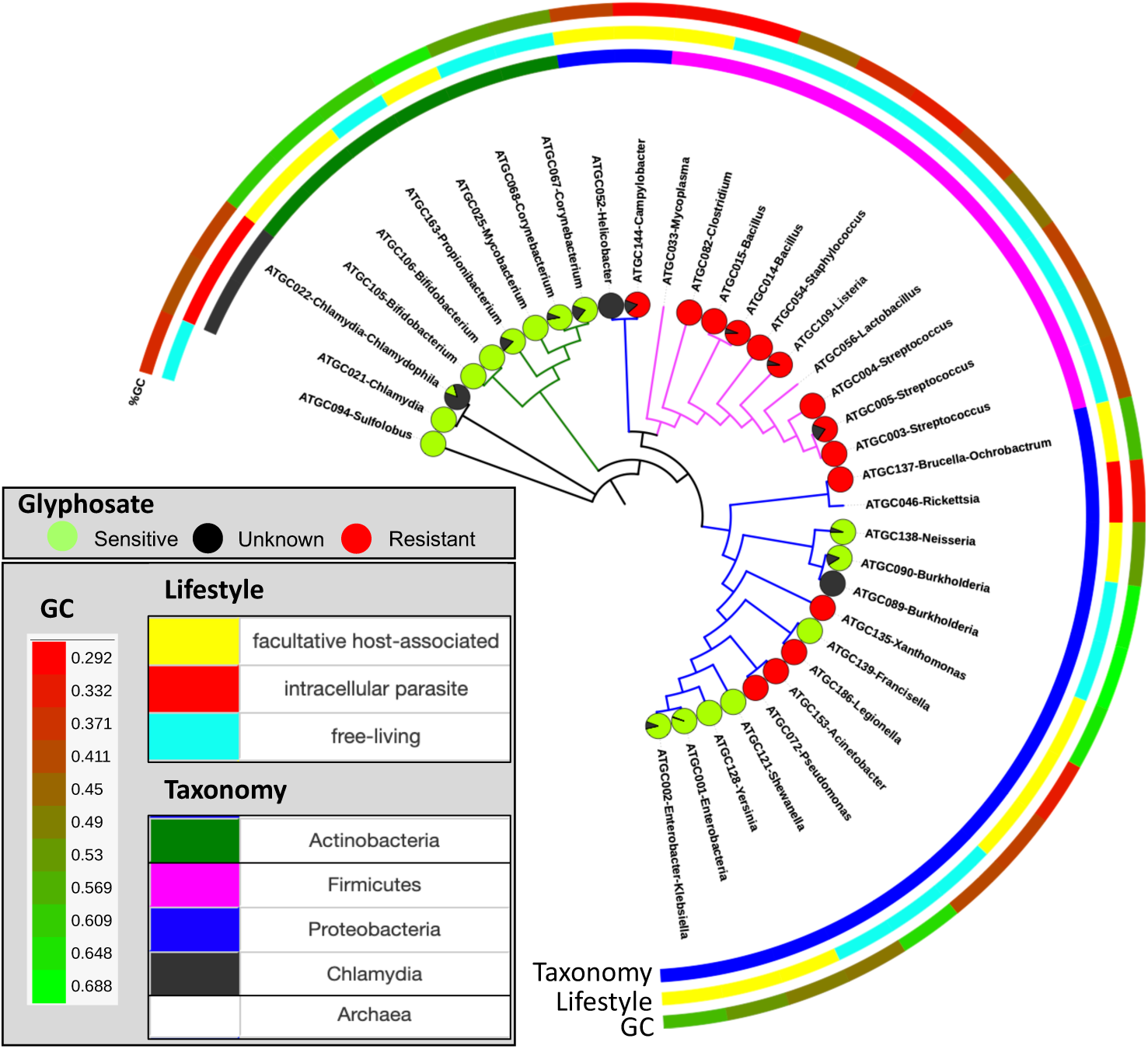
Distribution of the sensitivity to glyphosate across the species tree. See full description in supplementary table 1.

EPSPS is the central enzyme in the metabolic pathway for the synthesis of three essential amino acids in bacteria ^16^; thus, its protein sequence is presumably evolutionarily conserved ^17^. Indeed, the EPSPS sequence is highly conserved within 17 ATGCs that preserve sensitivity status across species. However, 12 ATGCs contained species with different sensitivities to glyphosate (figure 2, supplementary figures 3-14 and supplementary table 1). Although the active site of the EPSPS is highly conserved in prokaryotes, in some species, the sensitivity to glyphosate may be altered with few changes in the active site (figure 2). In Enterobacteria (ATGC001), only *Enterobacter* R4 368 (out of 109 species) is putatively resistant to glyphosate after recently acquiring a second copy of the EPSPS *via* horizontal gene transfer (HGT) from a resistant strain of *Klebsiella pneumoniae* (supplementary figure 15). Other Proteobacteria clusters have species with differential sensitivity status after few amino acid changes in the active site (figure 2). This microevolutionary analysis shows that some bacteria may become putatively resistant to glyphosate through small changes, analogous to the antibiotic resistance mechanism ^7,13^. Thus, the wide use of glyphosate may have a very large impact on the species diversity and composition of microbial communities not only because of a potential purifying selection effect against sensitive bacteria but also because (i) some bacterial groups may adapt rapidly to become resistant to glyphosate and (ii) glyphosate-based herbicides may enhance multidrug resistance in bacteria ^18^.

**Figure 2.**
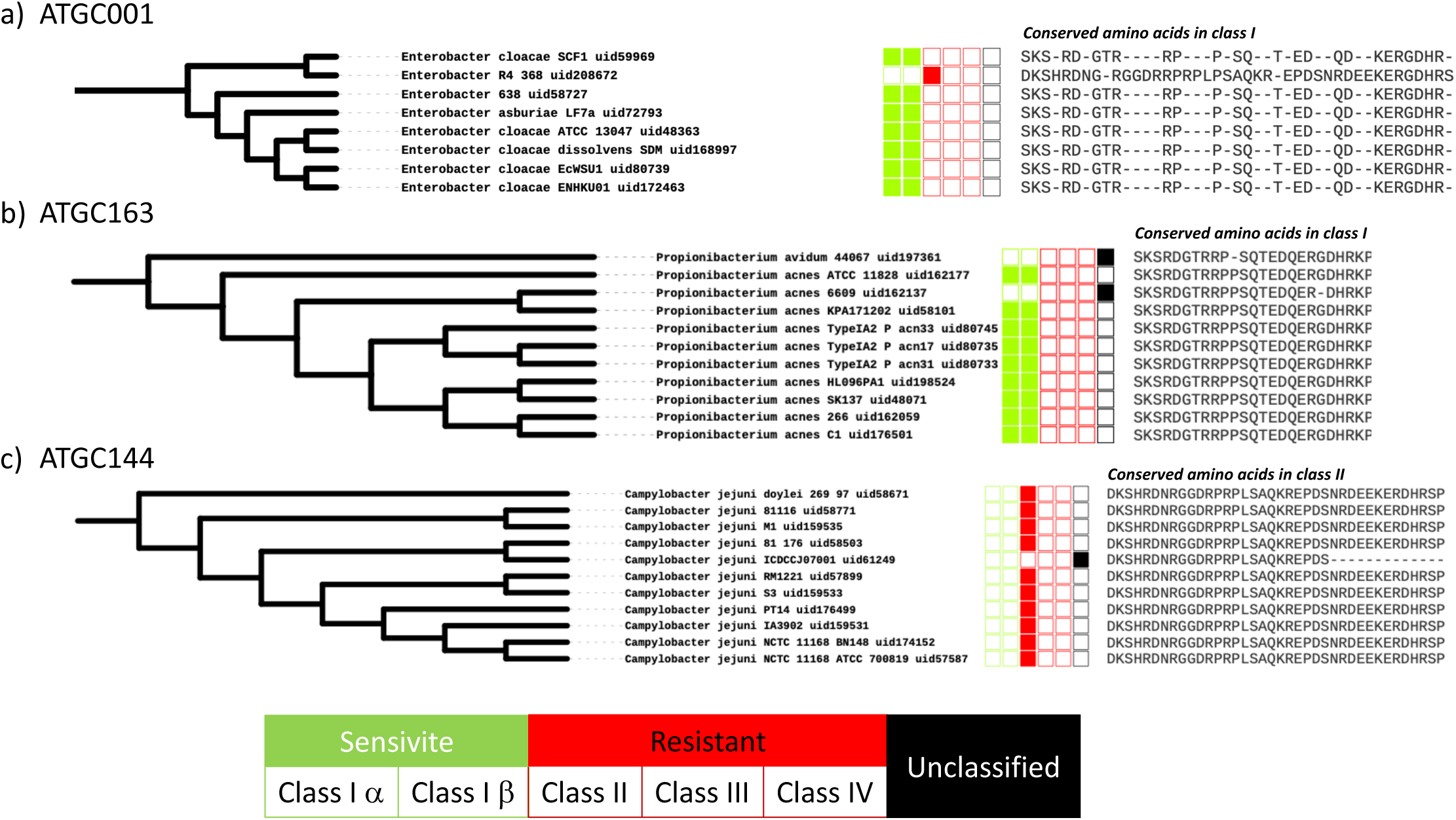
Differential sensitivity to glyphosate in three Proteobacteria ATGCs. a) Change from sensitive to resistant in *Enterobacter* R4 368 through HGT b) Two independent changes from sensitive to unclassified in propionibacterium with single amino acid changes c) Change from resistant to unclassified in Campylobacter with several amino acid changes

An analysis of variability in the amino acid landscape (see methods) shows that the amino acids in the active site of the EPSPS that bind PEP are 23% more conserved than the rest of the amino acid residues (supplementary figure 16), in agreement with results from previous studies ^3^. Moreover, amino acids flanking binding sites (+/- 5 residues) also show higher levels of conservation than average, possibly due to selection preserving the function of the EPSPS. Bacteria more resistant to glyphosate tend to have fewer amino acid substitutions than sensitive ones, e.g., the binding amino acids of actinobacteria and FL bacteria (more resistant to glyphosate) tend to be more conserved (supplementary figure 17 and supplementary table 6-7). Although there are homogeneous and heterogeneous clusters in relation to glyphosate sensitivity, the active site is equally conserved (supplementary table 8). Further analyses of amino acid variabilities in EPSPS (supplementary table 9) should help elucidate the effects of glyphosate on microbial communities.

In conclusion, this study shows that taxonomic groups and bacterial lifestyles are key factors determining sensitivity to glyphosate. Our results suggest that the sensitivity of bacterial species to glyphosate varies and can change in the short evolutionary time of ATGC. Moreover, microbes more exposed to the herbicide (FL bacteria) are putatively more resistant. Nevertheless, because single mutations in the EPSPS active site and rapid HGT events may change the status of sensitivity, heavy use of glyphosate may impact microbial biodiversity independently of taxonomy and lifestyle. The central and unanswered questions are (i) what are the effects of glyphosate-modulated microbiomes on ecosystem functions and services and (ii) on human wellbeing.

## METHODS

### Protein sequences

Protein data were obtained from the ATGC database, which contains data on >4.5 million proteins encoded in >1,500 genomes of prokaryotes (approximately 60% of proteins and 62% of genomes from RefSeq as of June 2013) that met the same criteria as the original ATGCs ^19^, as described in ^14^. EPSPS protein sequences were annotated through BLAST mapping onto the Clusters of Orthologous Genes (COG) database ^20^. EPSPS belongs to COG0128 (category E, amino acid transport and metabolism). We analyzed 32 ATGCs that amply represented bacteria and one archaeal group. The EPSPS sequence was not found in ATGC033 (mycoplasma), a small-genome membrane-free bacteria ^21^; ATGC046 (rickettsia), an obligate intracellular parasite ^22^; and ATGC056 (lactobacillus). Even though lactobacillus strains in the ATGC dataset do not have the EPSPS sequence, other strains have the enzyme and present differential sensitivity to glyphosate (supplementary figure 10). Previous studies have shown that some Lactobacillus species are susceptible to glyphosate ^23^. Each ATGC contains at least 10 species by the definition in the original study ^14^, and up to 109 species are present in the enterobacterial cluster (ATGC001). The final ATGC dataset used in this study includes 332 FHA species (13 ATGCs), 312 FL species (17 ATGCs), and 19 intracellular parasites (2 ATGCs).

### Classification of the EPSPS

Glyphosate inhibits the shikimate pathway by inhibiting the enzyme 5-enolpyruvylshikimate-3-phosphate synthase (EPSPS) ^3,24^. The EPSPS class has been classified with the *EPSPSClass* server (http://ppuigbo.me/programs/EPSPSClass), which assesses the type of potential sensitivity of the EPSPS enzyme to the herbicide glyphosate ^15^. PSPS enzymes can be classified as class I (alpha or beta) ^25,26^, class II ^27,28^, III ^29^ or IV ^30^ based on the presence of amino acid markers (class I, class II and class IV) and motifs (class III). In addition, a large portion of microbial species are yet unclassified, and their potential sensitivities to glyphosate are unknown ^15^. Different uses of amino acid markers distinguish the EPSPS enzymes into four classes using the reference EPSPS enzymes from *Vibrio cholerae serotype* O1 (vcEPSPS, class I), *Coxiella burnetii* (cbEPSPS, class II), *Brevundimonas vesicularis* (bvEPSPS, class III) and *Streptomyces davawensis* (sdEPSPS, class IV).

### Analysis of the amino acid substitution landscape

We have performed a protein sequence alignment of the EPSPS sequences from the ATGCs and the reference EPSPS of *Vibrio cholerae serotype* O1 (vcEPSPS; Class I ^25,26^; supplementary table 6) with the program Muscle ^31^. The vcEPSPS sequence was used to determine the location of residues in the active site that interact directly with PEP ^3^. Glyphosate inhibits EPSPS by competing with PEP ^3^. We counted the number of alternative amino acids in each position of the alignment to determine the degree of amino acid substitution in the active site and the rest of the residues. We determined an active site region, i.e., a wider area around the active site residues (+/- 5 amino acids) that interacts with PEP.

### Statistical analysis

The tested Spearman correlation coefficients between sensitivity and total genome dynamics (as the sum of gains, losses, expansions and reductions; GLER), genome number (genome no.), genome size, protein-coding genes, gene families, median gene cluster content, median guanine+cytosine (median GC), median dN/dS, synteny distance (dY) and percentage of aromatic amino acids were not statistically significant (p-value > 0.05) (supplementary table 5). We performed a chi-squared test to determine associations between sensitivity to glyphosate and lifestyle (FL vs FHA) and taxonomic group (Actinobacteria, Firmicutes and Proteobacteria). A Kruskal-Wallis test was used to analyze conservation of the amino acids in the EPSPS

## ACKNOWLEDGMENTS

Pere Puigbò’s research is supported by funds from the Turku Collegium for Science and Medicine (Turku, Finland) and by the AEI (Grant no. PTQ2018-009846 to Pere Puigbo). Miia Rainio was funded by the Academy of Finland (grant no. 311077 to Marjo Helander).

## Notes

http://ppuigbo.me/programs/EPSPSClass/

### Competing Interest Statement

The authors have declared no competing interest.

