## Supplementary Material for "Adaptation of bacteria to glyphosate: a microevolutionary perspective of the enzyme 5-enolpyruvylshikimate-3-phosphate (EPSP) synthase"

### SUPPLEMENTARY TABLES

**Supplementary table 1.** Summary table of the percentage of species sensitive / resistant to glyphosate. ATGCs containing species with potential differential sensitivity to glyphosate are highlighted in blue.

| ATGC | Genera | Taxonomy | Lifestyle | N | S | CDS | G | GC | A | P | Class I | Class II | Class III | Class IV | Unknown | % Sensitive Species |
|---|---|---|---|---|---|---|---|---|---|---|---|---|---|---|---|---|
| ATGC001 | *Enterobacteria* | Proteobacteria | FHA | 109 | 4989733 | 4652 | 4215 | 0.528 | 12688949 | 8.24% | 109 | 1 | 0 | 0 | 0 | **99%** |
| ATGC002 | *Enterobacter–Klebsiella* | Proteobacteria | FHA | 11 | 5680456 | 5375 | 5234 | 0.583 | 1430903 | 8.16% | 10 | 0 | 0 | 0 | 1 | **91%** |
| ATGC003 | *Streptococcus* | Firmicutes | FL | 22 | 2133789 | 2114 | 2020 | 0.408 | 1208968 | 9.37% | 0 | 22 | 0 | 0 | 0 | **0%** |
| ATGC004 | *Streptococcus* | Firmicutes | FL | 22 | 1910066 | 1865 | 1808 | 0.395 | 1065353 | 8.97% | 0 | 22 | 0 | 0 | 0 | **0%** |
| ATGC005 | *Streptococcus* | Firmicutes | FL | 16 | 2084900 | 1977 | 1948 | 0.422 | 860870 | 9.21% | 0 | 15 | 0 | 0 | 4 | **0%** |
| ATGC014 | *Bacillus* | Firmicutes | FL | 31 | 5691541 | 5704 | 5533 | 0.361 | 4463047 | 9.42% | 0 | 30 | 0 | 0 | 2 | **0%** |
| ATGC015 | *Bacillus* | Firmicutes | FL | 24 | 4063283 | 4089 | 3976 | 0.459 | 2481021 | 8.91% | 0 | 24 | 0 | 0 | 0 | **0%** |
| ATGC021 | *Chlamydia* | Chlamydia | P | 44 | 1046020 | 898 | 901 | 0.417 | 1215063 | 8.87% | 44 | 0 | 0 | 0 | 0 | **100%** |
| ATGC022 | *Chlamydia–Chlamydophila* | Chlamydia | P | 19 | 1169029 | 1126 | 1011 | 0.397 | 587259 | 8.93% | 3 | 0 | 0 | 0 | 17 | **15%** |
| ATGC025 | *Mycobacterium* | Actinobacteria | FHA | 32 | 4409130 | 3981 | 3836 | 0.659 | 2636219 | 6.53% | 34 | 0 | 0 | 0 | 0 | **100%** |
| ATGC052 | *Helicobacter* | Proteobacteria | FHA | 51 | 1628207 | 1564 | 1501 | 0.397 | 2390676 | 9.80% | 0 | 0 | 0 | 0 | 51 | **0%** |
| ATGC054 | *Staphylococcus* | Firmicutes | FL | 41 | 2837953 | 2624 | 2452 | 0.337 | 2881366 | 9.15% | 0 | 41 | 0 | 0 | 0 | **0%** |
| ATGC067 | *Corynebacterium* | Actinobacteria | FL | 15 | 2322675 | 2089 | 2074 | 0.530 | 690404 | 7.06% | 12 | 0 | 0 | 0 | 3 | **80%** |
| ATGC068 | *Corynebacterium* | Actinobacteria | FL | 12 | 2466919 | 2262 | 2227 | 0.543 | 602885 | 7.05% | 11 | 0 | 0 | 0 | 1 | **92%** |
| ATGC072 | *Pseudomonas* | Proteobacteria | FL | 12 | 6095696 | 5386 | 5297 | 0.633 | 1536547 | 7.52% | 0 | 12 | 0 | 0 | 0 | **0%** |
| ATGC082 | *Clostridium* | Firmicutes | FL | 10 | 4025029 | 3691 | 3655 | 0.292 | 966729 | 9.20% | 0 | 10 | 0 | 0 | 0 | **0%** |
| ATGC089 | *Burkholderia* | Proteobacteria | FL | 13 | 6406482 | 5743 | 5707 | 0.688 | 1676489 | 7.30% | 0 | 0 | 0 | 0 | 25 | **0%** |
| ATGC090 | *Burkholderia* | Proteobacteria | FL | 12 | 7530187 | 6697 | 6428 | 0.678 | 1844146 | 7.43% | 12 | 0 | 0 | 0 | 2 | **86%** |
| ATGC094 | *Sulfolobus* | Archaea | FL | 12 | 2686155 | 2735 | 2638 | 0.356 | 915829 | 10.20% | 12 | 0 | 0 | 0 | 0 | **100%** |
| ATGC105 | *Bifidobacterium* | Actinobacteria | FHA | 11 | 2454110 | 1972 | 1952 | 0.609 | 558314 | 7.52% | 11 | 0 | 0 | 0 | 0 | **100%** |
| ATGC106 | *Bifidobacterium* | Actinobacteria | FHA | 10 | 1938894 | 1567 | 1568 | 0.615 | 412746 | 7.47% | 10 | 0 | 0 | 0 | 0 | **100%** |
| ATGC109 | *Listeria* | Firmicutes | FL | 31 | 2934366 | 2891 | 2843 | 0.385 | 2369308 | 8.93% | 0 | 29 | 0 | 0 | 2 | **0%** |
| ATGC121 | *Shewanella* | Proteobacteria | FL | 14 | 5125160 | 4360 | 4166 | 0.475 | 1604716 | 8.22% | 14 | 0 | 0 | 0 | 0 | **100%** |
| ATGC128 | *Yersinia* | Proteobacteria | FHA | 19 | 4744469 | 4114 | 3934 | 0.489 | 1966701 | 8.12% | 19 | 0 | 0 | 0 | 0 | **100%** |

| | | | | | | | | | | | | | | | | |
|---|---|---|---|---|---|---|---|---|---|---|---|---|---|---|---|---|
| ATGC135 | *Xanthomonas* | Proteobacteria | FHA | 13 | 5119661 | 4447 | 4204 | 0.654 | 1312376 | 7.19% | 0 | 13 | 0 | 0 | 0 | **0%** |
| ATGC137 | *Brucella–Ochrobactrum* | Proteobacteria | FHA | 19 | 3407297 | 3268 | 3212 | 0.584 | 1343990 | 7.59% | 0 | 19 | 0 | 0 | 0 | **0%** |
| ATGC138 | *Neisseria* | Proteobacteria | FHA | 18 | 2218847 | 1961 | 1937 | 0.537 | 857313 | 8.27% | 17 | 0 | 0 | 0 | 1 | **94%** |
| ATGC139 | *Francisella* | Proteobacteria | FHA | 18 | 1916517 | 1634 | 1613 | 0.333 | 905217 | 9.93% | 18 | 0 | 0 | 0 | 0 | **100%** |
| ATGC144 | *Campylobacter* | Proteobacteria | FHA | 11 | 1690063 | 1646 | 1650 | 0.309 | 570516 | 10.31% | 0 | 10 | 0 | 0 | 2 | **0%** |
| ATGC153 | *Acinetobacter* | Proteobacteria | FL | 14 | 3959662 | 3759 | 3517 | 0.403 | 1350481 | 8.82% | 0 | 13 | 0 | 0 | 0 | **0%** |
| ATGC163 | *Propionibacterium* | Actinobacteria | FL | 11 | 2518661 | 2280 | 2270 | 0.608 | 514221 | 6.51% | 9 | 0 | 0 | 0 | 2 | **82%** |
| ATGC186 | *Legionella* | Proteobacteria | FHA | 10 | 3496368 | 3166 | 3038 | 0.391 | 920655 | 9.16% | 0 | 10 | 0 | 0 | 0 | **0%** |

FHA: facultative host-associated; FL: free-living; P: intracellular parasite.

CDS: number of protein coding genes; N: number of genomes; S: genome size (bp); G: number of gene families; GC: median GC; A: number of aromatic amino acids; P: percentage of aromatic amino acids.

**Supplementary table 2.** Predictions of type of EPSPS in the ATGCS

| ATGC | SPECIES | GI | REF | Class Ia | Class Ib | Class II | Class III | Class IV | Prediction |
|---|---|---|---|---|---|---|---|---|---|
| ATGC001 | Citrobacter_koseri_ATCC_BAA_895_uid58143 | 157146405 | YP_001453724.1 | 1 | 1 | 0.605 | 0/18 | 0.233 | Sensitive |
| ATGC001 | Citrobacter_rodentium_ICC168_uid43089 | 283784735 | YP_003364600.1 | 1 | 1 | 0.605 | 0/18 | 0.233 | Sensitive |
| ATGC001 | Enterobacter_638_uid58727 | 146311085 | YP_001176159.1 | 1 | 1 | 0.605 | 0/18 | 0.233 | Sensitive |
| ATGC001 | Enterobacter_R4_368_uid208672 | 512583468 | YP_008098272.1 | 0.8 | 0.889 | 1 | 0/18 | 0.167 | Resistant |
| ATGC001 | Enterobacter_R4_368_uid208672 | 512652300 | YP_008110400.1 | 1 | 1 | 0.605 | 0/18 | 0.233 | Sensitive |
| ATGC001 | Enterobacter_asburiae_LF7a_uid72793 | 345298586 | YP_004827944.1 | 1 | 1 | 0.605 | 0/18 | 0.233 | Sensitive |
| ATGC001 | Enterobacter_cloacae_ATCC_13047_uid48363 | 296103089 | YP_003613235.1 | 1 | 1 | 0.605 | 0/18 | 0.233 | Sensitive |
| ATGC001 | Enterobacter_cloacae_ENHKU01_uid172463 | 401763025 | YP_006578032.1 | 1 | 1 | 0.605 | 0/18 | 0.233 | Sensitive |
| ATGC001 | Enterobacter_cloacae_EcWSU1_uid80739 | 365969793 | YP_004951354.1 | 1 | 1 | 0.605 | 0/18 | 0.233 | Sensitive |
| ATGC001 | Enterobacter_cloacae_SCF1_uid59969 | 311280177 | YP_003942408.1 | 1 | 1 | 0.579 | 0/18 | 0.233 | Sensitive |
| ATGC001 | Enterobacter_cloacae_dissolvens_SDM_uid168997 | 392978365 | YP_006476953.1 | 1 | 1 | 0.605 | 0/18 | 0.233 | Sensitive |
| ATGC001 | Escherichia_coli_042_uid161985 | 387606459 | YP_006095315.1 | 1 | 1 | 0.605 | 0/18 | 0.233 | Sensitive |
| ATGC001 | Escherichia_coli_536_uid58531 | 110641105 | YP_668835.1 | 1 | 1 | 0.605 | 0/18 | 0.233 | Sensitive |
| ATGC001 | Escherichia_coli_55989_uid59383 | 218694381 | YP_002402048.1 | 1 | 1 | 0.605 | 0/18 | 0.233 | Sensitive |
| ATGC001 | Escherichia_coli_ABU_83972_uid161975 | 386638255 | YP_006105053.1 | 1 | 1 | 0.605 | 0/18 | 0.233 | Sensitive |
| ATGC001 | Escherichia_coli_APEC_O1_uid58623 | 117623126 | YP_852039.1 | 1 | 1 | 0.605 | 0/18 | 0.233 | Sensitive |
| ATGC001 | Escherichia_coli_APEC_O78_uid187277 | 443617023 | YP_007380879.1 | 1 | 1 | 0.605 | 0/18 | 0.233 | Sensitive |
| ATGC001 | Escherichia_coli_ATCC_8739_uid58783 | 170020690 | YP_001725644.1 | 1 | 1 | 0.605 | 0/18 | 0.233 | Sensitive |
| ATGC001 | Escherichia_coli_BL21_DE3__uid161947 | 254287830 | YP_003053578.1 | 1 | 1 | 0.605 | 0/18 | 0.233 | Sensitive |

| | | | | | | | | | |
|---|---|---|---|---|---|---|---|---|---|
| ATGC001 | Escherichia_coli_BL21_DE3__uid161949 | 251784450 | YP_002998754.1 | 1 | 1 | 0.605 | 0/18 | 0.233 | Sensitive |
| ATGC001 | Escherichia_coli_BW2952_uid59391 | 238900166 | YP_002925962.1 | 1 | 1 | 0.605 | 0/18 | 0.233 | Sensitive |
| ATGC001 | Escherichia_coli_B_REL606_uid58803 | 254161022 | YP_003044130.1 | 1 | 1 | 0.605 | 0/18 | 0.233 | Sensitive |
| ATGC001 | Escherichia_coli_CFT073_uid57915 | 26246933 | NP_752973.1 | 1 | 1 | 0.605 | 0/18 | 0.233 | Sensitive |
| ATGC001 | Escherichia_coli_DH1_uid161951 | 386596257 | YP_006092657.1 | 1 | 1 | 0.605 | 0/18 | 0.233 | Sensitive |
| ATGC001 | Escherichia_coli_DH1_uid162051 | 387620633 | YP_006128260.1 | 1 | 1 | 0.605 | 0/18 | 0.233 | Sensitive |
| ATGC001 | Escherichia_coli_E24377A_uid58395 | 157155708 | YP_001462126.1 | 1 | 1 | 0.605 | 0/18 | 0.233 | Sensitive |
| ATGC001 | Escherichia_coli_ED1a_uid59379 | 218688748 | YP_002396960.1 | 1 | 1 | 0.605 | 0/18 | 0.233 | Sensitive |
| ATGC001 | Escherichia_coli_ETEC_H10407_uid161993 | 387611448 | YP_006114564.1 | 1 | 1 | 0.605 | 0/18 | 0.233 | Sensitive |
| ATGC001 | Escherichia_coli_HS_uid58393 | 157160430 | YP_001457748.1 | 1 | 1 | 0.605 | 0/18 | 0.233 | Sensitive |
| ATGC001 | Escherichia_coli_IAI1_uid59377 | 218553494 | YP_002386407.1 | 1 | 1 | 0.605 | 0/18 | 0.233 | Sensitive |
| ATGC001 | Escherichia_coli_IAI39_uid59381 | 218700574 | YP_002408203.1 | 1 | 1 | 0.605 | 0/18 | 0.233 | Sensitive |
| ATGC001 | Escherichia_coli_IHE3034_uid162007 | 386598687 | YP_006100193.1 | 1 | 1 | 0.605 | 0/18 | 0.233 | Sensitive |
| ATGC001 | Escherichia_coli_KO11FL_uid162099 | 386702120 | YP_006165957.1 | 1 | 1 | 0.605 | 0/18 | 0.233 | Sensitive |
| ATGC001 | Escherichia_coli_KO11FL_uid52593 | 378713686 | YP_005278579.1 | 1 | 1 | 0.605 | 0/18 | 0.233 | Sensitive |
| ATGC001 | Escherichia_coli_K_12_substr__DH10B_uid58979 | 170080566 | YP_001729886.1 | 1 | 1 | 0.605 | 0/18 | 0.233 | Sensitive |
| ATGC001 | Escherichia_coli_K_12_substr__MDS42_uid193705 | 471332962 | YP_007556439.1 | 1 | 1 | 0.605 | 0/18 | 0.233 | Sensitive |
| ATGC001 | Escherichia_coli_K_12_substr__MG1655_uid57779 | 16128875 | NP_415428.1 | 1 | 1 | 0.605 | 0/18 | 0.233 | Sensitive |
| ATGC001 | Escherichia_coli_K_12_substr__W3110_uid161931 | 388476992 | YP_489180.1 | 1 | 1 | 0.605 | 0/18 | 0.233 | Sensitive |
| ATGC001 | Escherichia_coli_LF82_uid161965 | 222155634 | YP_002555773.1 | 1 | 1 | 0.605 | 0/18 | 0.233 | Sensitive |
| ATGC001 | Escherichia_coli_NA114_uid162139 | 386618457 | YP_006138037.1 | 1 | 1 | 0.605 | 0/18 | 0.233 | Sensitive |

| | | | | | | | | | |
|---|---|---|---|---|---|---|---|---|---|
| ATGC001 | Escherichia_coli_O103_H2_12009_uid41013 | 260843158 | YP_003220936.1 | 1 | 1 | 0.605 | 0/18 | 0.233 | Sensitive |
| ATGC001 | Escherichia_coli_O104_H4_2009EL_2050_uid175905 | 410483439 | YP_006770985.1 | 1 | 1 | 0.605 | 0/18 | 0.233 | Sensitive |
| ATGC001 | Escherichia_coli_O104_H4_2009EL_2071_uid176128 | 407468381 | YP_006785177.1 | 1 | 1 | 0.605 | 0/18 | 0.233 | Sensitive |
| ATGC001 | Escherichia_coli_O104_H4_2011C_3493_uid176127 | 407482887 | YP_006780036.1 | 1 | 1 | 0.605 | 0/18 | 0.233 | Sensitive |
| ATGC001 | Escherichia_coli_O111_H__11128_uid41023 | 260867080 | YP_003233482.1 | 1 | 1 | 0.605 | 0/18 | 0.233 | Sensitive |
| ATGC001 | Escherichia_coli_O127_H6_E2348_69_uid59343 | 215486033 | YP_002328464.1 | 1 | 1 | 0.605 | 0/18 | 0.233 | Sensitive |
| ATGC001 | Escherichia_coli_O157_H7_EC4115_uid59091 | 209400293 | YP_002269580.1 | 1 | 1 | 0.605 | 0/18 | 0.233 | Sensitive |
| ATGC001 | Escherichia_coli_O157_H7_EDL933_uid57831 | 15800769 | NP_286783.1 | 1 | 1 | 0.605 | 0/18 | 0.233 | Sensitive |
| ATGC001 | Escherichia_coli_O157_H7_Sakai_uid57781 | 15830245 | NP_309018.1 | 1 | 1 | 0.605 | 0/18 | 0.233 | Sensitive |
| ATGC001 | Escherichia_coli_O157_H7_TW14359_uid59235 | 254792107 | YP_003076944.1 | 1 | 1 | 0.605 | 0/18 | 0.233 | Sensitive |
| ATGC001 | Escherichia_coli_O26_H11_11368_uid41021 | 260854199 | YP_003228090.1 | 1 | 1 | 0.605 | 0/18 | 0.233 | Sensitive |
| ATGC001 | Escherichia_coli_O55_H7_CB9615_uid46655 | 291281909 | YP_003498727.1 | 1 | 1 | 0.605 | 0/18 | 0.233 | Sensitive |
| ATGC001 | Escherichia_coli_O55_H7_RM12579_uid162153 | 387506019 | YP_006158275.1 | 1 | 1 | 0.605 | 0/18 | 0.233 | Sensitive |
| ATGC001 | Escherichia_coli_O7_K1_CE10_uid162115 | 386623313 | YP_006143041.1 | 1 | 1 | 0.605 | 0/18 | 0.233 | Sensitive |
| ATGC001 | Escherichia_coli_O83_H1_NRG_857C_uid161987 | 387616167 | YP_006119189.1 | 1 | 1 | 0.605 | 0/18 | 0.233 | Sensitive |
| ATGC001 | Escherichia_coli_P12b_uid162061 | 386704088 | YP_006167935.1 | 1 | 1 | 0.605 | 0/18 | 0.233 | Sensitive |
| ATGC001 | Escherichia_coli_S88_uid62979 | 218557813 | YP_002390726.1 | 1 | 1 | 0.605 | 0/18 | 0.233 | Sensitive |
| ATGC001 | Escherichia_coli_SE11_uid59425 | 209918158 | YP_002292242.1 | 1 | 1 | 0.605 | 0/18 | 0.233 | Sensitive |
| ATGC001 | Escherichia_coli_SE15_uid161939 | 387828882 | YP_003348819.1 | 1 | 1 | 0.605 | 0/18 | 0.233 | Sensitive |
| ATGC001 | Escherichia_coli_SMS_3_5_uid58919 | 170683917 | YP_001744263.1 | 1 | 1 | 0.605 | 0/18 | 0.233 | Sensitive |
| ATGC001 | Escherichia_coli_UM146_uid162043 | 386605181 | YP_006111481.1 | 1 | 1 | 0.605 | 0/18 | 0.233 | Sensitive |

| | | | | | | | | | |
|---|---|---|---|---|---|---|---|---|---|
| ATGC001 | Escherichia_coli_UMN026_uid62981 | 218704335 | YP_002411854.1 | 1 | 1 | 0.605 | 0/18 | 0.233 | Sensitive |
| ATGC001 | Escherichia_coli_UMNK88_uid161991 | 386613182 | YP_006132848.1 | 1 | 1 | 0.605 | 0/18 | 0.233 | Sensitive |
| ATGC001 | Escherichia_coli_UTI89_uid58541 | 91210009 | YP_539995.1 | 1 | 1 | 0.605 | 0/18 | 0.233 | Sensitive |
| ATGC001 | Escherichia_coli_W_uid162011 | 386608276 | YP_006123762.1 | 1 | 1 | 0.605 | 0/18 | 0.233 | Sensitive |
| ATGC001 | Escherichia_coli_W_uid162101 | 386708719 | YP_006172440.1 | 1 | 1 | 0.605 | 0/18 | 0.233 | Sensitive |
| ATGC001 | Escherichia_coli_Xuzhou21_uid163995 | 387881520 | YP_006311822.1 | 1 | 1 | 0.605 | 0/18 | 0.233 | Sensitive |
| ATGC001 | Escherichia_coli__BL21_Gold_DE3_pLysS_AG__uid59245 | 253774063 | YP_003036894.1 | 1 | 1 | 0.605 | 0/18 | 0.233 | Sensitive |
| ATGC001 | Escherichia_coli__clone_D_i14__uid162049 | 386633367 | YP_006153086.1 | 1 | 1 | 0.605 | 0/18 | 0.233 | Sensitive |
| ATGC001 | Escherichia_coli__clone_D_i2__uid162047 | 386628447 | YP_006148167.1 | 1 | 1 | 0.605 | 0/18 | 0.233 | Sensitive |
| ATGC001 | Escherichia_fergusonii_ATCC_35469_uid59375 | 218548425 | YP_002382216.1 | 1 | 1 | 0.605 | 0/18 | 0.233 | Sensitive |
| ATGC001 | Salmonella_bongori_NCTC_12419_uid70155 | 339998835 | YP_004729718.1 | 1 | 1 | 0.579 | 0/18 | 0.233 | Sensitive |
| ATGC001 | Salmonella_enterica_arizonae_serovar_62_z4_z23__uid58191 | 161503895 | YP_001571007.1 | 1 | 1 | 0.579 | 0/18 | 0.233 | Sensitive |
| ATGC001 | Salmonella_enterica_serovar_Agona_SL483_uid59431 | 197249917 | YP_002145896.1 | 1 | 1 | 0.579 | 0/18 | 0.233 | Sensitive |
| ATGC001 | Salmonella_enterica_serovar_Choleraesuis_SC_B67_uid58017 | 62179502 | YP_215919.1 | 1 | 1 | 0.579 | 0/18 | 0.233 | Sensitive |
| ATGC001 | Salmonella_enterica_serovar_Dublin_CT_02021853_uid58917 | 198246031 | YP_002214902.1 | 1 | 1 | 0.579 | 0/18 | 0.233 | Sensitive |
| ATGC001 | Salmonella_enterica_serovar_Enteritidis_P125109_uid59247 | 207856369 | YP_002243020.1 | 1 | 1 | 0.579 | 0/18 | 0.233 | Sensitive |
| ATGC001 | Salmonella_enterica_serovar_Gallinarum_287_91_uid59249 | 205352185 | YP_002225986.1 | 1 | 1 | 0.579 | 0/18 | 0.233 | Sensitive |
| ATGC001 | Salmonella_enterica_serovar_Gallinarum_pullorum_RKS5078_uid87035 | 378955698 | YP_005213185.1 | 1 | 1 | 0.579 | 0/18 | 0.233 | Sensitive |
| ATGC001 | Salmonella_enterica_serovar_Heidelberg_B182_uid162195 | 386590843 | YP_006087243.1 | 1 | 1 | 0.579 | 0/18 | 0.233 | Sensitive |
| ATGC001 | Salmonella_enterica_serovar_Heidelberg_SL476_uid58973 | 194449431 | YP_002044970.1 | 1 | 1 | 0.579 | 0/18 | 0.233 | Sensitive |
| ATGC001 | Salmonella_enterica_serovar_Javiana_CFSAN001992_uid190101 | 452120859 | YP_007471107.1 | 1 | 1 | 0.579 | 0/18 | 0.233 | Sensitive |

| | | | | | | | | | |
|---|---|---|---|---|---|---|---|---|---|
| ATGC001 | Salmonella_enterica_serovar_Newport_SL254_uid58831 | 194446151 | YP_002040176.1 | 1 | 1 | 0.579 | 0/18 | 0.233 | Sensitive |
| ATGC001 | Salmonella_enterica_serovar_Paratyphi_A_AKU_12601_uid59269 | 197362895 | YP_002142532.1 | 1 | 1 | 0.579 | 0/18 | 0.233 | Sensitive |
| ATGC001 | Salmonella_enterica_serovar_Paratyphi_A_ATCC_9150_uid58201 | 56413972 | YP_151047.1 | 1 | 1 | 0.579 | 0/18 | 0.233 | Sensitive |
| ATGC001 | Salmonella_enterica_serovar_Paratyphi_B_SPB7_uid59097 | 161614786 | YP_001588751.1 | 1 | 1 | 0.579 | 0/18 | 0.233 | Sensitive |
| ATGC001 | Salmonella_enterica_serovar_Paratyphi_C_RKS4594_uid59063 | 224582787 | YP_002636585.1 | 1 | 1 | 0.579 | 0/18 | 0.233 | Sensitive |
| ATGC001 | Salmonella_enterica_serovar_Schwarzengrund_CVM19633_uid58915 | 194737397 | YP_002114029.1 | 1 | 1 | 0.579 | 0/18 | 0.233 | Sensitive |
| ATGC001 | Salmonella_enterica_serovar_Typhi_CT18_uid57793 | 16759848 | NP_455465.1 | 1 | 1 | 0.579 | 0/18 | 0.233 | Sensitive |
| ATGC001 | Salmonella_enterica_serovar_Typhi_P_stx_12_uid87001 | 378960121 | YP_005217607.1 | 1 | 1 | 0.579 | 0/18 | 0.233 | Sensitive |
| ATGC001 | Salmonella_enterica_serovar_Typhi_Ty21a_uid201427 | 488654470 | YP_007926273.1 | 1 | 1 | 0.579 | 0/18 | 0.233 | Sensitive |
| ATGC001 | Salmonella_enterica_serovar_Typhi_Ty2_uid57973 | 29142379 | NP_805721.1 | 1 | 1 | 0.579 | 0/18 | 0.233 | Sensitive |
| ATGC001 | Salmonella_enterica_serovar_Typhimurium_14028S_uid86059 | 378449337 | YP_005236696.1 | 1 | 1 | 0.579 | 0/18 | 0.233 | Sensitive |
| ATGC001 | Salmonella_enterica_serovar_Typhimurium_798_uid158047 | 383495717 | YP_005396406.1 | 1 | 1 | 0.579 | 0/18 | 0.233 | Sensitive |
| ATGC001 | Salmonella_enterica_serovar_Typhimurium_LT2_uid57799 | 16764338 | NP_459953.1 | 1 | 1 | 0.579 | 0/18 | 0.233 | Sensitive |
| ATGC001 | Salmonella_enterica_serovar_Typhimurium_SL1344_uid86645 | 378698873 | YP_005180830.1 | 1 | 1 | 0.579 | 0/18 | 0.233 | Sensitive |
| ATGC001 | Salmonella_enterica_serovar_Typhimurium_ST4_74_uid84393 | 379700144 | YP_005241872.1 | 1 | 1 | 0.579 | 0/18 | 0.233 | Sensitive |
| ATGC001 | Salmonella_enterica_serovar_Typhimurium_T000240_uid84397 | 378983536 | YP_005246691.1 | 1 | 1 | 0.579 | 0/18 | 0.233 | Sensitive |
| ATGC001 | Salmonella_enterica_serovar_Typhimurium_U288_uid198746 | 482904482 | YP_007903042.1 | 1 | 1 | 0.579 | 0/18 | 0.233 | Sensitive |
| ATGC001 | Salmonella_enterica_serovar_Typhimurium_UK_1_uid87049 | 378988322 | YP_005251486.1 | 1 | 1 | 0.579 | 0/18 | 0.233 | Sensitive |
| ATGC001 | Salmonella_enterica_serovar_Typhimurium_uid86061 | 378444415 | YP_005232047.1 | 1 | 1 | 0.579 | 0/18 | 0.233 | Sensitive |
| ATGC001 | Shigella_boydii_CDC_3083_94_uid58415 | 187733570 | YP_001880897.1 | 1 | 1 | 0.605 | 0/18 | 0.233 | Sensitive |
| ATGC001 | Shigella_boydii_Sb227_uid58215 | 82544646 | YP_408593.1 | 1 | 1 | 0.605 | 0/18 | 0.233 | Sensitive |

| | | | | | | | | | |
|---|---|---|---|---|---|---|---|---|---|
| ATGC001 | Shigella_dysenteriae_Sd197_uid58213 | 82777572 | YP_403921.1 | 1 | 1 | 0.605 | 0/18 | 0.233 | Sensitive |
| ATGC001 | Shigella_flexneri_2002017_uid159233 | 384542482 | YP_005726544.1 | 1 | 1 | 0.605 | 0/18 | 0.233 | Sensitive |
| ATGC001 | Shigella_flexneri_2a_2457T_uid57991 | 30062442 | NP_836613.1 | 1 | 1 | 0.605 | 0/18 | 0.233 | Sensitive |
| ATGC001 | Shigella_flexneri_2a_301_uid62907 | 24112316 | NP_706826.1 | 1 | 1 | 0.605 | 0/18 | 0.233 | Sensitive |
| ATGC001 | Shigella_flexneri_5_8401_uid58583 | 110804915 | YP_688435.1 | 1 | 1 | 0.605 | 0/18 | 0.233 | Sensitive |
| ATGC001 | Shigella_sonnei_53G_uid84383 | 383177549 | YP_005455554.1 | 1 | 1 | 0.605 | 0/18 | 0.233 | Sensitive |
| ATGC001 | Shigella_sonnei_Ss046_uid58217 | 74311464 | YP_309883.1 | 1 | 1 | 0.605 | 0/18 | 0.233 | Sensitive |
| ATGC002 | Enterobacter_aerogenes_EA1509E_uid187411 | 444352328 | YP_007388472.1 | 1 | 1 | 0.605 | 0/18 | 0.233 | Sensitive |
| ATGC002 | Enterobacter_aerogenes_KCTC_2190_uid68103 | 336249511 | YP_004593221.1 | 1 | 1 | 0.605 | 0/18 | 0.233 | Sensitive |
| ATGC002 | Klebsiella_oxytoca_E718_uid170256 | 397657065 | YP_006497767.1 | 0.84 | 0.833 | 0.526 | 0/18 | 0.133 | Unclassified |
| ATGC002 | Klebsiella_oxytoca_KCTC_1686_uid83159 | 375259991 | YP_005019161.1 | 1 | 1 | 0.605 | 0/18 | 0.233 | Sensitive |
| ATGC002 | Klebsiella_pneumoniae_1084_uid174151 | 402781548 | YP_006637094.1 | 1 | 1 | 0.579 | 0/18 | 0.233 | Sensitive |
| ATGC002 | Klebsiella_pneumoniae_342_uid59145 | 206580881 | YP_002239442.1 | 1 | 1 | 0.579 | 0/18 | 0.233 | Sensitive |
| ATGC002 | Klebsiella_pneumoniae_HS11286_uid84387 | 378977975 | YP_005226116.1 | 1 | 1 | 0.579 | 0/18 | 0.233 | Sensitive |
| ATGC002 | Klebsiella_pneumoniae_KCTC_2242_uid162147 | 386034120 | YP_005954033.1 | 1 | 1 | 0.579 | 0/18 | 0.233 | Sensitive |
| ATGC002 | Klebsiella_pneumoniae_MGH_78578_uid57619 | 152969493 | YP_001334602.1 | 1 | 1 | 0.579 | 0/18 | 0.233 | Sensitive |
| ATGC002 | Klebsiella_pneumoniae_NTUH_K2044_uid59073 | 238893965 | YP_002918699.1 | 1 | 1 | 0.579 | 0/18 | 0.233 | Sensitive |
| ATGC002 | Klebsiella_variicola_At_22_uid42113 | 288936293 | YP_003440352.1 | 1 | 1 | 0.579 | 0/18 | 0.233 | Sensitive |
| ATGC003 | Streptococcus_mitis_B6_uid46097 | 289167609 | YP_003445878.1 | 0.76 | 0.833 | 1 | 0/18 | 0.133 | Resistant |
| ATGC003 | Streptococcus_pneumoniae_670_6B_uid52533 | 307127059 | YP_003879090.1 | 0.76 | 0.833 | 1 | 0/18 | 0.133 | Resistant |
| ATGC003 | Streptococcus_pneumoniae_70585_uid59125 | 225859138 | YP_002740648.1 | 0.76 | 0.833 | 1 | 0/18 | 0.133 | Resistant |

| | | | | | | | | | |
|---|---|---|---|---|---|---|---|---|---|
| ATGC003 | Streptococcus_pneumoniae_AP200_uid52453 | 307068023 | YP_003876989.1 | 0.8 | 0.889 | 1 | 0/18 | 0.133 | Resistant |
| ATGC003 | Streptococcus_pneumoniae_ATCC_700669_uid59287 | 221232113 | YP_002511266.1 | 0.76 | 0.833 | 1 | 0/18 | 0.133 | Resistant |
| ATGC003 | Streptococcus_pneumoniae_CGSP14_uid59181 | 182684331 | YP_001836078.1 | 0.8 | 0.889 | 1 | 0/18 | 0.133 | Resistant |
| ATGC003 | Streptococcus_pneumoniae_D39_uid58581 | 116515889 | YP_816672.1 | 0.76 | 0.833 | 1 | 0/18 | 0.133 | Resistant |
| ATGC003 | Streptococcus_pneumoniae_G54_uid59167 | 194397424 | YP_002038019.1 | 0.8 | 0.889 | 1 | 0/18 | 0.133 | Resistant |
| ATGC003 | Streptococcus_pneumoniae_Hungary19A_6_uid59117 | 169833234 | YP_001694807.1 | 0.76 | 0.833 | 1 | 0/18 | 0.133 | Resistant |
| ATGC003 | Streptococcus_pneumoniae_INV104_uid162039 | 387626630 | YP_006062805.1 | 0.8 | 0.889 | 1 | 0/18 | 0.133 | Resistant |
| ATGC003 | Streptococcus_pneumoniae_INV200_uid162035 | 387759525 | YP_006066503.1 | 0.8 | 0.889 | 1 | 0/18 | 0.133 | Resistant |
| ATGC003 | Streptococcus_pneumoniae_JJA_uid59121 | 225854819 | YP_002736331.1 | 0.76 | 0.833 | 1 | 0/18 | 0.133 | Resistant |
| ATGC003 | Streptococcus_pneumoniae_OXC141_uid162037 | 387757652 | YP_006064631.1 | 0.8 | 0.889 | 1 | 0/18 | 0.133 | Resistant |
| ATGC003 | Streptococcus_pneumoniae_P1031_uid59123 | 225857007 | YP_002738518.1 | 0.76 | 0.833 | 1 | 0/18 | 0.133 | Resistant |
| ATGC003 | Streptococcus_pneumoniae_R6_uid57859 | 15903272 | NP_358822.1 | 0.76 | 0.833 | 1 | 0/18 | 0.133 | Resistant |
| ATGC003 | Streptococcus_pneumoniae_SPNA45_uid174986 | 405760692 | YP_006701288.1 | 0.8 | 0.889 | 1 | 0/18 | 0.133 | Resistant |
| ATGC003 | Streptococcus_pneumoniae_ST556_uid162191 | 387788045 | YP_006253113.1 | 0.76 | 0.833 | 1 | 0/18 | 0.133 | Resistant |
| ATGC003 | Streptococcus_pneumoniae_TCH8431_19A_uid49735 | 298502679 | YP_003724619.1 | 0.76 | 0.833 | 1 | 0/18 | 0.133 | Resistant |
| ATGC003 | Streptococcus_pneumoniae_TIGR4_uid57857 | 15901225 | NP_345829.1 | 0.8 | 0.889 | 1 | 0/18 | 0.133 | Resistant |
| ATGC003 | Streptococcus_pneumoniae_Taiwan19F_14_uid59119 | 225860847 | YP_002742356.1 | 0.76 | 0.833 | 1 | 0/18 | 0.133 | Resistant |
| ATGC003 | Streptococcus_pneumoniae_gamPNI0373_uid175861 | 410476756 | YP_006743515.1 | 0.76 | 0.833 | 1 | 0/18 | 0.133 | Resistant |
| ATGC003 | Streptococcus_pseudopneumoniae_IS7493_uid71153 | 342164003 | YP_004768642.1 | 0.76 | 0.833 | 1 | 0/18 | 0.133 | Resistant |
| ATGC004 | Streptococcus_dysgalactiae_equisimilis_AC_2713_uid178644 | 410495137 | YP_006904983.1 | 0.76 | 0.778 | 1 | 0/18 | 0.2 | Resistant |
| ATGC004 | Streptococcus_dysgalactiae_equisimilis_ATCC_12394_uid161979 | 386317316 | YP_006013480.1 | 0.76 | 0.778 | 1 | 0/18 | 0.2 | Resistant |

| | | | | | | | | | |
|---|---|---|---|---|---|---|---|---|---|
| ATGC004 | Streptococcus_dysgalactiae_equisimilis_GGS_124_uid59103 | 251782759 | YP_002997062.1 | 0.76 | 0.778 | 1 | 0/18 | 0.2 | Resistant |
| ATGC004 | Streptococcus_dysgalactiae_equisimilis_RE378_uid176684 | 408401937 | YP_006859901.1 | 0.76 | 0.778 | 1 | 0/18 | 0.2 | Resistant |
| ATGC004 | Streptococcus_pyogenes_A20_uid178106 | 410680768 | YP_006933170.1 | 0.76 | 0.778 | 1 | 0/18 | 0.2 | Resistant |
| ATGC004 | Streptococcus_pyogenes_Alab49_uid162171 | 386362919 | YP_006072250.1 | 0.76 | 0.778 | 1 | 0/18 | 0.2 | Resistant |
| ATGC004 | Streptococcus_pyogenes_M1_476_uid193766 | 470201258 | YP_007587687.1 | 0.76 | 0.778 | 1 | 0/18 | 0.2 | Resistant |
| ATGC004 | Streptococcus_pyogenes_M1_GAS_uid57845 | 15675285 | NP_269459.1 | 0.76 | 0.778 | 1 | 0/18 | 0.2 | Resistant |
| ATGC004 | Streptococcus_pyogenes_MGAS10270_uid58571 | 94990667 | YP_598767.1 | 0.76 | 0.778 | 1 | 0/18 | 0.2 | Resistant |
| ATGC004 | Streptococcus_pyogenes_MGAS10394_uid58105 | 50914419 | YP_060391.1 | 0.76 | 0.778 | 1 | 0/18 | 0.2 | Resistant |
| ATGC004 | Streptococcus_pyogenes_MGAS10750_uid58575 | 94994596 | YP_602694.1 | 0.76 | 0.778 | 1 | 0/18 | 0.2 | Resistant |
| ATGC004 | Streptococcus_pyogenes_MGAS15252_uid158037 | 383480183 | YP_005389077.1 | 0.76 | 0.778 | 1 | 0/18 | 0.2 | Resistant |
| ATGC004 | Streptococcus_pyogenes_MGAS1882_uid158061 | 383494100 | YP_005411776.1 | 0.76 | 0.778 | 1 | 0/18 | 0.2 | Resistant |
| ATGC004 | Streptococcus_pyogenes_MGAS2096_uid58573 | 94992660 | YP_600759.1 | 0.76 | 0.778 | 1 | 0/18 | 0.2 | Resistant |
| ATGC004 | Streptococcus_pyogenes_MGAS315_uid57911 | 21910563 | NP_664831.1 | 0.76 | 0.778 | 1 | 0/18 | 0.2 | Resistant |
| ATGC004 | Streptococcus_pyogenes_MGAS5005_uid58337 | 71910914 | YP_282464.1 | 0.76 | 0.778 | 1 | 0/18 | 0.2 | Resistant |
| ATGC004 | Streptococcus_pyogenes_MGAS6180_uid58335 | 71903758 | YP_280561.1 | 0.76 | 0.778 | 1 | 0/18 | 0.2 | Resistant |
| ATGC004 | Streptococcus_pyogenes_MGAS8232_uid57871 | 19746325 | NP_607461.1 | 0.76 | 0.778 | 1 | 0/18 | 0.2 | Resistant |
| ATGC004 | Streptococcus_pyogenes_MGAS9429_uid58569 | 94988775 | YP_596876.1 | 0.76 | 0.778 | 1 | 0/18 | 0.2 | Resistant |
| ATGC004 | Streptococcus_pyogenes_Manfredo_uid57847 | 139473606 | YP_001128322.1 | 0.76 | 0.778 | 1 | 0/18 | 0.2 | Resistant |
| ATGC004 | Streptococcus_pyogenes_NZ131_uid59035 | 209559592 | YP_002286064.1 | 0.76 | 0.778 | 1 | 0/18 | 0.2 | Resistant |
| ATGC004 | Streptococcus_pyogenes_SSI_1_uid57895 | 28895745 | NP_802095.1 | 0.76 | 0.778 | 1 | 0/18 | 0.2 | Resistant |
| ATGC005 | Streptococcus_suis_05ZYH33_uid58663 | 146318250 | YP_001197962.1 | 0.28 | 0.167 | 0.237 | 0/18 | 0.033 | Unclassified |

| | | | | | | | | | |
|---|---|---|---|---|---|---|---|---|---|
| ATGC005 | Streptococcus_suis_05ZYH33_uid58663 | 146318251 | YP_001197963.1 | 0.2 | 0.222 | 0.368 | 0/18 | 0 | Unclassified |
| ATGC005 | Streptococcus_suis_05ZYH33_uid58663 | 146318252 | YP_001197964.1 | 0.28 | 0.333 | 0.263 | 0/18 | 0 | Unclassified |
| ATGC005 | Streptococcus_suis_98HAH33_uid58665 | 146320447 | YP_001200158.1 | 0.72 | 0.778 | 0.974 | 0/18 | 0.133 | Unclassified |
| ATGC005 | Streptococcus_suis_A7_uid162111 | 386587826 | YP_006084227.1 | 0.76 | 0.833 | 1 | 0/18 | 0.133 | Resistant |
| ATGC005 | Streptococcus_suis_BM407_uid59321 | 253755847 | YP_003028987.1 | 0.76 | 0.833 | 1 | 0/18 | 0.133 | Resistant |
| ATGC005 | Streptococcus_suis_D12_uid162127 | 386586487 | YP_006082889.1 | 0.76 | 0.833 | 1 | 0/18 | 0.133 | Resistant |
| ATGC005 | Streptococcus_suis_D9_uid162125 | 386583831 | YP_006080234.1 | 0.76 | 0.833 | 1 | 0/18 | 0.133 | Resistant |
| ATGC005 | Streptococcus_suis_D9_uid162125 | 386583834 | YP_006080237.1 | 0.76 | 0.833 | 1 | 0/18 | 0.133 | Resistant |
| ATGC005 | Streptococcus_suis_GZ1_uid161937 | 386577587 | YP_006073993.1 | 0.76 | 0.833 | 1 | 0/18 | 0.133 | Resistant |
| ATGC005 | Streptococcus_suis_JS14_uid162095 | 386579524 | YP_006075929.1 | 0.76 | 0.833 | 1 | 0/18 | 0.133 | Resistant |
| ATGC005 | Streptococcus_suis_P1_7_uid32235 | 253753323 | YP_003026464.1 | 0.76 | 0.833 | 1 | 0/18 | 0.133 | Resistant |
| ATGC005 | Streptococcus_suis_S735_uid174333 | 403061227 | YP_006649443.1 | 0.76 | 0.833 | 1 | 0/18 | 0.133 | Resistant |
| ATGC005 | Streptococcus_suis_SC070731_uid193769 | 476419516 | YP_007570512.1 | 0.76 | 0.833 | 1 | 0/18 | 0.133 | Resistant |
| ATGC005 | Streptococcus_suis_SC84_uid59323 | 253751422 | YP_003024563.1 | 0.76 | 0.833 | 1 | 0/18 | 0.133 | Resistant |
| ATGC005 | Streptococcus_suis_SS12_uid162123 | 386581593 | YP_006077997.1 | 0.76 | 0.833 | 1 | 0/18 | 0.133 | Resistant |
| ATGC005 | Streptococcus_suis_ST1_uid167482 | 389856845 | YP_006359088.1 | 0.76 | 0.833 | 1 | 0/18 | 0.133 | Resistant |
| ATGC005 | Streptococcus_suis_ST3_uid66327 | 330832574 | YP_004401399.1 | 0.76 | 0.833 | 1 | 0/18 | 0.133 | Resistant |
| ATGC005 | Streptococcus_suis_TL13_uid203123 | 499143469 | YP_007975533.1 | 0.76 | 0.833 | 1 | 0/18 | 0.133 | Resistant |
| ATGC014 | Bacillus_anthracis_A0248_uid59385 | 229603924 | YP_002867201.1 | 0.8 | 0.889 | 1 | 0/18 | 0.2 | Resistant |
| ATGC014 | Bacillus_anthracis_Ames_uid57909 | 30262911 | NP_845288.1 | 0.8 | 0.889 | 1 | 0/18 | 0.2 | Resistant |
| ATGC014 | Bacillus_anthracis_CDC_684_uid59303 | 227814241 | YP_002814250.1 | 0.8 | 0.889 | 1 | 0/18 | 0.2 | Resistant |

| | | | | | | | | | |
|---|---|---|---|---|---|---|---|---|---|
| ATGC014 | Bacillus_anthracis_H9401_uid162021 | 386736690 | YP_006209871.1 | 0.8 | 0.889 | 1 | 0/18 | 0.2 | Resistant |
| ATGC014 | Bacillus_anthracis_Sterne_uid58091 | 49185750 | YP_029002.1 | 0.8 | 0.889 | 1 | 0/18 | 0.2 | Resistant |
| ATGC014 | Bacillus_anthracis__Ames_Ancestor__uid58083 | 47778099 | YP_019596.2 | 0.8 | 0.889 | 1 | 0/18 | 0.2 | Resistant |
| ATGC014 | Bacillus_cereus_03BB102_uid59299 | 225864918 | YP_002750296.1 | 0.8 | 0.889 | 1 | 0/18 | 0.2 | Resistant |
| ATGC014 | Bacillus_cereus_AH187_uid58753 | 217960388 | YP_002338950.1 | 0.8 | 0.889 | 1 | 0/18 | 0.2 | Resistant |
| ATGC014 | Bacillus_cereus_AH820_uid58751 | 218904069 | YP_002451903.1 | 0.8 | 0.889 | 1 | 0/18 | 0.2 | Resistant |
| ATGC014 | Bacillus_cereus_ATCC_10987_uid57673 | 42782050 | NP_979297.1 | 0.8 | 0.889 | 1 | 0/18 | 0.2 | Resistant |
| ATGC014 | Bacillus_cereus_ATCC_14579_uid57975 | 30021054 | NP_832685.1 | 0.8 | 0.889 | 1 | 0/18 | 0.2 | Resistant |
| ATGC014 | Bacillus_cereus_B4264_uid58757 | 218234153 | YP_002367664.1 | 0.8 | 0.889 | 1 | 0/18 | 0.2 | Resistant |
| ATGC014 | Bacillus_cereus_E33L_uid58103 | 52142569 | YP_084260.1 | 0.8 | 0.889 | 1 | 0/18 | 0.2 | Resistant |
| ATGC014 | Bacillus_cereus_F837_76_uid83611 | 376266805 | YP_005119517.1 | 0.8 | 0.889 | 1 | 0/18 | 0.2 | Resistant |
| ATGC014 | Bacillus_cereus_FRI_35_uid173403 | 402556845 | YP_006598116.1 | 0.8 | 0.889 | 1 | 0/18 | 0.2 | Resistant |
| ATGC014 | Bacillus_cereus_G9842_uid58759 | 218898022 | YP_002446433.1 | 0.8 | 0.889 | 1 | 0/18 | 0.2 | Resistant |
| ATGC014 | Bacillus_cereus_NC7401_uid82815 | 375284901 | YP_005105340.1 | 0.8 | 0.889 | 1 | 0/18 | 0.2 | Resistant |
| ATGC014 | Bacillus_cereus_Q1_uid58529 | 222096443 | YP_002530500.1 | 0.68 | 0.778 | 0.868 | 0/18 | 0.133 | Unclassified |
| ATGC014 | Bacillus_cereus_biovar_anthracis_CI_uid50615 | 301054461 | YP_003792672.1 | 0.8 | 0.889 | 1 | 0/18 | 0.2 | Resistant |
| ATGC014 | Bacillus_thuringiensis_Al_Hakam_uid58795 | 118478280 | YP_895431.1 | 0.8 | 0.889 | 1 | 0/18 | 0.2 | Resistant |
| ATGC014 | Bacillus_thuringiensis_BMB171_uid49135 | 296503477 | YP_003665177.1 | 0.8 | 0.889 | 1 | 0/18 | 0.2 | Resistant |
| ATGC014 | Bacillus_thuringiensis_Bt407_uid177931 | 410675318 | YP_006927689.1 | 0.8 | 0.889 | 1 | 0/18 | 0.2 | Resistant |
| ATGC014 | Bacillus_thuringiensis_HD_771_uid173374 | 402559734 | YP_006602458.1 | 0.8 | 0.889 | 1 | 0/18 | 0.2 | Resistant |
| ATGC014 | Bacillus_thuringiensis_HD_789_uid173860 | 434375919 | YP_006610563.1 | 0.8 | 0.889 | 1 | 0/18 | 0.2 | Resistant |

| | | | | | | | | | |
|---|---|---|---|---|---|---|---|---|---|
| ATGC014 | Bacillus_thuringiensis_MC28_uid176369 | 407705375 | YP_006828960.1 | 0.8 | 0.889 | 1 | 0/18 | 0.2 | Resistant |
| ATGC014 | Bacillus_thuringiensis_serovar_IS5056_uid190186 | 452199370 | YP_007479451.1 | 0.8 | 0.889 | 1 | 0/18 | 0.2 | Resistant |
| ATGC014 | Bacillus_thuringiensis_serovar_chinensis_CT_43_uid158151 | 384187010 | YP_005572906.1 | 0.8 | 0.889 | 1 | 0/18 | 0.2 | Resistant |
| ATGC014 | Bacillus_thuringiensis_serovar_finitimus_YBT_020_uid158875 | 384180831 | YP_005566593.1 | 0.8 | 0.889 | 1 | 0/18 | 0.2 | Resistant |
| ATGC014 | Bacillus_thuringiensis_serovar_konkukian_97_27_uid58089 | 49479054 | YP_037018.1 | 0.8 | 0.889 | 1 | 0/18 | 0.2 | Resistant |
| ATGC014 | Bacillus_thuringiensis_serovar_kurstaki_HD73_uid189188 | 449089689 | YP_007422130.1 | 0.8 | 0.889 | 1 | 0/18 | 0.2 | Resistant |
| ATGC014 | Bacillus_weihenstephanensis_KBAB4_uid58315 | 163939434 | YP_001644318.1 | 0.6 | 0.722 | 0.947 | 0/18 | 0.2 | Unclassified |
| ATGC014 | Bacillus_weihenstephanensis_KBAB4_uid58315 | 163940691 | YP_001645575.1 | 0.8 | 0.889 | 1 | 0/18 | 0.2 | Resistant |
| ATGC015 | Bacillus_JS_uid162189 | 386758839 | YP_006232055.1 | 0.8 | 0.833 | 1 | 0/18 | 0.167 | Resistant |
| ATGC015 | Bacillus_amyloliquefaciens_DSM_7_uid53535 | 308174052 | YP_003920757.1 | 0.8 | 0.833 | 1 | 0/18 | 0.167 | Resistant |
| ATGC015 | Bacillus_amyloliquefaciens_FZB42_uid58271 | 154686508 | YP_001421669.1 | 0.8 | 0.833 | 1 | 0/18 | 0.167 | Resistant |
| ATGC015 | Bacillus_amyloliquefaciens_IT_45_uid181617 | 451346555 | YP_007445186.1 | 0.8 | 0.833 | 1 | 0/18 | 0.167 | Resistant |
| ATGC015 | Bacillus_amyloliquefaciens_LL3_uid158133 | 384164827 | YP_005546206.1 | 0.8 | 0.833 | 1 | 0/18 | 0.167 | Resistant |
| ATGC015 | Bacillus_amyloliquefaciens_TA208_uid158701 | 384158734 | YP_005540807.1 | 0.8 | 0.833 | 1 | 0/18 | 0.167 | Resistant |
| ATGC015 | Bacillus_amyloliquefaciens_XH7_uid158881 | 384167795 | YP_005549173.1 | 0.8 | 0.833 | 1 | 0/18 | 0.167 | Resistant |
| ATGC015 | Bacillus_amyloliquefaciens_Y2_uid165195 | 387898864 | YP_006329160.1 | 0.8 | 0.833 | 1 | 0/18 | 0.167 | Resistant |
| ATGC015 | Bacillus_amyloliquefaciens_plantarum_AS43_3_uid183682 | 429505647 | YP_007186831.1 | 0.8 | 0.833 | 1 | 0/18 | 0.167 | Resistant |
| ATGC015 | Bacillus_amyloliquefaciens_plantarum_CAU_B946_uid84215 | 375362774 | YP_005130813.1 | 0.8 | 0.833 | 1 | 0/18 | 0.167 | Resistant |
| ATGC015 | Bacillus_amyloliquefaciens_plantarum_UCMB5036_uid190705 | 452856020 | YP_007497703.1 | 0.8 | 0.833 | 1 | 0/18 | 0.167 | Resistant |
| ATGC015 | Bacillus_amyloliquefaciens_plantarum_YAU_B9601_Y2_uid159001 | 384265857 | YP_005421564.1 | 0.8 | 0.833 | 1 | 0/18 | 0.167 | Resistant |
| ATGC015 | Bacillus_atrophaeus_1942_uid59887 | 311068775 | YP_003973698.1 | 0.8 | 0.833 | 1 | 0/18 | 0.2 | Resistant |

| | | | | | | | | | |
|---|---|---|---|---|---|---|---|---|---|
| ATGC015 | Bacillus_subtilis_168_uid57675 | 16079317 | NP_390141.1 | 0.8 | 0.833 | 1 | 0/18 | 0.167 | Resistant |
| ATGC015 | Bacillus_subtilis_6051_HGW_uid193706 | 470162463 | YP_007534238.1 | 0.8 | 0.833 | 1 | 0/18 | 0.167 | Resistant |
| ATGC015 | Bacillus_subtilis_BAB_1_uid195461 | 472330637 | YP_007662913.1 | 0.8 | 0.833 | 1 | 0/18 | 0.167 | Resistant |
| ATGC015 | Bacillus_subtilis_BSP1_uid184010 | 430758441 | YP_007209205.1 | 0.8 | 0.833 | 1 | 0/18 | 0.167 | Resistant |
| ATGC015 | Bacillus_subtilis_BSn5_uid62463 | 321311729 | YP_004204016.1 | 0.8 | 0.833 | 1 | 0/18 | 0.167 | Resistant |
| ATGC015 | Bacillus_subtilis_QB928_uid173926 | 402776518 | YP_006630462.1 | 0.8 | 0.833 | 1 | 0/18 | 0.167 | Resistant |
| ATGC015 | Bacillus_subtilis_RO_NN_1_uid158879 | 384175867 | YP_005557252.1 | 0.8 | 0.833 | 1 | 0/18 | 0.167 | Resistant |
| ATGC015 | Bacillus_subtilis_XF_1_uid189187 | 449094755 | YP_007427246.1 | 0.8 | 0.833 | 1 | 0/18 | 0.167 | Resistant |
| ATGC015 | Bacillus_subtilis_natto_BEST195_uid183001 | 428279726 | YP_005561461.1 | 0.8 | 0.833 | 1 | 0/18 | 0.167 | Resistant |
| ATGC015 | Bacillus_subtilis_spizizenii_TU_B_10_uid73967 | 350266432 | YP_004877739.1 | 0.8 | 0.833 | 1 | 0/18 | 0.167 | Resistant |
| ATGC015 | Bacillus_subtilis_spizizenii_W23_uid51879 | 305674894 | YP_003866566.1 | 0.8 | 0.833 | 1 | 0/18 | 0.167 | Resistant |
| ATGC021 | Chlamydia_trachomatis_434_Bu_uid61633 | 166154577 | YP_001654695.1 | 0.92 | 1 | 0.605 | 0/18 | 0.267 | Sensitive |
| ATGC021 | Chlamydia_trachomatis_A2497_uid159863 | 376282372 | YP_005156198.1 | 0.92 | 1 | 0.605 | 0/18 | 0.267 | Sensitive |
| ATGC021 | Chlamydia_trachomatis_A2497_uid159993 | 385270053 | YP_005813213.1 | 0.92 | 1 | 0.605 | 0/18 | 0.267 | Sensitive |
| ATGC021 | Chlamydia_trachomatis_A_363_uid196769 | 478458919 | YP_007732103.1 | 0.92 | 1 | 0.605 | 0/18 | 0.267 | Sensitive |
| ATGC021 | Chlamydia_trachomatis_A_5291_uid196770 | 478448991 | YP_007723914.1 | 0.92 | 1 | 0.605 | 0/18 | 0.267 | Sensitive |
| ATGC021 | Chlamydia_trachomatis_A_HAR_13_uid58333 | 76789094 | YP_328180.1 | 0.92 | 1 | 0.605 | 0/18 | 0.267 | Sensitive |
| ATGC021 | Chlamydia_trachomatis_B_Jali20_OT_uid59351 | 237802791 | YP_002887985.1 | 0.92 | 1 | 0.605 | 0/18 | 0.267 | Sensitive |
| ATGC021 | Chlamydia_trachomatis_B_TZ1A828_OT_uid59349 | 237804713 | YP_002888867.1 | 0.92 | 1 | 0.605 | 0/18 | 0.267 | Sensitive |
| ATGC021 | Chlamydia_trachomatis_D_EC_uid159881 | 385243573 | YP_005811419.1 | 0.92 | 1 | 0.605 | 0/18 | 0.267 | Sensitive |
| ATGC021 | Chlamydia_trachomatis_D_LC_uid159879 | 385244453 | YP_005812297.1 | 0.92 | 1 | 0.605 | 0/18 | 0.267 | Sensitive |

| | | | | | | | | | |
|---|---|---|---|---|---|---|---|---|---|
| ATGC021 | Chlamydia_trachomatis_D_SotonD5_uid196773 | 478455300 | YP_007727490.1 | 0.92 | 1 | 0.605 | 0/18 | 0.267 | Sensitive |
| ATGC021 | Chlamydia_trachomatis_D_UW_3_CX_uid57637 | 15605090 | NP_219875.1 | 0.92 | 1 | 0.605 | 0/18 | 0.267 | Sensitive |
| ATGC021 | Chlamydia_trachomatis_E_11023_uid161369 | 385241738 | YP_005809578.1 | 0.92 | 1 | 0.605 | 0/18 | 0.267 | Sensitive |
| ATGC021 | Chlamydia_trachomatis_E_150_uid161403 | 385245345 | YP_005814168.1 | 0.92 | 1 | 0.605 | 0/18 | 0.267 | Sensitive |
| ATGC021 | Chlamydia_trachomatis_E_Bour_uid196775 | 478461624 | YP_007736574.1 | 0.92 | 1 | 0.605 | 0/18 | 0.267 | Sensitive |
| ATGC021 | Chlamydia_trachomatis_E_SW3_uid167483 | 389858934 | YP_006361175.1 | 0.92 | 1 | 0.605 | 0/18 | 0.267 | Sensitive |
| ATGC021 | Chlamydia_trachomatis_F_SW4_uid167484 | 389858058 | YP_006360300.1 | 0.92 | 1 | 0.605 | 0/18 | 0.267 | Sensitive |
| ATGC021 | Chlamydia_trachomatis_F_SW5_uid167485 | 389859810 | YP_006362050.1 | 0.92 | 1 | 0.605 | 0/18 | 0.267 | Sensitive |
| ATGC021 | Chlamydia_trachomatis_G_11074_uid161409 | 385246268 | YP_005815090.1 | 0.92 | 1 | 0.605 | 0/18 | 0.267 | Sensitive |
| ATGC021 | Chlamydia_trachomatis_G_11222_uid161361 | 385240805 | YP_005808646.1 | 0.92 | 1 | 0.605 | 0/18 | 0.267 | Sensitive |
| ATGC021 | Chlamydia_trachomatis_G_9301_uid161377 | 385242658 | YP_005810497.1 | 0.92 | 1 | 0.605 | 0/18 | 0.267 | Sensitive |
| ATGC021 | Chlamydia_trachomatis_G_9768_uid161353 | 385239882 | YP_005807724.1 | 0.92 | 1 | 0.605 | 0/18 | 0.267 | Sensitive |
| ATGC021 | Chlamydia_trachomatis_G_SotonG1_uid196779 | 478457109 | YP_007725702.1 | 0.92 | 1 | 0.605 | 0/18 | 0.267 | Sensitive |
| ATGC021 | Chlamydia_trachomatis_IU824_uid193712 | 471328538 | YP_007540063.1 | 0.92 | 1 | 0.605 | 0/18 | 0.267 | Sensitive |
| ATGC021 | Chlamydia_trachomatis_IU888_uid193713 | 478428499 | YP_007540939.1 | 0.92 | 1 | 0.605 | 0/18 | 0.267 | Sensitive |
| ATGC021 | Chlamydia_trachomatis_Ia_SotonIa1_uid196780 | 478460725 | YP_007735676.1 | 0.92 | 1 | 0.605 | 0/18 | 0.267 | Sensitive |
| ATGC021 | Chlamydia_trachomatis_K_SotonK1_uid196782 | 478458019 | YP_007731204.1 | 0.92 | 1 | 0.605 | 0/18 | 0.267 | Sensitive |
| ATGC021 | Chlamydia_trachomatis_L1_115_uid196784 | 478451690 | YP_007714996.1 | 0.92 | 1 | 0.605 | 0/18 | 0.267 | Sensitive |
| ATGC021 | Chlamydia_trachomatis_L1_224_uid196785 | 478462529 | YP_007738350.1 | 0.92 | 1 | 0.605 | 0/18 | 0.267 | Sensitive |
| ATGC021 | Chlamydia_trachomatis_L1_440_LN_uid196783 | 478448090 | YP_007722124.1 | 0.92 | 1 | 0.605 | 0/18 | 0.267 | Sensitive |
| ATGC021 | Chlamydia_trachomatis_L2_25667R_uid196786 | 478447198 | YP_007715886.1 | 0.92 | 1 | 0.605 | 0/18 | 0.267 | Sensitive |

| | | | | | | | | | |
|---|---|---|---|---|---|---|---|---|---|
| ATGC021 | Chlamydia_trachomatis_L2_434_Bu_f__uid198644 | 482546674 | YP_007852590.1 | 0.92 | 1 | 0.605 | 0/18 | 0.267 | Sensitive |
| ATGC021 | Chlamydia_trachomatis_L2_434_Bu_i__uid198643 | 482545731 | YP_007851649.1 | 0.92 | 1 | 0.605 | 0/18 | 0.267 | Sensitive |
| ATGC021 | Chlamydia_trachomatis_L2b_795_uid196791 | 478469734 | YP_007723014.1 | 0.92 | 1 | 0.605 | 0/18 | 0.267 | Sensitive |
| ATGC021 | Chlamydia_trachomatis_L2b_8200_07_uid196787 | 478470633 | YP_007729274.1 | 0.92 | 1 | 0.605 | 0/18 | 0.267 | Sensitive |
| ATGC021 | Chlamydia_trachomatis_L2b_Ams1_uid196792 | 478467024 | YP_007718556.1 | 0.92 | 1 | 0.605 | 0/18 | 0.267 | Sensitive |
| ATGC021 | Chlamydia_trachomatis_L2b_UCH_1_proctitis_uid61635 | 166155452 | YP_001653707.1 | 0.92 | 1 | 0.605 | 0/18 | 0.267 | Sensitive |
| ATGC021 | Chlamydia_trachomatis_L2b_UCH_2_uid196788 | 478454395 | YP_007716774.1 | 0.92 | 1 | 0.605 | 0/18 | 0.267 | Sensitive |
| ATGC021 | Chlamydia_trachomatis_L2c_uid68843 | 339626037 | YP_004717516.1 | 0.92 | 1 | 0.605 | 0/18 | 0.267 | Sensitive |
| ATGC021 | Chlamydia_trachomatis_L3_404_LN_uid196797 | 478463428 | YP_007739240.1 | 0.92 | 1 | 0.605 | 0/18 | 0.267 | Sensitive |
| ATGC021 | Chlamydia_trachomatis_Sweden2_uid161995 | 386262719 | YP_005815998.1 | 0.92 | 1 | 0.605 | 0/18 | 0.267 | Sensitive |
| ATGC021 | Chlamydia_trachomatis_uid196771 | 478466136 | YP_007728387.1 | 0.92 | 1 | 0.605 | 0/18 | 0.267 | Sensitive |
| ATGC021 | Chlamydia_trachomatis_uid196776 | 478473334 | YP_007734782.1 | 0.92 | 1 | 0.605 | 0/18 | 0.267 | Sensitive |
| ATGC021 | Chlamydia_trachomatis_uid196781 | 478475140 | YP_007724809.1 | 0.92 | 1 | 0.605 | 0/18 | 0.267 | Sensitive |
| ATGC022 | Chlamydia_psittaci_01DC12_uid179070 | 410858661 | YP_006974601.1 | 0.92 | 1 | 0.526 | 0/18 | 0.4 | Sensitive |
| ATGC022 | Chlamydia_psittaci_84_55_uid175571 | 407454277 | YP_006733385.1 | 0.88 | 0.944 | 0.526 | 0/18 | 0.4 | Unclassified |
| ATGC022 | Chlamydia_psittaci_GR9_uid175572 | 407455553 | YP_006734444.1 | 0.88 | 0.944 | 0.526 | 0/18 | 0.4 | Unclassified |
| ATGC022 | Chlamydia_psittaci_M56_uid175576 | 407459538 | YP_006737641.1 | 0.88 | 0.944 | 0.526 | 0/18 | 0.4 | Unclassified |
| ATGC022 | Chlamydia_psittaci_MN_uid175573 | 406594195 | YP_006741886.1 | 0.92 | 1 | 0.526 | 0/18 | 0.4 | Sensitive |
| ATGC022 | Chlamydia_psittaci_VS225_uid175574 | 407456967 | YP_006735540.1 | 0.88 | 0.944 | 0.526 | 0/18 | 0.4 | Unclassified |
| ATGC022 | Chlamydia_psittaci_WC_uid175577 | 407460909 | YP_006738684.1 | 0.88 | 0.944 | 0.526 | 0/18 | 0.4 | Unclassified |
| ATGC022 | Chlamydia_psittaci_WS_RT_E30_uid175575 | 407458288 | YP_006736593.1 | 0.88 | 0.944 | 0.526 | 0/18 | 0.4 | Unclassified |

| | | | | | | | | | |
|---|---|---|---|---|---|---|---|---|---|
| ATGC022 | Chlamydophila_abortus_S26_3_uid57963 | 62185303 | YP_220088.1 | 0.92 | 1 | 0.5 | 0/18 | 0.4 | Sensitive |
| ATGC022 | Chlamydophila_psittaci_01DC11_uid159527 | 384451800 | YP_005664398.1 | 0.88 | 0.944 | 0.526 | 0/18 | 0.4 | Unclassified |
| ATGC022 | Chlamydophila_psittaci_02DC15_uid159521 | 384454732 | YP_005667327.1 | 0.88 | 0.944 | 0.526 | 0/18 | 0.4 | Unclassified |
| ATGC022 | Chlamydophila_psittaci_08DC60_uid159525 | 384452774 | YP_005665371.1 | 0.88 | 0.944 | 0.526 | 0/18 | 0.4 | Unclassified |
| ATGC022 | Chlamydophila_psittaci_6BC_uid159845 | 384450805 | YP_005663405.1 | 0.88 | 0.944 | 0.526 | 0/18 | 0.4 | Unclassified |
| ATGC022 | Chlamydophila_psittaci_6BC_uid63621 | 332287648 | YP_004422549.1 | 0.88 | 0.944 | 0.526 | 0/18 | 0.4 | Unclassified |
| ATGC022 | Chlamydophila_psittaci_C19_98_uid159523 | 384453753 | YP_005666349.1 | 0.88 | 0.944 | 0.526 | 0/18 | 0.4 | Unclassified |
| ATGC022 | Chlamydophila_psittaci_CP3_uid175578 | 406592606 | YP_006739786.1 | 0.28 | 0.222 | 0.132 | 0/18 | 0.367 | Unclassified |
| ATGC022 | Chlamydophila_psittaci_CP3_uid175578 | 406592607 | YP_006739787.1 | 0.6 | 0.722 | 0.421 | 0/18 | 0.1 | Unclassified |
| ATGC022 | Chlamydophila_psittaci_Mat116_uid189026 | 449071367 | YP_007438447.1 | 0.88 | 0.944 | 0.526 | 0/18 | 0.4 | Unclassified |
| ATGC022 | Chlamydophila_psittaci_NJ1_uid175579 | 406593666 | YP_006740845.1 | 0.88 | 0.944 | 0.526 | 0/18 | 0.4 | Unclassified |
| ATGC022 | Chlamydophila_psittaci_RD1_uid162063 | 392376878 | YP_004064656.1 | 0.88 | 0.944 | 0.526 | 0/18 | 0.4 | Unclassified |
| ATGC025 | Mycobacterium_africanum_GM041182_uid68839 | 339633238 | YP_004724880.1 | 1 | 0.889 | 0.5 | 0/18 | 0.467 | Sensitive |
| ATGC025 | Mycobacterium_bovis_AF2122_97_uid57695 | 31794408 | NP_856901.1 | 1 | 0.889 | 0.5 | 0/18 | 0.467 | Sensitive |
| ATGC025 | Mycobacterium_bovis_BCG_Korea_1168P_uid189029 | 449065332 | YP_007432415.1 | 1 | 0.889 | 0.5 | 0/18 | 0.467 | Sensitive |
| ATGC025 | Mycobacterium_bovis_BCG_Korea_1168P_uid189029 | 461474902 | YP_007517340.1 | 1 | 0.889 | 0.5 | 0/18 | 0.467 | Sensitive |
| ATGC025 | Mycobacterium_bovis_BCG_Korea_1168P_uid189029 | 461474936 | YP_007517374.1 | 1 | 0.889 | 0.5 | 0/18 | 0.467 | Sensitive |
| ATGC025 | Mycobacterium_bovis_BCG_Mexico_uid86889 | 378772976 | YP_005172709.1 | 1 | 0.889 | 0.5 | 0/18 | 0.467 | Sensitive |
| ATGC025 | Mycobacterium_bovis_BCG_Mexico_uid86889 | 378773069 | YP_005172802.1 | 1 | 0.889 | 0.5 | 0/18 | 0.467 | Sensitive |
| ATGC025 | Mycobacterium_bovis_BCG_Pasteur_1173P2_uid58781 | 121639117 | YP_979341.1 | 1 | 0.889 | 0.5 | 0/18 | 0.467 | Sensitive |
| ATGC025 | Mycobacterium_bovis_BCG_Pasteur_1173P2_uid58781 | 121639209 | YP_979433.1 | 1 | 0.889 | 0.5 | 0/18 | 0.467 | Sensitive |

| | | | | | | | | | |
|---|---|---|---|---|---|---|---|---|---|
| ATGC025 | Mycobacterium_bovis_BCG_Tokyo_172_uid59281 | 224991609 | YP_002646298.1 | 1 | 0.889 | 0.5 | 0/18 | 0.467 | Sensitive |
| ATGC025 | Mycobacterium_canettii_CIPT_140010059_uid70731 | 340628207 | YP_004746659.1 | 1 | 0.889 | 0.5 | 0/18 | 0.467 | Sensitive |
| ATGC025 | Mycobacterium_canettii_CIPT_140060008_uid184829 | 433628362 | YP_007261991.1 | 1 | 0.889 | 0.5 | 0/18 | 0.467 | Sensitive |
| ATGC025 | Mycobacterium_canettii_CIPT_140070008_uid184832 | 433643421 | YP_007289180.1 | 1 | 0.889 | 0.5 | 0/18 | 0.467 | Sensitive |
| ATGC025 | Mycobacterium_canettii_CIPT_140070010_uid184828 | 433632327 | YP_007265955.1 | 1 | 0.889 | 0.5 | 0/18 | 0.467 | Sensitive |
| ATGC025 | Mycobacterium_canettii_CIPT_140070017_uid184830 | 433636323 | YP_007269950.1 | 1 | 0.889 | 0.5 | 0/18 | 0.467 | Sensitive |
| ATGC025 | Mycobacterium_tuberculosis_Beijing_NITR203_uid197218 | 479316217 | YP_007856269.1 | 1 | 0.889 | 0.5 | 0/18 | 0.467 | Sensitive |
| ATGC025 | Mycobacterium_tuberculosis_CCDC5079_uid161943 | 385996103 | YP_005914401.1 | 1 | 0.889 | 0.5 | 0/18 | 0.467 | Sensitive |
| ATGC025 | Mycobacterium_tuberculosis_CCDC5079_uid203790 | 506927803 | YP_008008253.1 | 1 | 0.889 | 0.5 | 0/18 | 0.467 | Sensitive |
| ATGC025 | Mycobacterium_tuberculosis_CCDC5180_uid161941 | 385992474 | YP_005910772.1 | 1 | 0.889 | 0.5 | 0/18 | 0.467 | Sensitive |
| ATGC025 | Mycobacterium_tuberculosis_CDC1551_uid57775 | 15842816 | NP_337853.1 | 1 | 0.889 | 0.5 | 0/18 | 0.467 | Sensitive |
| ATGC025 | Mycobacterium_tuberculosis_CTRI_2_uid161997 | 386000016 | YP_005918315.1 | 1 | 0.889 | 0.5 | 0/18 | 0.467 | Sensitive |
| ATGC025 | Mycobacterium_tuberculosis_EAI5_NITR206_uid202218 | 494701686 | YP_007967168.1 | 1 | 0.889 | 0.5 | 0/18 | 0.467 | Sensitive |
| ATGC025 | Mycobacterium_tuberculosis_Erdman___ATCC_35801_uid193763 | 471339278 | YP_007611991.1 | 1 | 0.889 | 0.5 | 0/18 | 0.467 | Sensitive |
| ATGC025 | Mycobacterium_tuberculosis_F11_uid58417 | 148824429 | YP_001289183.1 | 1 | 0.889 | 0.5 | 0/18 | 0.467 | Sensitive |
| ATGC025 | Mycobacterium_tuberculosis_H37Ra_uid58853 | 148663090 | YP_001284613.1 | 1 | 0.889 | 0.5 | 0/18 | 0.467 | Sensitive |
| ATGC025 | Mycobacterium_tuberculosis_H37Rv_uid170532 | 397675168 | YP_006516703.1 | 1 | 0.889 | 0.5 | 0/18 | 0.467 | Sensitive |
| ATGC025 | Mycobacterium_tuberculosis_H37Rv_uid57777 | 15610363 | NP_217744.1 | 1 | 0.889 | 0.5 | 0/18 | 0.467 | Sensitive |
| ATGC025 | Mycobacterium_tuberculosis_KZN_1435_uid59069 | 253800269 | YP_003033270.1 | 1 | 0.889 | 0.5 | 0/18 | 0.467 | Sensitive |
| ATGC025 | Mycobacterium_tuberculosis_KZN_4207_uid83619 | 375297499 | YP_005101766.1 | 1 | 0.889 | 0.5 | 0/18 | 0.467 | Sensitive |
| ATGC025 | Mycobacterium_tuberculosis_KZN_605_uid54947 | 392433709 | YP_006474753.1 | 1 | 0.889 | 0.5 | 0/18 | 0.467 | Sensitive |

| | | | | | | | | | |
|---|---|---|---|---|---|---|---|---|---|
| ATGC025 | Mycobacterium_tuberculosis_RGTB327_uid157907 | 383308969 | YP_005361780.1 | 1 | 0.889 | 0.5 | 0/18 | 0.467 | Sensitive |
| ATGC025 | Mycobacterium_tuberculosis_RGTB423_uid162179 | 386006070 | YP_005924349.1 | 1 | 0.889 | 0.5 | 0/18 | 0.467 | Sensitive |
| ATGC025 | Mycobacterium_tuberculosis_UT205_uid162183 | 392387851 | YP_005309480.1 | 1 | 0.889 | 0.5 | 0/18 | 0.467 | Sensitive |
| ATGC025 | Mycobacterium_tuberculosis_uid185758 | 479057200 | YP_007353006.1 | 1 | 0.889 | 0.5 | 0/18 | 0.467 | Sensitive |
| ATGC052 | Helicobacter_acinonychis_Sheeba_uid58685 | 109947061 | YP_664289.1 | 0.76 | 0.833 | 0.947 | 0/18 | 0.233 | Unclassified |
| ATGC052 | Helicobacter_cetorum_MIT_99_5656_uid162215 | 386747550 | YP_006220758.1 | 0.72 | 0.778 | 0.947 | 0/18 | 0.233 | Unclassified |
| ATGC052 | Helicobacter_pylori_2017_uid161151 | 385224085 | YP_005784011.1 | 0.76 | 0.833 | 0.947 | 0/18 | 0.233 | Unclassified |
| ATGC052 | Helicobacter_pylori_2018_uid161159 | 385231941 | YP_005791860.1 | 0.76 | 0.833 | 0.947 | 0/18 | 0.233 | Unclassified |
| ATGC052 | Helicobacter_pylori_26695_uid178201 | 410681923 | YP_006934325.1 | 0.76 | 0.833 | 0.947 | 0/18 | 0.267 | Unclassified |
| ATGC052 | Helicobacter_pylori_26695_uid57787 | 15645029 | NP_207199.1 | 0.76 | 0.833 | 0.947 | 0/18 | 0.267 | Unclassified |
| ATGC052 | Helicobacter_pylori_35A_uid49903 | 384896326 | YP_005770315.1 | 0.76 | 0.833 | 0.947 | 0/18 | 0.233 | Unclassified |
| ATGC052 | Helicobacter_pylori_51_uid161925 | 387782627 | YP_005793340.1 | 0.76 | 0.833 | 0.947 | 0/18 | 0.233 | Unclassified |
| ATGC052 | Helicobacter_pylori_83_uid161153 | 385225708 | YP_005785633.1 | 0.76 | 0.833 | 0.947 | 0/18 | 0.233 | Unclassified |
| ATGC052 | Helicobacter_pylori_908_uid159985 | 384891406 | YP_005765539.1 | 0.76 | 0.833 | 0.947 | 0/18 | 0.233 | Unclassified |
| ATGC052 | Helicobacter_pylori_Aklavik117_uid182201 | 425789595 | YP_007017515.1 | 0.76 | 0.833 | 0.947 | 0/18 | 0.233 | Unclassified |
| ATGC052 | Helicobacter_pylori_Aklavik86_uid182202 | 425791258 | YP_007019175.1 | 0.76 | 0.833 | 0.947 | 0/18 | 0.233 | Unclassified |
| ATGC052 | Helicobacter_pylori_B38_uid59415 | 254779607 | YP_003057713.1 | 0.76 | 0.833 | 0.947 | 0/18 | 0.233 | Unclassified |
| ATGC052 | Helicobacter_pylori_B8_uid49873 | 298735948 | YP_003728473.1 | 0.76 | 0.833 | 0.947 | 0/18 | 0.233 | Unclassified |
| ATGC052 | Helicobacter_pylori_Cuz20_uid159987 | 384893031 | YP_005767124.1 | 0.76 | 0.833 | 0.947 | 0/18 | 0.233 | Unclassified |
| ATGC052 | Helicobacter_pylori_ELS37_uid158157 | 383749330 | YP_005424433.1 | 0.76 | 0.833 | 0.947 | 0/18 | 0.233 | Unclassified |
| ATGC052 | Helicobacter_pylori_F16_uid161145 | 385217754 | YP_005779230.1 | 0.76 | 0.833 | 0.947 | 0/18 | 0.233 | Unclassified |

| | | | | | | | | | |
|---|---|---|---|---|---|---|---|---|---|
| ATGC052 | Helicobacter_pylori_F30_uid159991 | 384898700 | YP_005774079.1 | 0.76 | 0.833 | 0.947 | 0/18 | 0.233 | Unclassified |
| ATGC052 | Helicobacter_pylori_F32_uid161139 | 385215619 | YP_005775575.1 | 0.76 | 0.833 | 0.947 | 0/18 | 0.233 | Unclassified |
| ATGC052 | Helicobacter_pylori_F57_uid161143 | 385249511 | YP_005777730.1 | 0.76 | 0.833 | 0.947 | 0/18 | 0.233 | Unclassified |
| ATGC052 | Helicobacter_pylori_G27_uid59305 | 208434949 | YP_002266615.1 | 0.76 | 0.833 | 0.947 | 0/18 | 0.233 | Unclassified |
| ATGC052 | Helicobacter_pylori_Gambia94_24_uid159493 | 385219279 | YP_005780754.1 | 0.76 | 0.833 | 0.947 | 0/18 | 0.2 | Unclassified |
| ATGC052 | Helicobacter_pylori_HPAG1_uid58517 | 108563416 | YP_627732.1 | 0.76 | 0.833 | 0.947 | 0/18 | 0.233 | Unclassified |
| ATGC052 | Helicobacter_pylori_HUP_B14_uid162213 | 386746470 | YP_006219687.1 | 0.76 | 0.833 | 0.947 | 0/18 | 0.233 | Unclassified |
| ATGC052 | Helicobacter_pylori_India7_uid161149 | 385220855 | YP_005782327.1 | 0.76 | 0.833 | 0.947 | 0/18 | 0.233 | Unclassified |
| ATGC052 | Helicobacter_pylori_J99_uid57789 | 15612045 | NP_223697.1 | 0.76 | 0.833 | 0.947 | 0/18 | 0.233 | Unclassified |
| ATGC052 | Helicobacter_pylori_Lithuania75_uid159491 | 384897721 | YP_005773149.1 | 0.76 | 0.833 | 0.947 | 0/18 | 0.233 | Unclassified |
| ATGC052 | Helicobacter_pylori_OK113_uid193715 | 470165414 | YP_007537128.1 | 0.76 | 0.833 | 0.947 | 0/18 | 0.233 | Unclassified |
| ATGC052 | Helicobacter_pylori_OK310_uid193716 | 478429973 | YP_007538585.1 | 0.76 | 0.833 | 0.947 | 0/18 | 0.233 | Unclassified |
| ATGC052 | Helicobacter_pylori_P12_uid59327 | 210135212 | YP_002301651.1 | 0.76 | 0.833 | 0.947 | 0/18 | 0.233 | Unclassified |
| ATGC052 | Helicobacter_pylori_PeCan18_uid162211 | 386756048 | YP_006229265.1 | 0.76 | 0.833 | 0.947 | 0/18 | 0.233 | Unclassified |
| ATGC052 | Helicobacter_pylori_PeCan4_uid53539 | 308183156 | YP_003927283.1 | 0.76 | 0.833 | 0.947 | 0/18 | 0.2 | Unclassified |
| ATGC052 | Helicobacter_pylori_Puno120_uid159611 | 385228733 | YP_005788666.1 | 0.76 | 0.833 | 0.947 | 0/18 | 0.233 | Unclassified |
| ATGC052 | Helicobacter_pylori_Puno135_uid161157 | 385230335 | YP_005790251.1 | 0.76 | 0.833 | 0.947 | 0/18 | 0.233 | Unclassified |
| ATGC052 | Helicobacter_pylori_Rif1_uid178202 | 410023637 | YP_006892890.1 | 0.76 | 0.833 | 0.947 | 0/18 | 0.267 | Unclassified |
| ATGC052 | Helicobacter_pylori_Rif2_uid178203 | 410501404 | YP_006935931.1 | 0.76 | 0.833 | 0.947 | 0/18 | 0.267 | Unclassified |
| ATGC052 | Helicobacter_pylori_SJM180_uid53541 | 308184791 | YP_003928924.1 | 0.76 | 0.833 | 0.947 | 0/18 | 0.233 | Unclassified |
| ATGC052 | Helicobacter_pylori_SNT49_uid159615 | 385227238 | YP_005787162.1 | 0.76 | 0.833 | 0.947 | 0/18 | 0.2 | Unclassified |

| | | | | | | | | | |
|---|---|---|---|---|---|---|---|---|---|
| ATGC052 | Helicobacter_pylori_Sat464_uid159467 | 384894581 | YP_005768630.1 | 0.76 | 0.833 | 0.947 | 0/18 | 0.233 | Unclassified |
| ATGC052 | Helicobacter_pylori_Shi112_uid162207 | 386754513 | YP_006227731.1 | 0.76 | 0.833 | 0.947 | 0/18 | 0.233 | Unclassified |
| ATGC052 | Helicobacter_pylori_Shi169_uid162209 | 386752982 | YP_006226201.1 | 0.76 | 0.833 | 0.947 | 0/18 | 0.2 | Unclassified |
| ATGC052 | Helicobacter_pylori_Shi417_uid162205 | 386751400 | YP_006224620.1 | 0.76 | 0.833 | 0.947 | 0/18 | 0.233 | Unclassified |
| ATGC052 | Helicobacter_pylori_Shi470_uid59165 | 188527834 | YP_001910521.1 | 0.76 | 0.833 | 0.947 | 0/18 | 0.233 | Unclassified |
| ATGC052 | Helicobacter_pylori_SouthAfrica7_uid159989 | 385222469 | YP_005771602.1 | 0.72 | 0.778 | 0.947 | 0/18 | 0.233 | Unclassified |
| ATGC052 | Helicobacter_pylori_UM032_uid203025 | 499081750 | YP_007979358.1 | 0.64 | 0.667 | 0.842 | 0/18 | 0.233 | Unclassified |
| ATGC052 | Helicobacter_pylori_UM037_uid203027 | 499087520 | YP_007982704.1 | 0.76 | 0.833 | 0.947 | 0/18 | 0.233 | Unclassified |
| ATGC052 | Helicobacter_pylori_UM066_uid203028 | 499084224 | YP_007983467.1 | 0.76 | 0.833 | 0.947 | 0/18 | 0.233 | Unclassified |
| ATGC052 | Helicobacter_pylori_UM299_uid203026 | 499082296 | YP_007979903.1 | 0.76 | 0.833 | 0.947 | 0/18 | 0.233 | Unclassified |
| ATGC052 | Helicobacter_pylori_XZ274_uid165869 | 387908310 | YP_006338644.1 | 0.76 | 0.833 | 0.947 | 0/18 | 0.2 | Unclassified |
| ATGC052 | Helicobacter_pylori_uid159983 | 384887956 | YP_005762467.1 | 0.76 | 0.833 | 0.947 | 0/18 | 0.233 | Unclassified |
| ATGC052 | Helicobacter_pylori_v225d_uid159639 | 384889663 | YP_005763965.1 | 0.76 | 0.833 | 0.947 | 0/18 | 0.233 | Unclassified |
| ATGC054 | Staphylococcus_aureus_04_02981_uid161969 | 387150605 | YP_005742169.1 | 0.76 | 0.778 | 1 | 0/18 | 0.267 | Resistant |
| ATGC054 | Staphylococcus_aureus_08BA02176_uid175257 | 404478812 | YP_006710242.1 | 0.76 | 0.778 | 1 | 0/18 | 0.267 | Resistant |
| ATGC054 | Staphylococcus_aureus_11819_97_uid159981 | 385781690 | YP_005757861.1 | 0.76 | 0.778 | 1 | 0/18 | 0.267 | Resistant |
| ATGC054 | Staphylococcus_aureus_71193_uid162141 | 386729163 | YP_006195546.1 | 0.76 | 0.778 | 1 | 0/18 | 0.267 | Resistant |
| ATGC054 | Staphylococcus_aureus_CC45_uid209174 | 514066339 | YP_008128259.1 | 0.76 | 0.778 | 1 | 0/18 | 0.267 | Resistant |
| ATGC054 | Staphylococcus_aureus_COL_uid57797 | 57650419 | YP_186348.1 | 0.76 | 0.778 | 1 | 0/18 | 0.267 | Resistant |
| ATGC054 | Staphylococcus_aureus_ECT_R_2_uid159389 | 384864689 | YP_005750048.1 | 0.76 | 0.778 | 1 | 0/18 | 0.267 | Resistant |
| ATGC054 | Staphylococcus_aureus_ED133_uid159689 | 384547706 | YP_005736959.1 | 0.76 | 0.778 | 1 | 0/18 | 0.267 | Resistant |

| | | | | | | | | | |
|---|---|---|---|---|---|---|---|---|---|
| ATGC054 | Staphylococcus_aureus_ED98_uid41455 | 269203089 | YP_003282358.1 | 0.76 | 0.778 | 1 | 0/18 | 0.267 | Resistant |
| ATGC054 | Staphylococcus_aureus_HO_5096_0412_uid162163 | 386831016 | YP_006237670.1 | 0.76 | 0.778 | 1 | 0/18 | 0.267 | Resistant |
| ATGC054 | Staphylococcus_aureus_JH1_uid58457 | 150394013 | YP_001316688.1 | 0.76 | 0.778 | 1 | 0/18 | 0.267 | Resistant |
| ATGC054 | Staphylococcus_aureus_JH9_uid58455 | 148267950 | YP_001246893.1 | 0.76 | 0.778 | 1 | 0/18 | 0.267 | Resistant |
| ATGC054 | Staphylococcus_aureus_JKD6008_uid159855 | 384862066 | YP_005744786.1 | 0.76 | 0.778 | 1 | 0/18 | 0.267 | Resistant |
| ATGC054 | Staphylococcus_aureus_JKD6159_uid159691 | 384550226 | YP_005739478.1 | 0.76 | 0.778 | 1 | 0/18 | 0.267 | Resistant |
| ATGC054 | Staphylococcus_aureus_LGA251_uid159391 | 387780565 | YP_005755363.1 | 0.76 | 0.778 | 1 | 0/18 | 0.267 | Resistant |
| ATGC054 | Staphylococcus_aureus_M013_uid88065 | 379021179 | YP_005297841.1 | 0.76 | 0.778 | 1 | 0/18 | 0.267 | Resistant |
| ATGC054 | Staphylococcus_aureus_M1_uid197263 | 479329406 | YP_007868465.1 | 0.76 | 0.778 | 1 | 0/18 | 0.267 | Resistant |
| ATGC054 | Staphylococcus_aureus_MRSA252_uid57839 | 49483653 | YP_040877.1 | 0.76 | 0.778 | 1 | 0/18 | 0.267 | Resistant |
| ATGC054 | Staphylococcus_aureus_MSSA476_uid57841 | 49486304 | YP_043525.1 | 0.76 | 0.778 | 1 | 0/18 | 0.267 | Resistant |
| ATGC054 | Staphylococcus_aureus_MW2_uid57903 | 21283083 | NP_646171.1 | 0.76 | 0.778 | 1 | 0/18 | 0.267 | Resistant |
| ATGC054 | Staphylococcus_aureus_Mu3_uid58817 | 156979783 | YP_001442042.1 | 0.76 | 0.778 | 1 | 0/18 | 0.267 | Resistant |
| ATGC054 | Staphylococcus_aureus_Mu50_uid57835 | 15924454 | NP_371988.1 | 0.76 | 0.778 | 1 | 0/18 | 0.267 | Resistant |
| ATGC054 | Staphylococcus_aureus_N315_uid57837 | 15927045 | NP_374578.1 | 0.76 | 0.778 | 1 | 0/18 | 0.267 | Resistant |
| ATGC054 | Staphylococcus_aureus_NCTC_8325_uid57795 | 88195198 | YP_499999.1 | 0.76 | 0.778 | 1 | 0/18 | 0.267 | Resistant |
| ATGC054 | Staphylococcus_aureus_Newman_uid58839 | 151221587 | YP_001332409.1 | 0.76 | 0.778 | 1 | 0/18 | 0.267 | Resistant |
| ATGC054 | Staphylococcus_aureus_RF122_uid57661 | 82751063 | YP_416804.1 | 0.76 | 0.778 | 1 | 0/18 | 0.267 | Resistant |
| ATGC054 | Staphylococcus_aureus_ST228_10388_uid193754 | 470191350 | YP_007577660.1 | 0.76 | 0.778 | 1 | 0/18 | 0.267 | Resistant |
| ATGC054 | Staphylococcus_aureus_ST228_10497_uid193755 | 470218983 | YP_007619377.1 | 0.76 | 0.778 | 1 | 0/18 | 0.267 | Resistant |
| ATGC054 | Staphylococcus_aureus_ST228_15532_uid193756 | 470193316 | YP_007579669.1 | 0.76 | 0.778 | 1 | 0/18 | 0.267 | Resistant |

| | | | | | | | | | |
|---|---|---|---|---|---|---|---|---|---|
| ATGC054 | Staphylococcus_aureus_ST228_16035_uid193757 | 470195281 | YP_007581633.1 | 0.76 | 0.778 | 1 | 0/18 | 0.267 | Resistant |
| ATGC054 | Staphylococcus_aureus_ST228_18412_uid193760 | 470199214 | YP_007585614.1 | 0.76 | 0.778 | 1 | 0/18 | 0.267 | Resistant |
| ATGC054 | Staphylococcus_aureus_ST398_uid159247 | 387602739 | YP_005734260.1 | 0.76 | 0.778 | 1 | 0/18 | 0.267 | Resistant |
| ATGC054 | Staphylococcus_aureus_T0131_uid159861 | 384870006 | YP_005752720.1 | 0.76 | 0.778 | 1 | 0/18 | 0.267 | Resistant |
| ATGC054 | Staphylococcus_aureus_TCH60_uid159859 | 384867626 | YP_005747822.1 | 0.76 | 0.778 | 1 | 0/18 | 0.267 | Resistant |
| ATGC054 | Staphylococcus_aureus_TW20_uid159241 | 387143074 | YP_005731467.1 | 0.76 | 0.778 | 1 | 0/18 | 0.267 | Resistant |
| ATGC054 | Staphylococcus_aureus_USA300_FPR3757_uid58555 | 87159926 | YP_494052.1 | 0.76 | 0.778 | 1 | 0/18 | 0.267 | Resistant |
| ATGC054 | Staphylococcus_aureus_USA300_TCH1516_uid58925 | 161509632 | YP_001575291.1 | 0.76 | 0.778 | 1 | 0/18 | 0.267 | Resistant |
| ATGC054 | Staphylococcus_aureus_VC40_uid88071 | 379014674 | YP_005290910.1 | 0.76 | 0.778 | 1 | 0/18 | 0.267 | Resistant |
| ATGC054 | Staphylococcus_aureus_uid193758 | 470222804 | YP_007621361.1 | 0.76 | 0.778 | 1 | 0/18 | 0.267 | Resistant |
| ATGC054 | Staphylococcus_aureus_uid193759 | 470197246 | YP_007583647.1 | 0.76 | 0.778 | 1 | 0/18 | 0.267 | Resistant |
| ATGC054 | Staphylococcus_aureus_uid193761 | 470224771 | YP_007623350.1 | 0.76 | 0.778 | 1 | 0/18 | 0.267 | Resistant |
| ATGC067 | Corynebacterium_pseudotuberculosis_1002_uid159677 | 384504127 | YP_005680797.1 | 1 | 0.889 | 0.526 | 0/18 | 0.433 | Sensitive |
| ATGC067 | Corynebacterium_pseudotuberculosis_1_06_A_uid159665 | 387140152 | YP_005696130.1 | 1 | 0.889 | 0.526 | 0/18 | 0.433 | Sensitive |
| ATGC067 | Corynebacterium_pseudotuberculosis_258_uid167260 | 389849880 | YP_006352115.1 | 1 | 0.889 | 0.526 | 0/18 | 0.433 | Sensitive |
| ATGC067 | Corynebacterium_pseudotuberculosis_267_uid162175 | 385806983 | YP_005843380.1 | 1 | 0.889 | 0.526 | 0/18 | 0.433 | Sensitive |
| ATGC067 | Corynebacterium_pseudotuberculosis_316_uid89381 | 379714813 | YP_005303150.1 | 1 | 0.889 | 0.526 | 0/18 | 0.433 | Sensitive |
| ATGC067 | Corynebacterium_pseudotuberculosis_31_uid162167 | 386739874 | YP_006213054.1 | 0.88 | 0.778 | 0.474 | 0/18 | 0.433 | Unclassified |
| ATGC067 | Corynebacterium_pseudotuberculosis_3_99_5_uid83609 | 375288112 | YP_005122653.1 | 1 | 0.889 | 0.526 | 0/18 | 0.433 | Sensitive |
| ATGC067 | Corynebacterium_pseudotuberculosis_42_02_A_uid159669 | 387136069 | YP_005692049.1 | 1 | 0.889 | 0.526 | 0/18 | 0.433 | Sensitive |
| ATGC067 | Corynebacterium_pseudotuberculosis_C231_uid159675 | 384506220 | YP_005682889.1 | 1 | 0.889 | 0.526 | 0/18 | 0.433 | Sensitive |

| | | | | | | | | | |
|---|---|---|---|---|---|---|---|---|---|
| ATGC067 | Corynebacterium_pseudotuberculosis_CIP_52_97_uid159667 | 387138133 | YP_005694112.1 | 0.88 | 0.778 | 0.474 | 0/18 | 0.433 | Unclassified |
| ATGC067 | Corynebacterium_pseudotuberculosis_Cp162_uid168258 | 392400082 | YP_006436682.1 | 0.84 | 0.667 | 0.421 | 0/18 | 0.367 | Unclassified |
| ATGC067 | Corynebacterium_pseudotuberculosis_FRC41_uid50585 | 300857947 | YP_003782930.1 | 1 | 0.889 | 0.526 | 0/18 | 0.433 | Sensitive |
| ATGC067 | Corynebacterium_pseudotuberculosis_I19_uid159673 | 384508308 | YP_005684976.1 | 1 | 0.889 | 0.526 | 0/18 | 0.433 | Sensitive |
| ATGC067 | Corynebacterium_pseudotuberculosis_P54B96_uid157909 | 383313707 | YP_005374562.1 | 1 | 0.889 | 0.526 | 0/18 | 0.433 | Sensitive |
| ATGC067 | Corynebacterium_pseudotuberculosis_PAT10_uid159671 | 384510401 | YP_005689979.1 | 1 | 0.889 | 0.526 | 0/18 | 0.433 | Sensitive |
| ATGC068 | Corynebacterium_diphtheriae_241_uid83607 | 375290373 | YP_005124913.1 | 1 | 0.889 | 0.553 | 0/18 | 0.4 | Sensitive |
| ATGC068 | Corynebacterium_diphtheriae_31A_uid84309 | 376284205 | YP_005157415.1 | 1 | 0.889 | 0.553 | 0/18 | 0.4 | Sensitive |
| ATGC068 | Corynebacterium_diphtheriae_BH8_uid84311 | 376287193 | YP_005159759.1 | 1 | 0.889 | 0.553 | 0/18 | 0.4 | Sensitive |
| ATGC068 | Corynebacterium_diphtheriae_CDCE_8392_uid84295 | 376242348 | YP_005133200.1 | 1 | 0.889 | 0.553 | 0/18 | 0.4 | Sensitive |
| ATGC068 | Corynebacterium_diphtheriae_HC01_uid84297 | 376245206 | YP_005135445.1 | 1 | 0.889 | 0.553 | 0/18 | 0.4 | Sensitive |
| ATGC068 | Corynebacterium_diphtheriae_HC02_uid84317 | 376292762 | YP_005164436.1 | 1 | 0.889 | 0.553 | 0/18 | 0.4 | Sensitive |
| ATGC068 | Corynebacterium_diphtheriae_HC03_uid84299 | 376250799 | YP_005137680.1 | 1 | 0.889 | 0.553 | 0/18 | 0.4 | Sensitive |
| ATGC068 | Corynebacterium_diphtheriae_HC04_uid84301 | 376247977 | YP_005139921.1 | 1 | 0.889 | 0.553 | 0/18 | 0.4 | Sensitive |
| ATGC068 | Corynebacterium_diphtheriae_INCA_402_uid83605 | 375292590 | YP_005127129.1 | 1 | 0.889 | 0.553 | 0/18 | 0.4 | Sensitive |
| ATGC068 | Corynebacterium_diphtheriae_NCTC_13129_uid57691 | 38233313 | NP_939080.1 | 1 | 0.889 | 0.553 | 0/18 | 0.4 | Sensitive |
| ATGC068 | Corynebacterium_diphtheriae_PW8_uid84303 | 376253810 | YP_005142269.1 | 1 | 0.889 | 0.553 | 0/18 | 0.4 | Sensitive |
| ATGC068 | Corynebacterium_diphtheriae_VA01_uid84305 | 376256610 | YP_005144501.1 | 0.88 | 0.778 | 0.5 | 0/18 | 0.4 | Unclassified |
| ATGC082 | Clostridium_botulinum_A2_Kyoto_uid59229 | 226948809 | YP_002803900.1 | 0.72 | 0.778 | 1 | 0/18 | 0.2 | Resistant |
| ATGC082 | Clostridium_botulinum_A3_Loch_Maree_uid59149 | 170760882 | YP_001786896.1 | 0.72 | 0.778 | 1 | 0/18 | 0.2 | Resistant |
| ATGC082 | Clostridium_botulinum_A_ATCC_19397_uid58927 | 153932774 | YP_001383824.1 | 0.72 | 0.778 | 1 | 0/18 | 0.2 | Resistant |

| | | | | | | | | | |
|---|---|---|---|---|---|---|---|---|---|
| ATGC082 | Clostridium_botulinum_A_ATCC_3502_uid61579 | 148379445 | YP_001253986.1 | 0.72 | 0.778 | 1 | 0/18 | 0.2 | Resistant |
| ATGC082 | Clostridium_botulinum_A_Hall_uid58931 | 153934776 | YP_001387374.1 | 0.72 | 0.778 | 1 | 0/18 | 0.2 | Resistant |
| ATGC082 | Clostridium_botulinum_B1_Okra_uid59147 | 170755701 | YP_001781111.1 | 0.72 | 0.778 | 1 | 0/18 | 0.2 | Resistant |
| ATGC082 | Clostridium_botulinum_Ba4_657_uid59173 | 237794811 | YP_002862363.1 | 0.72 | 0.778 | 1 | 0/18 | 0.2 | Resistant |
| ATGC082 | Clostridium_botulinum_F_230613_uid159513 | 384461873 | YP_005674468.1 | 0.72 | 0.778 | 1 | 0/18 | 0.2 | Resistant |
| ATGC082 | Clostridium_botulinum_F_Langeland_uid58929 | 153940675 | YP_001390821.1 | 0.72 | 0.778 | 1 | 0/18 | 0.2 | Resistant |
| ATGC082 | Clostridium_botulinum_H04402_065_uid162091 | 387817749 | YP_005678094.1 | 0.72 | 0.778 | 1 | 0/18 | 0.2 | Resistant |
| ATGC089 | Burkholderia_mallei_ATCC_23344_uid57725 | 53724689 | YP_102068.1 | 0.72 | 0.833 | 0.474 | 0/18 | 0.167 | Unclassified |
| ATGC089 | Burkholderia_mallei_NCTC_10229_uid58383 | 124383336 | YP_001028329.1 | 0.72 | 0.833 | 0.474 | 0/18 | 0.167 | Unclassified |
| ATGC089 | Burkholderia_mallei_NCTC_10247_uid58385 | 126448754 | YP_001081973.1 | 0.72 | 0.833 | 0.474 | 0/18 | 0.167 | Unclassified |
| ATGC089 | Burkholderia_mallei_SAVP1_uid58387 | 121600779 | YP_994007.1 | 0.72 | 0.833 | 0.474 | 0/18 | 0.167 | Unclassified |
| ATGC089 | Burkholderia_pseudomallei_1026b_uid162511 | 386862877 | YP_006275826.1 | 0.72 | 0.833 | 0.474 | 0/18 | 0.167 | Unclassified |
| ATGC089 | Burkholderia_pseudomallei_1106a_uid58515 | 126453471 | YP_001065016.1 | 0.72 | 0.833 | 0.474 | 0/18 | 0.167 | Unclassified |
| ATGC089 | Burkholderia_pseudomallei_1710b_uid58391 | 76811814 | YP_332315.1 | 0.72 | 0.833 | 0.474 | 0/18 | 0.167 | Unclassified |
| ATGC089 | Burkholderia_pseudomallei_668_uid58389 | 126440301 | YP_001057772.1 | 0.72 | 0.833 | 0.474 | 0/18 | 0.167 | Unclassified |
| ATGC089 | Burkholderia_pseudomallei_BPC006_uid174460 | 403517385 | YP_006651518.1 | 0.72 | 0.833 | 0.474 | 0/18 | 0.167 | Unclassified |
| ATGC089 | Burkholderia_pseudomallei_K96243_uid57733 | 53718326 | YP_107312.1 | 0.72 | 0.833 | 0.474 | 0/18 | 0.167 | Unclassified |
| ATGC089 | Burkholderia_pseudomallei_MSHR346_uid55259 | 237810924 | YP_002895375.1 | 0.72 | 0.833 | 0.474 | 0/18 | 0.167 | Unclassified |
| ATGC089 | Burkholderia_thailandensis_MSMB121_uid201037 | 488602285 | YP_007919571.1 | 0.72 | 0.833 | 0.474 | 0/18 | 0.167 | Unclassified |
| ATGC089 | Burkholderia_mallei_NCTC_10229_uid58383 | 124383385 | YP_001028330.1 | 0.56 | 0.611 | 0.368 | 0/18 | 0.267 | Unclassified |
| ATGC089 | Burkholderia_mallei_NCTC_10247_uid58385 | 126449744 | YP_001081974.1 | 0.56 | 0.611 | 0.368 | 0/18 | 0.267 | Unclassified |

| | | | | | | | | | |
|---|---|---|---|---|---|---|---|---|---|
| ATGC089 | Burkholderia_mallei_SAVP1_uid58387 | 121600031 | YP_994006.1 | 0.56 | 0.611 | 0.368 | 0/18 | 0.267 | Unclassified |
| ATGC089 | Burkholderia_pseudomallei_1026b_uid162511 | 386862876 | YP_006275825.1 | 0.56 | 0.611 | 0.316 | 0/18 | 0.267 | Unclassified |
| ATGC089 | Burkholderia_pseudomallei_1106a_uid58515 | 126453347 | YP_001065017.1 | 0.56 | 0.611 | 0.316 | 0/18 | 0.267 | Unclassified |
| ATGC089 | Burkholderia_pseudomallei_1710b_uid58391 | 76810140 | YP_332316.1 | 0.56 | 0.611 | 0.368 | 0/18 | 0.267 | Unclassified |
| ATGC089 | Burkholderia_pseudomallei_668_uid58389 | 126440942 | YP_001057773.1 | 0.56 | 0.611 | 0.316 | 0/18 | 0.267 | Unclassified |
| ATGC089 | Burkholderia_pseudomallei_BPC006_uid174460 | 403517386 | YP_006651519.1 | 0.56 | 0.611 | 0.316 | 0/18 | 0.267 | Unclassified |
| ATGC089 | Burkholderia_pseudomallei_K96243_uid57733 | 53718327 | YP_107313.1 | 0.56 | 0.611 | 0.368 | 0/18 | 0.267 | Unclassified |
| ATGC089 | Burkholderia_pseudomallei_MSHR346_uid55259 | 237810925 | YP_002895376.1 | 0.56 | 0.611 | 0.316 | 0/18 | 0.267 | Unclassified |
| ATGC089 | Burkholderia_thailandensis_MSMB121_uid201037 | 488604835 | YP_007919570.1 | 0.6 | 0.611 | 0.316 | 0/18 | 0.267 | Unclassified |
| ATGC089 | Burkholderia_pseudomallei_668_uid58389 | 126440409 | YP_001057853.1 | 0.72 | 0.722 | 0.395 | 0/18 | 0.233 | Unclassified |
| ATGC089 | Burkholderia_thailandensis_MSMB121_uid201037 | 488604336 | YP_007919517.1 | 0.72 | 0.722 | 0.421 | 0/18 | 0.233 | Unclassified |
| ATGC090 | Burkholderia_383_uid58073 | 78065630 | YP_368399.1 | 1 | 1 | 0.579 | 0/18 | 0.233 | Sensitive |
| ATGC090 | Burkholderia_KJ006_uid165871 | 387901695 | YP_006332034.1 | 1 | 1 | 0.579 | 0/18 | 0.233 | Sensitive |
| ATGC090 | Burkholderia_ambifaria_AMMD_uid58303 | 115350975 | YP_772814.1 | 1 | 1 | 0.579 | 0/18 | 0.233 | Sensitive |
| ATGC090 | Burkholderia_ambifaria_MC40_6_uid58701 | 172059980 | YP_001807632.1 | 1 | 1 | 0.579 | 0/18 | 0.233 | Sensitive |
| ATGC090 | Burkholderia_cenocepacia_AU_1054_uid58371 | 107022123 | YP_620450.1 | 1 | 1 | 0.579 | 0/18 | 0.233 | Sensitive |
| ATGC090 | Burkholderia_cenocepacia_HI2424_uid58369 | 116689068 | YP_834691.1 | 1 | 1 | 0.579 | 0/18 | 0.233 | Sensitive |
| ATGC090 | Burkholderia_cenocepacia_J2315_uid57953 | 206561287 | YP_002232052.1 | 1 | 1 | 0.579 | 0/18 | 0.233 | Sensitive |
| ATGC090 | Burkholderia_cenocepacia_MC0_3_uid58769 | 170732356 | YP_001764303.1 | 1 | 1 | 0.579 | 0/18 | 0.233 | Sensitive |
| ATGC090 | Burkholderia_cepacia_GG4_uid173858 | 402567231 | YP_006616576.1 | 1 | 1 | 0.579 | 0/18 | 0.233 | Sensitive |
| ATGC090 | Burkholderia_multivorans_ATCC_17616_uid58697 | 161525429 | YP_001580441.1 | 1 | 1 | 0.579 | 0/18 | 0.233 | Sensitive |

| | | | | | | | | | |
|---|---|---|---|---|---|---|---|---|---|
| ATGC090 | Burkholderia_multivorans_ATCC_17616_uid58909 | 189349834 | YP_001945462.1 | 1 | 1 | 0.579 | 0/18 | 0.233 | Sensitive |
| ATGC090 | Burkholderia_vietnamiensis_G4_uid58075 | 134295077 | YP_001118812.1 | 1 | 1 | 0.579 | 0/18 | 0.233 | Sensitive |
| ATGC090 | Burkholderia_KJ006_uid165871 | 387905030 | YP_006335368.1 | 0.72 | 0.722 | 0.421 | 0/18 | 0.2 | Unclassified |
| ATGC090 | Burkholderia_vietnamiensis_G4_uid58075 | 134293620 | YP_001117356.1 | 0.72 | 0.722 | 0.421 | 0/18 | 0.2 | Unclassified |
| ATGC094 | Sulfolobus_islandicus_HVE10_4_uid162067 | 385773764 | YP_005646331.1 | 0.92 | 1 | 0.5 | 0/18 | 0.367 | Sensitive |
| ATGC094 | Sulfolobus_islandicus_LAL14_1_uid197216 | 479326079 | YP_007866134.1 | 0.92 | 1 | 0.5 | 0/18 | 0.367 | Sensitive |
| ATGC094 | Sulfolobus_islandicus_L_D_8_5_uid43679 | 284998307 | YP_003420075.1 | 0.92 | 1 | 0.5 | 0/18 | 0.367 | Sensitive |
| ATGC094 | Sulfolobus_islandicus_L_S_2_15_uid58871 | 227830793 | YP_002832573.1 | 0.92 | 1 | 0.5 | 0/18 | 0.367 | Sensitive |
| ATGC094 | Sulfolobus_islandicus_M_14_25_uid58849 | 227828056 | YP_002829836.1 | 0.92 | 1 | 0.5 | 0/18 | 0.367 | Sensitive |
| ATGC094 | Sulfolobus_islandicus_M_16_27_uid58851 | 229585325 | YP_002843827.1 | 0.92 | 1 | 0.5 | 0/18 | 0.367 | Sensitive |
| ATGC094 | Sulfolobus_islandicus_M_16_4_uid58841 | 238620286 | YP_002915112.1 | 0.92 | 1 | 0.5 | 0/18 | 0.367 | Sensitive |
| ATGC094 | Sulfolobus_islandicus_REY15A_uid162071 | 385776399 | YP_005648967.1 | 0.92 | 1 | 0.5 | 0/18 | 0.367 | Sensitive |
| ATGC094 | Sulfolobus_islandicus_Y_G_57_14_uid58923 | 229579689 | YP_002838088.1 | 0.92 | 1 | 0.5 | 0/18 | 0.367 | Sensitive |
| ATGC094 | Sulfolobus_islandicus_Y_N_15_51_uid58825 | 229581645 | YP_002840044.1 | 0.92 | 1 | 0.5 | 0/18 | 0.367 | Sensitive |
| ATGC094 | Sulfolobus_solfataricus_98_2_uid167998 | 384433763 | YP_005643121.1 | 0.88 | 1 | 0.526 | 0/18 | 0.433 | Sensitive |
| ATGC094 | Sulfolobus_solfataricus_P2_uid57721 | 15897251 | NP_341856.1 | 0.88 | 1 | 0.526 | 0/18 | 0.433 | Sensitive |
| ATGC105 | Bifidobacterium_breve_ACS_071_V_Sch8b_uid158863 | 384197175 | YP_005582919.1 | 1 | 0.889 | 0.553 | 0/18 | 0.333 | Sensitive |
| ATGC105 | Bifidobacterium_breve_UCC2003_uid193702 | 476417889 | YP_007554587.1 | 1 | 0.889 | 0.553 | 0/18 | 0.333 | Sensitive |
| ATGC105 | Bifidobacterium_longum_BBMN68_uid60163 | 312132994 | YP_004000333.1 | 1 | 0.889 | 0.553 | 0/18 | 0.333 | Sensitive |
| ATGC105 | Bifidobacterium_longum_DJO10A_uid58833 | 189439586 | YP_001954667.1 | 1 | 0.889 | 0.553 | 0/18 | 0.333 | Sensitive |
| ATGC105 | Bifidobacterium_longum_JCM_1217_uid62695 | 322690832 | YP_004220402.1 | 1 | 0.889 | 0.553 | 0/18 | 0.333 | Sensitive |

| | | | | | | | | | |
|---|---|---|---|---|---|---|---|---|---|
| ATGC105 | Bifidobacterium_longum_JDM301_uid49131 | 296453911 | YP_003661054.1 | 1 | 0.889 | 0.553 | 0/18 | 0.333 | Sensitive |
| ATGC105 | Bifidobacterium_longum_KACC_91563_uid158861 | 384201791 | YP_005587538.1 | 1 | 0.889 | 0.553 | 0/18 | 0.333 | Sensitive |
| ATGC105 | Bifidobacterium_longum_NCC2705_uid57939 | 23465541 | NP_696144.1 | 1 | 0.889 | 0.553 | 0/18 | 0.333 | Sensitive |
| ATGC105 | Bifidobacterium_longum_infantis_157F_uid62693 | 322688850 | YP_004208584.1 | 1 | 0.889 | 0.553 | 0/18 | 0.333 | Sensitive |
| ATGC105 | Bifidobacterium_longum_infantis_ATCC_15697_uid159865 | 384199800 | YP_005585543.1 | 1 | 0.889 | 0.526 | 0/18 | 0.333 | Sensitive |
| ATGC105 | Bifidobacterium_longum_infantis_ATCC_15697_uid58677 | 213692599 | YP_002323185.1 | 1 | 0.889 | 0.526 | 0/18 | 0.333 | Sensitive |
| ATGC106 | Bifidobacterium_animalis_ATCC_25527_uid162513 | 386867140 | YP_006280134.1 | 1 | 0.889 | 0.553 | 0/18 | 0.333 | Sensitive |
| ATGC106 | Bifidobacterium_animalis_lactis_AD011_uid58911 | 219683961 | YP_002470344.1 | 1 | 0.889 | 0.553 | 0/18 | 0.333 | Sensitive |
| ATGC106 | Bifidobacterium_animalis_lactis_B420_uid163691 | 387820869 | YP_006300912.1 | 1 | 0.889 | 0.553 | 0/18 | 0.333 | Sensitive |
| ATGC106 | Bifidobacterium_animalis_lactis_BB_12_uid158871 | 384191252 | YP_005577000.1 | 1 | 0.889 | 0.553 | 0/18 | 0.333 | Sensitive |
| ATGC106 | Bifidobacterium_animalis_lactis_BLC1_uid158867 | 384193995 | YP_005579741.1 | 1 | 0.889 | 0.553 | 0/18 | 0.333 | Sensitive |
| ATGC106 | Bifidobacterium_animalis_lactis_Bi_07_uid163693 | 387822544 | YP_006302493.1 | 1 | 0.889 | 0.553 | 0/18 | 0.333 | Sensitive |
| ATGC106 | Bifidobacterium_animalis_lactis_Bl_04_uid59359 | 241191003 | YP_002968397.1 | 1 | 0.889 | 0.553 | 0/18 | 0.333 | Sensitive |
| ATGC106 | Bifidobacterium_animalis_lactis_CNCM_I_2494_uid158869 | 384192399 | YP_005578146.1 | 1 | 0.889 | 0.553 | 0/18 | 0.333 | Sensitive |
| ATGC106 | Bifidobacterium_animalis_lactis_DSM_10140_uid59357 | 241196409 | YP_002969964.1 | 1 | 0.889 | 0.553 | 0/18 | 0.333 | Sensitive |
| ATGC106 | Bifidobacterium_animalis_lactis_V9_uid158865 | 384195561 | YP_005581306.1 | 1 | 0.889 | 0.553 | 0/18 | 0.333 | Sensitive |
| ATGC109 | Listeria_innocua_Clip11262_uid61567 | 16801103 | NP_471371.1 | 0.8 | 0.833 | 1 | 0/18 | 0.2 | Resistant |
| ATGC109 | Listeria_ivanovii_PAM_55_uid73473 | 347549322 | YP_004855650.1 | 0.8 | 0.833 | 1 | 0/18 | 0.2 | Resistant |
| ATGC109 | Listeria_monocytogenes_07PF0776_uid162185 | 386732664 | YP_006206160.1 | 0.8 | 0.833 | 1 | 0/18 | 0.2 | Resistant |
| ATGC109 | Listeria_monocytogenes_08_5923_uid43727 | 284995510 | YP_003417278.1 | 0.8 | 0.833 | 1 | 0/18 | 0.2 | Resistant |
| ATGC109 | Listeria_monocytogenes_10403S_uid54461 | 386044231 | YP_005963036.1 | 0.8 | 0.833 | 1 | 0/18 | 0.2 | Resistant |

| | | | | | | | | | |
|---|---|---|---|---|---|---|---|---|---|
| ATGC109 | Listeria_monocytogenes_ATCC_19117_uid175109 | 405750276 | YP_006673742.1 | 0.8 | 0.833 | 1 | 0/18 | 0.2 | Resistant |
| ATGC109 | Listeria_monocytogenes_Clip80459_uid59317 | 226224527 | YP_002758634.1 | 0.8 | 0.833 | 1 | 0/18 | 0.2 | Resistant |
| ATGC109 | Listeria_monocytogenes_EGD_e_uid61583 | 16803962 | NP_465447.1 | 0.8 | 0.833 | 1 | 0/18 | 0.2 | Resistant |
| ATGC109 | Listeria_monocytogenes_FSL_R2_561_uid54441 | 386050899 | YP_005968890.1 | 0.8 | 0.833 | 1 | 0/18 | 0.2 | Resistant |
| ATGC109 | Listeria_monocytogenes_Finland_1998_uid54443 | 386054178 | YP_005971736.1 | 0.8 | 0.833 | 1 | 0/18 | 0.2 | Resistant |
| ATGC109 | Listeria_monocytogenes_HCC23_uid59203 | 217963925 | YP_002349603.1 | 0.8 | 0.833 | 1 | 0/18 | 0.2 | Resistant |
| ATGC109 | Listeria_monocytogenes_J0161_uid54459 | 386047575 | YP_005965907.1 | 0.8 | 0.833 | 1 | 0/18 | 0.2 | Resistant |
| ATGC109 | Listeria_monocytogenes_L312_uid175768 | 406704708 | YP_006755062.1 | 0.8 | 0.833 | 1 | 0/18 | 0.2 | Resistant |
| ATGC109 | Listeria_monocytogenes_La111_uid193768 | 470207793 | YP_007604368.1 | 0.16 | 0.167 | 0.158 | 0/18 | 0.1 | Unclassified |
| ATGC109 | Listeria_monocytogenes_M7_uid162131 | 386027303 | YP_005948079.1 | 0.8 | 0.833 | 1 | 0/18 | 0.2 | Resistant |
| ATGC109 | Listeria_monocytogenes_N53_1_uid193767 | 470210925 | YP_007607499.1 | 0.16 | 0.167 | 0.158 | 0/18 | 0.033 | Unclassified |
| ATGC109 | Listeria_monocytogenes_SLCC2372_uid174872 | 404284419 | YP_006685316.1 | 0.8 | 0.833 | 1 | 0/18 | 0.2 | Resistant |
| ATGC109 | Listeria_monocytogenes_SLCC2376_uid175111 | 404408368 | YP_006691083.1 | 0.8 | 0.833 | 1 | 0/18 | 0.2 | Resistant |
| ATGC109 | Listeria_monocytogenes_SLCC2378_uid175105 | 405753150 | YP_006676615.1 | 0.8 | 0.833 | 1 | 0/18 | 0.2 | Resistant |
| ATGC109 | Listeria_monocytogenes_SLCC2479_uid175108 | 405758973 | YP_006688249.1 | 0.8 | 0.833 | 1 | 0/18 | 0.2 | Resistant |
| ATGC109 | Listeria_monocytogenes_SLCC2540_uid175106 | 405756083 | YP_006679547.1 | 0.8 | 0.833 | 1 | 0/18 | 0.2 | Resistant |
| ATGC109 | Listeria_monocytogenes_SLCC5850_uid175110 | 404411224 | YP_006696812.1 | 0.8 | 0.833 | 1 | 0/18 | 0.2 | Resistant |
| ATGC109 | Listeria_monocytogenes_SLCC7179_uid175107 | 404414001 | YP_006699588.1 | 0.8 | 0.833 | 1 | 0/18 | 0.2 | Resistant |
| ATGC109 | Listeria_monocytogenes_serotype_1_2b_SLCC2755_uid52455 | 404281535 | YP_006682433.1 | 0.8 | 0.833 | 1 | 0/18 | 0.2 | Resistant |
| ATGC109 | Listeria_monocytogenes_serotype_4a_L99_uid161953 | 386008695 | YP_005926973.1 | 0.8 | 0.833 | 1 | 0/18 | 0.2 | Resistant |
| ATGC109 | Listeria_monocytogenes_serotype_4b_F2365_uid57689 | 46908156 | YP_014545.1 | 0.8 | 0.833 | 1 | 0/18 | 0.2 | Resistant |

| | | | | | | | | | |
|---|---|---|---|---|---|---|---|---|---|
| ATGC109 | Listeria_monocytogenes_serotype_4b_LL195_uid182103 | 424714798 | YP_007015513.1 | 0.8 | 0.833 | 1 | 0/18 | 0.2 | Resistant |
| ATGC109 | Listeria_monocytogenes_serotype_7_SLCC2482_uid174871 | 404287351 | YP_006693937.1 | 0.8 | 0.833 | 1 | 0/18 | 0.2 | Resistant |
| ATGC109 | Listeria_monocytogenes_uid43671 | 284802368 | YP_003414233.1 | 0.8 | 0.833 | 1 | 0/18 | 0.2 | Resistant |
| ATGC109 | Listeria_seeligeri_serovar_1_2b_SLCC3954_uid46215 | 289435272 | YP_003465144.1 | 0.8 | 0.833 | 1 | 0/18 | 0.2 | Resistant |
| ATGC109 | Listeria_welshimeri_serovar_6b_SLCC5334_uid61605 | 116873365 | YP_850146.1 | 0.8 | 0.833 | 1 | 0/18 | 0.2 | Resistant |
| ATGC121 | Shewanella_ANA_3_uid58347 | 117920420 | YP_869612.1 | 1 | 1 | 0.579 | 0/18 | 0.233 | Sensitive |
| ATGC121 | Shewanella_MR_4_uid58345 | 113970260 | YP_734053.1 | 1 | 1 | 0.579 | 0/18 | 0.233 | Sensitive |
| ATGC121 | Shewanella_MR_7_uid58343 | 114047551 | YP_738101.1 | 1 | 1 | 0.579 | 0/18 | 0.233 | Sensitive |
| ATGC121 | Shewanella_W3_18_1_uid58341 | 120598783 | YP_963357.1 | 1 | 1 | 0.579 | 0/18 | 0.233 | Sensitive |
| ATGC121 | Shewanella_baltica_BA175_uid52601 | 386324579 | YP_006020696.1 | 1 | 1 | 0.579 | 0/18 | 0.233 | Sensitive |
| ATGC121 | Shewanella_baltica_OS117_uid162025 | 386341018 | YP_006037384.1 | 1 | 1 | 0.579 | 0/18 | 0.233 | Sensitive |
| ATGC121 | Shewanella_baltica_OS155_uid58259 | 126174289 | YP_001050438.1 | 1 | 1 | 0.579 | 0/18 | 0.233 | Sensitive |
| ATGC121 | Shewanella_baltica_OS185_uid58743 | 153000801 | YP_001366482.1 | 1 | 1 | 0.579 | 0/18 | 0.233 | Sensitive |
| ATGC121 | Shewanella_baltica_OS195_uid58261 | 160875510 | YP_001554826.1 | 1 | 1 | 0.553 | 0/18 | 0.233 | Sensitive |
| ATGC121 | Shewanella_baltica_OS223_uid58775 | 217973240 | YP_002357991.1 | 1 | 1 | 0.579 | 0/18 | 0.233 | Sensitive |
| ATGC121 | Shewanella_baltica_OS678_uid50553 | 378708708 | YP_005273602.1 | 1 | 1 | 0.553 | 0/18 | 0.233 | Sensitive |
| ATGC121 | Shewanella_oneidensis_MR_1_uid57949 | 24373951 | NP_717994.1 | 1 | 1 | 0.579 | 0/18 | 0.233 | Sensitive |
| ATGC121 | Shewanella_putrefaciens_200_uid161927 | 386313723 | YP_006009888.1 | 1 | 1 | 0.579 | 0/18 | 0.233 | Sensitive |
| ATGC121 | Shewanella_putrefaciens_CN_32_uid58267 | 146293140 | YP_001183564.1 | 1 | 1 | 0.579 | 0/18 | 0.233 | Sensitive |
| ATGC128 | Yersinia_enterocolitica_8081_uid57741 | 123441850 | YP_001005833.1 | 1 | 1 | 0.553 | 0/18 | 0.233 | Sensitive |
| ATGC128 | Yersinia_enterocolitica_palearctica_105_5R_r__uid63663 | 332162208 | YP_004298785.1 | 1 | 1 | 0.553 | 0/18 | 0.233 | Sensitive |

| | | | | | | | | | |
|---|---|---|---|---|---|---|---|---|---|
| ATGC128 | Yersinia_enterocolitica_palearctica_Y11_uid162069 | 386307862 | YP_006003918.1 | 1 | 1 | 0.553 | 0/18 | 0.233 | Sensitive |
| ATGC128 | Yersinia_pestis_A1122_uid158119 | 384140747 | YP_005523449.1 | 1 | 1 | 0.579 | 0/18 | 0.233 | Sensitive |
| ATGC128 | Yersinia_pestis_Angola_uid58485 | 162418202 | YP_001606427.1 | 1 | 1 | 0.579 | 0/18 | 0.233 | Sensitive |
| ATGC128 | Yersinia_pestis_Antiqua_uid58607 | 108806678 | YP_650594.1 | 1 | 1 | 0.579 | 0/18 | 0.233 | Sensitive |
| ATGC128 | Yersinia_pestis_CO92_uid57621 | 218928538 | YP_002346413.1 | 1 | 1 | 0.579 | 0/18 | 0.233 | Sensitive |
| ATGC128 | Yersinia_pestis_D106004_uid158071 | 384121825 | YP_005504445.1 | 1 | 1 | 0.579 | 0/18 | 0.233 | Sensitive |
| ATGC128 | Yersinia_pestis_D182038_uid158073 | 384125379 | YP_005507993.1 | 1 | 1 | 0.579 | 0/18 | 0.233 | Sensitive |
| ATGC128 | Yersinia_pestis_KIM_10_uid57875 | 22126661 | NP_670084.1 | 1 | 1 | 0.579 | 0/18 | 0.233 | Sensitive |
| ATGC128 | Yersinia_pestis_Nepal516_uid58609 | 108812748 | YP_648515.1 | 1 | 1 | 0.579 | 0/18 | 0.233 | Sensitive |
| ATGC128 | Yersinia_pestis_Pestoides_F_uid58619 | 145599577 | YP_001163653.1 | 1 | 1 | 0.579 | 0/18 | 0.233 | Sensitive |
| ATGC128 | Yersinia_pestis_Z176003_uid47317 | 294503380 | YP_003567442.1 | 1 | 1 | 0.579 | 0/18 | 0.233 | Sensitive |
| ATGC128 | Yersinia_pestis_biovar_Medievalis_Harbin_35_uid158537 | 384415247 | YP_005624609.1 | 1 | 1 | 0.579 | 0/18 | 0.233 | Sensitive |
| ATGC128 | Yersinia_pestis_biovar_Microtus_91001_uid58037 | 45441030 | NP_992569.1 | 1 | 1 | 0.579 | 0/18 | 0.233 | Sensitive |
| ATGC128 | Yersinia_pseudotuberculosis_IP_31758_uid58487 | 153949635 | YP_001401547.1 | 1 | 1 | 0.579 | 0/18 | 0.233 | Sensitive |
| ATGC128 | Yersinia_pseudotuberculosis_IP_32953_uid58157 | 51595755 | YP_069946.1 | 1 | 1 | 0.579 | 0/18 | 0.233 | Sensitive |
| ATGC128 | Yersinia_pseudotuberculosis_PB1__uid59153 | 186894833 | YP_001871945.1 | 1 | 1 | 0.579 | 0/18 | 0.233 | Sensitive |
| ATGC128 | Yersinia_pseudotuberculosis_YPIII_uid59151 | 170024895 | YP_001721400.1 | 1 | 1 | 0.579 | 0/18 | 0.233 | Sensitive |
| ATGC135 | Xanthomonas_axonopodis_Xac29_1_uid193774 | 470471539 | YP_007636107.1 | 0.8 | 0.889 | 1 | 0/18 | 0.3 | Resistant |
| ATGC135 | Xanthomonas_axonopodis_citri_306_uid57889 | 21242400 | NP_641982.1 | 0.8 | 0.889 | 1 | 0/18 | 0.3 | Resistant |
| ATGC135 | Xanthomonas_axonopodis_citrumelo_F1_uid73179 | 346724530 | YP_004851199.1 | 0.8 | 0.889 | 1 | 0/18 | 0.3 | Resistant |
| ATGC135 | Xanthomonas_campestris_8004_uid57595 | 66768950 | YP_243712.1 | 0.8 | 0.889 | 1 | 0/18 | 0.267 | Resistant |

| | | | | | | | | | |
|---|---|---|---|---|---|---|---|---|---|
| ATGC135 | Xanthomonas_campestris_ATCC_33913_uid57887 | 21231045 | NP_636962.1 | 0.8 | 0.889 | 1 | 0/18 | 0.267 | Resistant |
| ATGC135 | Xanthomonas_campestris_raphani_756C_uid159539 | 384427525 | YP_005636884.1 | 0.8 | 0.889 | 1 | 0/18 | 0.267 | Resistant |
| ATGC135 | Xanthomonas_campestris_uid61643 | 188992064 | YP_001904074.1 | 0.8 | 0.889 | 1 | 0/18 | 0.267 | Resistant |
| ATGC135 | Xanthomonas_campestris_vesicatoria_85_10_uid58321 | 78047247 | YP_363422.1 | 0.8 | 0.889 | 1 | 0/18 | 0.3 | Resistant |
| ATGC135 | Xanthomonas_citri_Aw12879_uid194444 | 471268184 | YP_007650649.1 | 0.8 | 0.889 | 1 | 0/18 | 0.3 | Resistant |
| ATGC135 | Xanthomonas_oryzae_KACC_10331_uid58155 | 58582009 | YP_201025.1 | 0.76 | 0.833 | 1 | 0/18 | 0.3 | Resistant |
| ATGC135 | Xanthomonas_oryzae_MAFF_311018_uid58547 | 84623923 | YP_451295.1 | 0.76 | 0.833 | 1 | 0/18 | 0.3 | Resistant |
| ATGC135 | Xanthomonas_oryzae_PXO99A_uid59131 | 188576380 | YP_001913309.1 | 0.76 | 0.833 | 1 | 0/18 | 0.3 | Resistant |
| ATGC135 | Xanthomonas_oryzae_oryzicola_BLS256_uid54411 | 384419788 | YP_005629148.1 | 0.8 | 0.889 | 1 | 0/18 | 0.3 | Resistant |
| ATGC137 | Brucella_abortus_A13334_uid83615 | 376272066 | YP_005150644.1 | 0.76 | 0.833 | 1 | 0/18 | 0.233 | Resistant |
| ATGC137 | Brucella_abortus_S19_uid58873 | 189023290 | YP_001934058.1 | 0.76 | 0.833 | 1 | 0/18 | 0.233 | Resistant |
| ATGC137 | Brucella_abortus_bv__1_9_941_uid58019 | 62289014 | YP_220807.1 | 0.76 | 0.833 | 1 | 0/18 | 0.233 | Resistant |
| ATGC137 | Brucella_canis_ATCC_23365_uid59009 | 161618015 | YP_001591902.1 | 0.76 | 0.833 | 1 | 0/18 | 0.233 | Resistant |
| ATGC137 | Brucella_canis_HSK_A52141_uid83613 | 376275204 | YP_005115643.1 | 0.76 | 0.833 | 1 | 0/18 | 0.233 | Resistant |
| ATGC137 | Brucella_melitensis_ATCC_23457_uid59241 | 225851569 | YP_002731802.1 | 0.76 | 0.833 | 1 | 0/18 | 0.233 | Resistant |
| ATGC137 | Brucella_melitensis_M28_uid158857 | 384407490 | YP_005596111.1 | 0.76 | 0.833 | 1 | 0/18 | 0.233 | Resistant |
| ATGC137 | Brucella_melitensis_M5_90_uid158855 | 384210392 | YP_005599474.1 | 0.76 | 0.833 | 1 | 0/18 | 0.233 | Resistant |
| ATGC137 | Brucella_melitensis_NI_uid158853 | 384444114 | YP_005602833.1 | 0.76 | 0.833 | 1 | 0/18 | 0.233 | Resistant |
| ATGC137 | Brucella_melitensis_biovar_Abortus_2308_uid62937 | 82698952 | YP_413526.1 | 0.76 | 0.833 | 1 | 0/18 | 0.233 | Resistant |
| ATGC137 | Brucella_melitensis_bv__1_16M_uid57735 | 17988200 | NP_540834.1 | 0.76 | 0.833 | 1 | 0/18 | 0.233 | Resistant |
| ATGC137 | Brucella_microti_CCM_4915_uid59319 | 256368490 | YP_003105996.1 | 0.76 | 0.833 | 1 | 0/18 | 0.233 | Resistant |

| | | | | | | | | | |
|---|---|---|---|---|---|---|---|---|---|
| ATGC137 | Brucella_ovis_ATCC_25840_uid58113 | 148559649 | YP_001258071.1 | 0.76 | 0.833 | 1 | 0/18 | 0.233 | Resistant |
| ATGC137 | Brucella_pinnipedialis_B2_94_uid71131 | 340789655 | YP_004755119.1 | 0.76 | 0.833 | 1 | 0/18 | 0.233 | Resistant |
| ATGC137 | Brucella_suis_1330_uid159871 | 384223722 | YP_005614886.1 | 0.76 | 0.833 | 1 | 0/18 | 0.233 | Resistant |
| ATGC137 | Brucella_suis_1330_uid57927 | 23500940 | NP_697067.1 | 0.76 | 0.833 | 1 | 0/18 | 0.233 | Resistant |
| ATGC137 | Brucella_suis_ATCC_23445_uid59015 | 163842301 | YP_001626705.1 | 0.76 | 0.833 | 1 | 0/18 | 0.233 | Resistant |
| ATGC137 | Brucella_suis_VBI22_uid83617 | 376279728 | YP_005153734.1 | 0.76 | 0.833 | 1 | 0/18 | 0.233 | Resistant |
| ATGC137 | Ochrobactrum_anthropi_ATCC_49188_uid58921 | 153007376 | YP_001368591.1 | 0.76 | 0.833 | 1 | 0/18 | 0.233 | Resistant |
| ATGC138 | Neisseria_gonorrhoeae_FA_1090_uid57611 | 59801298 | YP_208010.1 | 1 | 1 | 0.605 | 0/18 | 0.267 | Sensitive |
| ATGC138 | Neisseria_gonorrhoeae_NCCP11945_uid59191 | 194098468 | YP_002001528.1 | 1 | 1 | 0.605 | 0/18 | 0.267 | Sensitive |
| ATGC138 | Neisseria_gonorrhoeae_TCDC_NG08107_uid161097 | 385335607 | YP_005889554.1 | 1 | 1 | 0.605 | 0/18 | 0.267 | Sensitive |
| ATGC138 | Neisseria_lactamica_020_06_uid60851 | 313668544 | YP_004048828.1 | 1 | 1 | 0.579 | 0/18 | 0.267 | Sensitive |
| ATGC138 | Neisseria_meningitidis_053442_uid58587 | 161870300 | YP_001599470.1 | 0.84 | 0.833 | 0.5 | 0/18 | 0.2 | Unclassified |
| ATGC138 | Neisseria_meningitidis_8013_uid161967 | 385323920 | YP_005878359.1 | 1 | 1 | 0.579 | 0/18 | 0.267 | Sensitive |
| ATGC138 | Neisseria_meningitidis_FAM18_uid57825 | 121635120 | YP_975365.1 | 1 | 1 | 0.579 | 0/18 | 0.267 | Sensitive |
| ATGC138 | Neisseria_meningitidis_G2136_uid162085 | 385340321 | YP_005894193.1 | 1 | 1 | 0.579 | 0/18 | 0.267 | Sensitive |
| ATGC138 | Neisseria_meningitidis_H44_76_uid162083 | 385852949 | YP_005899463.1 | 1 | 1 | 0.579 | 0/18 | 0.267 | Sensitive |
| ATGC138 | Neisseria_meningitidis_M01_240149_uid162079 | 385341665 | YP_005895536.1 | 1 | 1 | 0.579 | 0/18 | 0.267 | Sensitive |
| ATGC138 | Neisseria_meningitidis_M01_240355_uid162075 | 385855477 | YP_005901990.1 | 1 | 1 | 0.579 | 0/18 | 0.267 | Sensitive |
| ATGC138 | Neisseria_meningitidis_M04_240196_uid162081 | 385850992 | YP_005897507.1 | 1 | 1 | 0.553 | 0/18 | 0.267 | Sensitive |
| ATGC138 | Neisseria_meningitidis_MC58_uid57817 | 15677291 | NP_274444.1 | 1 | 1 | 0.579 | 0/18 | 0.267 | Sensitive |
| ATGC138 | Neisseria_meningitidis_NZ_05_33_uid162077 | 385857488 | YP_005904000.1 | 1 | 1 | 0.553 | 0/18 | 0.267 | Sensitive |

| | | | | | | | | | |
|---|---|---|---|---|---|---|---|---|---|
| ATGC138 | Neisseria_meningitidis_WUE_2594_uid162093 | 385338265 | YP_005892138.1 | 1 | 1 | 0.579 | 0/18 | 0.267 | Sensitive |
| ATGC138 | Neisseria_meningitidis_Z2491_uid57819 | 218768435 | YP_002342947.1 | 1 | 1 | 0.579 | 0/18 | 0.267 | Sensitive |
| ATGC138 | Neisseria_meningitidis_alpha14_uid61649 | 254805212 | YP_003083433.1 | 1 | 1 | 0.553 | 0/18 | 0.267 | Sensitive |
| ATGC138 | Neisseria_meningitidis_alpha710_uid161971 | 385328684 | YP_005882987.1 | 1 | 1 | 0.553 | 0/18 | 0.267 | Sensitive |
| ATGC139 | Francisella_TX077308_uid68321 | 337755608 | YP_004648119.1 | 1 | 0.944 | 0.526 | 0/18 | 0.3 | Sensitive |
| ATGC139 | Francisella_cf__novicida_3523_uid162107 | 387824367 | YP_005823838.1 | 1 | 0.944 | 0.526 | 0/18 | 0.3 | Sensitive |
| ATGC139 | Francisella_cf__novicida_Fx1_uid162105 | 385793069 | YP_005826045.1 | 1 | 0.944 | 0.553 | 0/18 | 0.3 | Sensitive |
| ATGC139 | Francisella_noatunensis_orientalis_Toba_04_uid164779 | 387886901 | YP_006317200.1 | 1 | 0.944 | 0.526 | 0/18 | 0.3 | Sensitive |
| ATGC139 | Francisella_novicida_U112_uid58499 | 118497681 | YP_898731.1 | 1 | 0.944 | 0.553 | 0/18 | 0.3 | Sensitive |
| ATGC139 | Francisella_philomiragia_ATCC_25017_uid59105 | 167627743 | YP_001678243.1 | 1 | 0.944 | 0.526 | 0/18 | 0.3 | Sensitive |
| ATGC139 | Francisella_tularensis_FSC198_uid58693 | 110670186 | YP_666743.1 | 1 | 0.944 | 0.553 | 0/18 | 0.3 | Sensitive |
| ATGC139 | Francisella_tularensis_NE061598_uid161973 | 385794348 | YP_005830754.1 | 1 | 0.944 | 0.553 | 0/18 | 0.3 | Sensitive |
| ATGC139 | Francisella_tularensis_SCHU_S4_uid57589 | 56707715 | YP_169611.1 | 1 | 0.944 | 0.553 | 0/18 | 0.3 | Sensitive |
| ATGC139 | Francisella_tularensis_TI0902_uid89373 | 379725570 | YP_005317756.1 | 1 | 0.944 | 0.553 | 0/18 | 0.3 | Sensitive |
| ATGC139 | Francisella_tularensis_TIGB03_uid89379 | 379716966 | YP_005305302.1 | 1 | 0.944 | 0.553 | 0/18 | 0.3 | Sensitive |
| ATGC139 | Francisella_tularensis_WY96_3418_uid58811 | 134302101 | YP_001122070.1 | 1 | 0.944 | 0.553 | 0/18 | 0.3 | Sensitive |
| ATGC139 | Francisella_tularensis_holarctica_F92_uid181998 | 423050560 | YP_007008994.1 | 1 | 0.944 | 0.553 | 0/18 | 0.3 | Sensitive |
| ATGC139 | Francisella_tularensis_holarctica_FSC200_uid54341 | 422938633 | YP_007011780.1 | 1 | 0.944 | 0.553 | 0/18 | 0.3 | Sensitive |
| ATGC139 | Francisella_tularensis_holarctica_FTNF002_00_uid58999 | 156502271 | YP_001428336.1 | 1 | 0.944 | 0.553 | 0/18 | 0.3 | Sensitive |
| ATGC139 | Francisella_tularensis_holarctica_LVS_uid58595 | 89256212 | YP_513574.1 | 1 | 0.944 | 0.553 | 0/18 | 0.3 | Sensitive |
| ATGC139 | Francisella_tularensis_holarctica_OSU18_uid58687 | 115314679 | YP_763402.1 | 1 | 0.944 | 0.553 | 0/18 | 0.3 | Sensitive |

| | | | | | | | | | |
|---|---|---|---|---|---|---|---|---|---|
| ATGC139 | Francisella_tularensis_mediasiatica_FSC147_uid58939 | 187931440 | YP_001891424.1 | 1 | 0.944 | 0.553 | 0/18 | 0.3 | Sensitive |
| ATGC144 | Campylobacter_jejuni_81116_uid58771 | 157415152 | YP_001482408.1 | 0.76 | 0.833 | 1 | 0/18 | 0.167 | Resistant |
| ATGC144 | Campylobacter_jejuni_81_176_uid58503 | 121613725 | YP_001000570.1 | 0.76 | 0.833 | 1 | 0/18 | 0.167 | Resistant |
| ATGC144 | Campylobacter_jejuni_IA3902_uid159531 | 384448148 | YP_005656199.1 | 0.76 | 0.833 | 1 | 0/18 | 0.167 | Resistant |
| ATGC144 | Campylobacter_jejuni_ICDCCJ07001_uid61249 | 315124391 | YP_004066395.1 | 0.32 | 0.5 | 0.342 | 0/18 | 0 | Unclassified |
| ATGC144 | Campylobacter_jejuni_ICDCCJ07001_uid61249 | 315124392 | YP_004066396.1 | 0.44 | 0.333 | 0.658 | 0/18 | 0.167 | Unclassified |
| ATGC144 | Campylobacter_jejuni_M1_uid159535 | 384441512 | YP_005657815.1 | 0.76 | 0.833 | 1 | 0/18 | 0.167 | Resistant |
| ATGC144 | Campylobacter_jejuni_NCTC_11168_BN148_uid174152 | 403055638 | YP_006633043.1 | 0.76 | 0.833 | 1 | 0/18 | 0.167 | Resistant |
| ATGC144 | Campylobacter_jejuni_NCTC_11168___ATCC_700819_uid57587 | 218562515 | YP_002344294.1 | 0.76 | 0.833 | 1 | 0/18 | 0.167 | Resistant |
| ATGC144 | Campylobacter_jejuni_PT14_uid176499 | 407942292 | YP_006857934.1 | 0.76 | 0.833 | 1 | 0/18 | 0.167 | Resistant |
| ATGC144 | Campylobacter_jejuni_RM1221_uid57899 | 57237724 | YP_178972.1 | 0.76 | 0.833 | 1 | 0/18 | 0.167 | Resistant |
| ATGC144 | Campylobacter_jejuni_S3_uid159533 | 384443248 | YP_005659500.1 | 0.76 | 0.833 | 1 | 0/18 | 0.167 | Resistant |
| ATGC144 | Campylobacter_jejuni_doylei_269_97_uid58671 | 153951777 | YP_001398032.1 | 0.76 | 0.833 | 1 | 0/18 | 0.167 | Resistant |
| ATGC163 | Propionibacterium_acnes_266_uid162059 | 386024214 | YP_005942519.1 | 1 | 0.889 | 0.5 | 0/18 | 0.433 | Sensitive |
| ATGC163 | Propionibacterium_acnes_6609_uid162137 | 387503643 | YP_005944872.1 | 0.96 | 0.833 | 0.5 | 0/18 | 0.433 | Unclassified |
| ATGC163 | Propionibacterium_acnes_ATCC_11828_uid162177 | 386071234 | YP_005986130.1 | 1 | 0.889 | 0.5 | 0/18 | 0.433 | Sensitive |
| ATGC163 | Propionibacterium_acnes_C1_uid176501 | 407935666 | YP_006851308.1 | 1 | 0.889 | 0.5 | 0/18 | 0.433 | Sensitive |
| ATGC163 | Propionibacterium_acnes_HL096PA1_uid198524 | 482890069 | YP_007887232.1 | 1 | 0.889 | 0.5 | 0/18 | 0.433 | Sensitive |
| ATGC163 | Propionibacterium_acnes_KPA171202_uid58101 | 50842747 | YP_055974.1 | 1 | 0.889 | 0.5 | 0/18 | 0.433 | Sensitive |
| ATGC163 | Propionibacterium_acnes_SK137_uid48071 | 295130828 | YP_003581491.1 | 1 | 0.889 | 0.5 | 0/18 | 0.433 | Sensitive |
| ATGC163 | Propionibacterium_acnes_TypeIA2_P_acn17_uid80735 | 365965201 | YP_004946766.1 | 1 | 0.889 | 0.5 | 0/18 | 0.433 | Sensitive |

| | | | | | | | | | |
|---|---|---|---|---|---|---|---|---|---|
| ATGC163 | Propionibacterium_acnes_TypeIA2_P_acn31_uid80733 | 365962960 | YP_004944526.1 | 1 | 0.889 | 0.5 | 0/18 | 0.433 | Sensitive |
| ATGC163 | Propionibacterium_acnes_TypeIA2_P_acn33_uid80745 | 365974135 | YP_004955694.1 | 1 | 0.889 | 0.5 | 0/18 | 0.433 | Sensitive |
| ATGC163 | Propionibacterium_avidum_44067_uid197361 | 480329133 | YP_007870895.1 | 0.96 | 0.889 | 0.474 | 0/18 | 0.467 | Unclassified |
| ATGC186 | Legionella_pneumophila_2300_99_Alcoy_uid48801 | 296106999 | YP_003618699.1 | 0.76 | 0.833 | 1 | 0/18 | 0.233 | Resistant |
| ATGC186 | Legionella_pneumophila_ATCC_43290_uid86885 | 378777284 | YP_005185721.1 | 0.76 | 0.833 | 1 | 0/18 | 0.233 | Resistant |
| ATGC186 | Legionella_pneumophila_Corby_uid58733 | 148358950 | YP_001250157.1 | 0.76 | 0.833 | 1 | 0/18 | 0.233 | Resistant |
| ATGC186 | Legionella_pneumophila_HL06041035_uid170534 | 397667037 | YP_006508574.1 | 0.76 | 0.833 | 1 | 0/18 | 0.233 | Resistant |
| ATGC186 | Legionella_pneumophila_Lens_uid58209 | 54294305 | YP_126720.1 | 0.76 | 0.833 | 1 | 0/18 | 0.233 | Resistant |
| ATGC186 | Legionella_pneumophila_Lorraine_uid170535 | 397663851 | YP_006505389.1 | 0.76 | 0.833 | 1 | 0/18 | 0.233 | Resistant |
| ATGC186 | Legionella_pneumophila_Paris_uid58211 | 54297329 | YP_123698.1 | 0.76 | 0.833 | 1 | 0/18 | 0.233 | Resistant |
| ATGC186 | Legionella_pneumophila_Philadelphia_1_uid193710 | 470182452 | YP_007568327.1 | 0.76 | 0.833 | 1 | 0/18 | 0.233 | Resistant |
| ATGC186 | Legionella_pneumophila_Philadelphia_1_uid57609 | 52841649 | YP_095448.1 | 0.76 | 0.833 | 1 | 0/18 | 0.233 | Resistant |
| ATGC186 | Legionella_pneumophila_Thunder_Bay_uid206517 | 509150737 | YP_008063111.1 | 0.76 | 0.833 | 1 | 0/18 | 0.233 | Resistant |

**Supplementary table 3. Differential sensitivity to glyphosate by lifestyle and taxonomy.** Percentage of species sensitive to glyphosate.

| Life style | n | Class I | Class II | Class III | Class IV | Unknown | Chi-squared test |
|---|---|---|---|---|---|---|---|
| Intracellular parasite (P) | 64 | 73% | 0% | 0% | 0% | 27% | - |
| Facultative host-associated (FHA) | 336 | 68% | 16% | 0% | 0% | 16% | p-value < 0.001 |
| free-living (FL) | 329 | 21% | 66% | 0% | 0% | 12% | |
| **Taxonomy** | **n** | **Class I** | **Class II** | **Class III** | **Class IV** | **Unknown** | |
| Firmicutes (F) | 201 | 73% | 0% | 0% | 0% | 27% | p-value < 0.001 |
| Actinobacteria (A) | 93 | 68% | 16% | 0% | 0% | 16% | |
| Proteobacteria (P) | 359 | 21% | 66% | 0% | 0% | 12% | |

**Supplementary table 4. Differential sensitivity to glyphosate by lifestyle and taxonomy.** Relative number of sensitive bacteria to glyphosate

| Taxonomy | Facultative host-associate | Free-living |
|---|---|---|
| Firmicutes | - (n = 0) | 0 (n = 201) |
| Actinobacteria | 1.00 (n = 55) | 0.84 (n = 38) |
| Proteobacteria | 0.62 (n = 62) | 0.33 (n = 78) |

**Supplementary table 5.** Spearman correlation coefficients ($r^2$, p-value, n) between the sensitivity and a several genome parameters: total genome dynamics (GLER gene), genome number (genome no), genome size, protein coding genes, gene families, median gene cluster content (median GC), median dN/dS, synteny distance (dY) and aromatic amino acid % (% Arom) in different ATGCs (alignable tight genome clusters).

| | | Sensitivity | GLER Gene | Genome no. | Genome size | Prot coding genes | Gene Families | Median GC | Median dN/dS | dY | % Arom |
|---|---|---|---|---|---|---|---|---|---|---|---|
| **Sensitivity** | $r^2$ | 1 | -0.21 | -0.07 | -0.09 | -0.16 | -0.16 | 0.26 | 0.18 | 0.06 | -0.30 |
| | **p** | 0 | 0.240 | 0.690 | 0.622 | 0.379 | 0.390 | 0.148 | 0.356 | 0.758 | 0.093 |
| | **n** | 32 | 32 | 32 | 32 | 32 | 32 | 32 | 27 | 31 | 32 |

**Supplementary table 6. EPSPS sequence of *Vibrio cholerae* serotype O1 (vcEPSPS) and amino acids in the active site**

| | |
|---|---|
| >sp\|Q9KRB0\|AROA_VIBCH 3-phosphoshikimate 1-carboxyvinyltransferase OS=Vibrio cholerae serotype O1 (strain ATCC 39315 / El Tor Inaba N16961) GN=aroA PE=1 SV=1<br>MESLTLQPIELISGEVNLPGS**KS**VSN**R**ALLLAALASGTTRLTNLLDSDDIRHMLNALTKL<br>GVNYRLSADKTTCEVEGLGQAFHTTQPLELFLG**N**A**G**TAMRPLAAALCLGQGDYVLTGEPR<br>MKE**R**PIGHLVDALRQAGAQIEYLEQENFPPLRIQGTGLQAGTVTIDGSI**SSQ**FLTAFLMS<br>APLAQGKVTIKIVGELV**S**KPYIDITLHIMEQFGVQVINHDYQEFVIPAGQSYVSPGQFLV<br>EGDASSASYFLAAAAIKGGEVKVTGIGKNSIQGDIQFADALEKMGAQIEWGDDYVIARRG<br>ELNAVDLDFNHIP**D**AAMTIATTALFAKGTTAIRNVY**N**WRV**KE**TD**R**LAAMATELRKVGATV<br>EEGEDFIVITPPTKLIHAAIDTYDDH**R**MAMCFSLVALSDTPVTINDPKCTS**K**TFPDYFDK<br>FAQLSR | **Active site:** (Amino acid,Position)<br><br>**K,22** **S,169** **K,340**<br>**S,23** S,170 **E,341**<br>**R,27** **Q,171** **R,344**<br>N,94 **S,197** **R,386**<br>**G,96** **D,313** K,411<br>**R,124** N,336 |

**Supplementary table 7. Amino acids substitution by lifestyle and taxonomy.**

| Lifestyle | N | AA substitutions: Active Site (AS) | Active Site Area (ASA) | No Active Site (NO_AS) | AS / NO_AS | Permutation test (p-value): FL | FHA | IP |
|---|---|---|---|---|---|---|---|---|
| free-living (FL) | 17 | 1.596 | 1.901 | 2.121 | 0.753 | 1 | - | - |
| facultative host-associated (FHA) | 13 | 1.457 | 1.670 | 1.917 | 0.769 | 0.059 | 1 | - |
| intracellular parasite (IP) | 2 | 2.000 | 2.172 | 2.263 | 0.876 | - | - | 1 |
| **Taxonomy** | | | | | | F | P | A |
| Firmicutes (F) | 8 | 1.618 | 2.025 | 2.170 | 0.738 | 1 | - | - |
| Proteobacteria (P) | 15 | 1.489 | 1.754 | 2.029 | 0.744 | 0.049 | - | - |
| Actinobacteria (A) | 6 | 1.529 | 1.593 | 1.853 | 0.826 | 0.023 | 0.049 | 1 |

**Supplementary table 8. Amino acids substitution in the active site divided by substitutions in other amino acids.**

| | | All | Class I [S] | [S/R/U] | Class II [R] |
|---|---|---|---|---|---|
| **Max** | | 0.91 | 0.91 | 0.88 | 0.88 |
| **Min** | | 0.54 | 0.68 | 0.65 | 0.54 |
| **Median** | | 0.75 | 0.75 | 0.75 | 0.77 |
| **t-test (p-value)** | **Class I [S]** | | 1 | 0.449 (n.s.) | 0.378 (n.s.) |
| | **Class** | | - | 1 | 0.441 (n.s.) |
| | **Class II** | | - | - | 1 |

S: sensitive; R: resistant; U: unclassified; n.s.: not significant

### Supplementary table 9. Variability of amino acids in the EPSPS sequences

| P | AS | vcEPSPS | Alternative |
|---|---|---|---|
| 1 | | M | MVNPSTKLD |
| 2 | | E | EKTQLCADMS |
| 3 | | S | SVFLRMEKADTQYHIPN |
| 4 | | L | LTKMFNRYWVASH |
| 5 | | T | TFKYLIPSNWAVREDH |
| 6 | | L | LSIRVQAMEDPKTHN |
| 7 | | Q | QGRDVESPIATHKYWN |
| 8 | | P | PRNTESGKMDILWHAV |
| 9 | | I | ILAVRSTKYQE |
| 10 | | E | ATERGNKQMSPDHV |
| 11 | | L | RQLHPKNSATIC |
| 12 | | I | VNILFPAK |
| 13 | | S | DPSEHQKNYRMTGLA |
| 14 | | G | GTARVSQHPE |
| 15 | | E | TAVESNKRDYIQ |
| 16 | | V | IVMYAL |
| 17 | | N | NERQCTVHASDKY |
| 18 | | L | LVITA |
| 19 | | P | PEAID |
| 20 | | G | GPSAQ |
| 21 | | S | SD |
| 22 | * | K | K |
| 23 | * | S | SP |
| 24 | | V | VIHQLMES |
| 25 | | S | STGVA |
| 26 | | N | NHLQTI |
| 27 | * | R | R |
| 28 | | A | ASVTLYF |
| 29 | | L | LVMIFA |
| 30 | | L | LMIFV |
| 31 | | L | LFWIAVT |
| 32 | | A | AGVS |
| 33 | | A | ASLTGV |
| 34 | | L | LFVIM |
| 35 | | A | ASPNTG |
| 36 | | S | HSCKNRETAQDGPI |
| 37 | | G | GKSRAEN |
| 38 | | T | KTVEAPIDQ |
| 39 | | T | TSCAYVIN |
| 40 | | R | VATRKIFNHQYE |
| 41 | | L | LVFIA |
| 42 | | T | TRMYKHESNIVDQ |
| 43 | | N | NKDGHS |
| 44 | | L | LCIFAPDVY |
| 45 | | L | LPSAV |
| 46 | | D | DLRKPSFAMHWTEN |
| 47 | | S | SAGNC |
| 48 | | D | DAEPRKGLIFVQ |
| 49 | | D | DEN |
| 50 | | I | VICST |
| 51 | | R | RVLQEDMNSIGY |
| 52 | | H | HYSTALRVDGN |
| 53 | | M | MTSQAC |
| 54 | | L | LIMTVKFAGS |
| 55 | | N | NGQSADETRHK |
| 56 | | A | AMVCISNG |
| 57 | | L | LMFCAIV |
| 58 | | T | TSKRANEQGD |
| 59 | | K | AGKDNEQTLRSH |
| 60 | | L | LMF |
| 61 | | G | G |
| 62 | | V | VLIATC |
| 63 | | N | SNAHDQERTKG |
| 64 | | Y | YICFVL |
| 65 | | R | TESRMKVDQ |
| 66 | | L | LIQREFVSNKYHP |
| 67 | | S | SAENDILYTRG |
| 68 | | A | ANDTSELGRKHPQW |
| 69 | | D | DGKNSEPHTL |
| 70 | | K | RQKDGENSAFPV |
| 71 | | T | TVGDQLSHAIWRK |
| 72 | | T | RTCVQISHGYPELA |
| 73 | | C | CILVAPQR |
| 74 | | E | EDTVIHYGKRNA |
| 75 | | V | IGVSLPKM |
| 76 | | E | ITAEVQHKNDPRL |
| 77 | | G | GNFSEP |
| 78 | | L | NQEVLITYPASFC |
| 79 | | G | GSEDPATYRKN |
| 80 | | Q | GQMFASDHKVYTREC |
| 81 | | A | PVALSDEQTFGHRKI |
| 82 | | F | LTFGQCKSHIPNVMY |
| 83 | | H | HRQKELTNAFDPSVG |
| 84 | | T | APSTKNQVIDRLG |
| 85 | | T | EKPNQGSATVDCFL |
| 86 | | Q | GSAEQDTPHRKINC |
| 87 | | P | AETKDPQCHVILNRS |
| 88 | | L | LRNQPTIWKVAEF |
| 89 | | E | EDATYSPVNK |
| 90 | | L | LVFIPD |
| 91 | | F | FNDHGYQAKI |
| 92 | | L | LAMVPCTGIF |
| 93 | | G | GNKAV |
| 94 | * | N | NSLTGEM |
| 95 | | A | ASV |
| 96 | * | G | GA |
| 97 | | T | TILMV |
| 98 | | A | ASTVLG |
| 99 | | M | MILFTVA |
| 100 | | R | R |
| 101 | | P | PLFTM |
| 102 | | L | LMIFVYS |
| 103 | | A | ASLTPMICV |
| 104 | | A | AGP |
| 105 | | A | AIVLFM |
| 106 | | L | LSATV |
| 107 | | C | CATSGLV |
| 108 | | L | LGNIFATMVP |
| 109 | | G | GDQATRSFVIPNL |
| 110 | | Q | SAQPEHTRDFKLGN |
| 111 | | G | NFGTEVAIRS |
| 112 | | D | DENSLFHYPKTRQG |
| 113 | | Y | ISTYVAF |
| 114 | | V | VEKTCALFRINHDYQ |
| 115 | | L | LMVIFT |
| 116 | | T | TVSFIQADRHLMY |
| 117 | | G | GECYA |
| 118 | | E | EDNSRVK |
| 119 | | P | PLHDENTASQVYK |
| 120 | | R | RSPTQIFY |
| 121 | | M | MLIA |
| 122 | | K | KSAGQRNFHYEML |
| 123 | | E | EKRGASVTCDN |
| 124 | * | R | RL |
| 125 | | P | PVDS |
| 126 | | I | IMSQLV |
| 127 | | G | GDKQARNVL |
| 128 | | H | HRPQEFD |
| 129 | | L | LVIDMQ |
| 130 | | V | VATLIFDMS |
| 131 | | D | DTELIYNSRQGAK |
| 132 | | A | ASPFG |
| 133 | | L | LFM |
| 134 | | R | RGKSIENHTQA |
| 135 | | Q | LQKENAVTSGHRI |
| 136 | | A | GMALFCIY |
| 137 | | G | GSNED |
| 138 | | A | AVLTIM |
| 139 | | Q | KNQGSEDVAHTRL |
| 140 | | I | IVFCMA |
| 141 | | E | TDQESRLHVMNFY |
| 142 | | Y | YCGQAF |
| 143 | | L | LQERKITVHAF |
| 144 | | E | ESTGKAQLDN |
| 145 | | Q | QAEDSNGHLKIPR |
| 146 | | E | ESKRLGPNDAVI |
| 147 | | N | NGDQHTESYARW |
| 148 | | F | YTFLISGAKRHM |
| 149 | | P | PALTRM |
| 150 | | P | PRQF |
| 151 | | L | LMVFITAS |
| 152 | | R | RHQKTSVAICML |
| 153 | | I | LIVNTM |
| 154 | | Q | QRTKSNHLVEWIGA |
| 155 | | G | GPSAHN |
| 156 | | T | TNSGAPKRLEQD |
| 157 | | G | GTSKNEDQPVARL |
| 158 | | L | FPLNVIMTQEAGSK |
| 159 | | Q | TSIKNQLAPDHRGC |
| 160 | | A | GLAPKSHFDTQNC |
| 161 | | G | GRKIPMSEANYQDF |
| 162 | | T | NHDGESTQPIYVKRM |
| 163 | | V | VIYRDNETCFLAH |
| 164 | | T | DETSAHQYINGKRFP |
| 165 | | I | VYILQSMHAR |
| 166 | | D | DEPLSIATRMVK |
| 167 | | G | GLIVPSATC |
| 168 | | S | SPAVQDELIRKNM |
| 169 | * | I | VIAPSGKELT |
| 170 | * | S | SDA |
| 171 | * | S | SAT |
| 172 | | Q | Q |
| 173 | | F | FVLIY |
| 174 | | L | LKAVI |
| 175 | | T | TS |
| 176 | | A | APGTCF |
| 177 | | F | LVFIAM |
| 178 | | L | LMIAV |
| 179 | | M | MLFIY |
| 180 | | S | TASVLIGP |
| 181 | | A | AGCSTL |
| 182 | | P | PLASHRFM |
| 183 | | L | LYQRSFCMIKN |
| 184 | | A | AVGFSYLPTM |
| 185 | | Q | PDVQEAKNTRGH |
| 186 | | G | EQKGNYDTRSC |
| 187 | | K | DAKNEVITPGSRQL |
| 188 | | V | TVSLFRAIC |
| 189 | | T | VTIASDEHRLNQ |
| 190 | | I | IVTFYLE |
| 191 | | K | RASTKVQEDHN |
| 192 | | I | IVHESPLTM |
| 193 | | V | KVIELTGADMQN |
| 194 | | G | GEKNDRPVATS |
| 195 | | E | DEGKNPSHQTA |
| 196 | | L | LIEHVGKSA |
| 197 | * | V | VEYCITKPLSM |
| 198 | | S | SRTQEAKP |
| 199 | | K | KRNEAILQPMHGT |
| 200 | | P | PNHLSMGKD |
| 201 | | Y | YTHMW |
| 202 | | I | IRTEKQFLVS |
| 203 | | D | DNEAHRL |
| 204 | | I | IYMHLTAQN |
| 205 | | T | TIMS |
| 206 | | L | LEVAIMTC |
| 207 | | H | NTHDEQRWAK |
| 208 | | I | LMIFWSKTANVD |
| 209 | | M | MVLIGFRKT |
| 210 | | E | KAEQGRNSHD |
| 211 | | Q | TAQSFHLIRGDE |
| 212 | | F | FLAYSGCR |
| 213 | | G | GEKNASHP |
| 214 | | V | VGIFCAKSMY |
| 215 | | Q | ENDQTHRKSPAVI |
| 216 | | V | IVLFYKE |
| 217 | | I | EASTMIDLRVQHKG |
| 218 | | N | NVSRAQDKYTFMEHI |
| 219 | | H | QHNRVDEASKYPM |
| 220 | | D | HNGRASQDETVIP |
| 221 | | Y | YQGSKENPLFDAWRI |
| 222 | | Q | QEKRTDNVSYFGLAC |
| 223 | | E | QREKISDLTVGH |
| 224 | | F | FITWAVLMKNRHSE |
| 225 | | V | VAITLRSYQEFHK |
| 226 | | I | VITKRSFLAYND |
| 227 | | P | KQRPLTDAFHEVISC |
| 228 | | A | GVASQEPLDIHTN |
| 229 | | G | GEAKNRSPDT |
| 230 | | Q | QRPGNKIDEVCSA |
| 231 | | S | SQKLHEDNATPRYIGC |
| 232 | | Y | YSKQTFRLHMI |
| 233 | | V | QRHLVIPEDAFKMTG |
| 234 | | S | SKTIARQENFDLG |
| 235 | | P | PGAQSFDT |
| 236 | | G | GRQTAFDKISNCE |
| 237 | | Q | TQRDVNHSKEI |
| 238 | | F | YVFIWLAMR |
| 239 | | L | LVMTQEFSDKIHNYGR |
| 240 | | V | VI |
| 241 | | E | EPGA |
| 242 | | G | GPNAT |
| 243 | | D | D |
| 244 | | A | AIVYFLP |
| 245 | | S | STAG |
| 246 | | S | STNLG |
| 247 | | A | AVLTSG |
| 248 | | S | SAVFT |
| 249 | | Y | YFAPLV |
| 250 | | F | FWLIY |
| 251 | | L | LMAIVWC |
| 252 | | A | AVISMGLF |
| 253 | | A | ATVSCLF |
| 254 | | A | AGFVTCLS |
| 255 | | A | ALIGVSYM |
| 256 | | I | IVLTMSY |
| 257 | | K | KNRPLSTQFVGAIME |
| 258 | | G | GNSQKEDPA |
| 259 | | G | GAST |
| 260 | | E | TSERHAKPDNVGQ |
| 261 | | V | VILSTYF |
| 262 | | K | KTVLIYRCDE |
| 263 | | V | VILP |
| 264 | | T | TRQKEHPANGVIM |
| 265 | | G | GNDHA |
| 266 | | I | IVLWYF |
| 267 | | G | GPSNLQRFET |
| 268 | | K | RIKVMLSQEPAT |
| 269 | | N | NKSGQDVRAETHL |
| 270 | | S | SPEDT |
| 271 | | I | MTVINSLEYRK |
| 272 | | Q | QRELF |
| 273 | | G | GTPSAN |
| 274 | | D | DGARES |
| 275 | | I | IVMKDNRAGEHTL |
| 276 | | Q | RHQLEGSVAFKI |
| 277 | | F | FLIAVY |
| 278 | | A | AVILFRPYM |
| 279 | | D | DESFANRHKGCQ |
| 280 | | A | VALIYTE |
| 281 | | L | LIVYFA |
| 282 | | E | ERKTQSMHA |
| 283 | | K | KLAQTNSRDE |
| 284 | | M | MLTIF |
| 285 | | G | GDN |
| 286 | | A | AGTVCIFL |
| 287 | | Q | TRVQKLSHNDAEMI |
| 288 | | I | IVLMFSA |
| 289 | | E | CTESVKAHLIQNR |
| 290 | | W | WLIVMYFHCRD |
| 291 | | G | GSTEDAYKLFHIV |
| 292 | | D | DEANCKSLVG |
| 293 | | D | DRASIGHNEPVTK |
| 294 | | Y | YFRTDNHSPAGWV |
| 295 | | V | IVLEMTDHNFYC |
| 296 | | I | SATIMVLERD |
| 297 | | A | CVANIMSGPLKETD |
| 298 | | R | TMREKPHFIADGS |
| 299 | | R | RGHQSKTPYENDAI |
| 300 | | G | GASETFIDQPN |
| 301 | | E | EKDQSATGPLYRIV |
| 302 | | L | LVSFYRIT |
| 303 | | N | NHSKRTADQEG |
| 304 | | A | AGSNPD |
| 305 | | V | IVTLAGFRQY |
| 306 | | D | DERGNSTAVKCQ |
| 307 | | L | MLIVFC |
| 308 | | D | DCGVAENSP |
| 309 | | F | MAFGYLQKRVIEP |
| 310 | | N | NDASETRH |
| 311 | | H | HILNRQPASDEWM |
| 312 | | I | IALMFVTG |
| 313 | * | P | PKIGTS |
| 314 | | D | DGREKNS |
| 315 | | A | AETLIP |
| 316 | | A | AFLITVY |
| 317 | | M | MPIDTL |
| 318 | | T | TVISEANF |
| 319 | | I | IFLV |
| 320 | | A | AFTPSGVI |
| 321 | | T | TILVAG |
| 322 | | T | ATLISRV |
| 323 | | A | ACMSGLF |
| 324 | | L | LATNCSVM |
| 325 | | F | FQLNYIAC |
| 326 | | A | ASVLT |
| 327 | | K | KNEQHTSVAGDR |
| 328 | | G | GNSTDKAQ |
| 329 | | T | TNVIPSKQDER |
| 330 | | T | TSLIFC |
| 331 | | A | TVACHIRYKMEDNS |
| 332 | | I | LVIMF |
| 333 | | R | RKCYSVGITDH |
| 334 | | N | NGDS |
| 335 | | V | IAVGL |
| 336 | * | Y | YEQHALRKSDG |
| 337 | | N | NEGSHDVIR |
| 338 | | W | WLATI |
| 339 | | R | RKN |
| 340 | * | V | VDNGSAFYLI |
| 341 | * | K | KTHQ |
| 342 | | E | EK |
| 343 | | T | TSC |
| 344 | * | D | DNSQPG |
| 345 | | R | RC |
| 346 | | L | LIVA |
| 347 | | A | FELAYQDIKNSTG |
| 348 | | A | ATVSD |
| 349 | | M | MVILTF |
| 350 | | A | ALTVSIHRNE |
| 351 | | T | TVDEASHQGRKN |
| 352 | | E | EGATINMV |
| 353 | | L | LIFV |
| 354 | | R | RNKTGQDESA |
| 355 | | K | KQSRATLGN |
| 356 | | V | VLMIAYNC |
| 357 | | G | GKED |
| 358 | | A | AVGILFS |
| 359 | | T | EVTQDNRKSCHILGAY |
| 360 | | V | VICLASF |
| 361 | | E | ETKDHQRGIAV |
| 362 | | E | ESPAYDVTQG |
| 363 | | G | GKTCFLDVE |
| 364 | | E | HPEDARNGVYSQ |
| 365 | | D | DGHRKYNEM |
| 366 | | F | YFGRLDNTPSA |
| 367 | | I | IAMLSVEF |
| 368 | | V | RVIKYLQHTMSED |
| 369 | | I | IVKRSH |
| 370 | | T | TSKEHYNGDFARV |
| 371 | | P | PGRNT |
| 372 | | P | PVLKASRNDGIT |
| 373 | | T | EAGTSCQKLIRNVD |
| 374 | | K | KQSRTAVMPFGE |
| 375 | | L | LRKYTIPF |
| 376 | | I | NQKMHIYARTLGPS |
| 377 | | H | FHLVTYGAPNDS |
| 378 | | A | AVNSTGKPE |
| 379 | | A | EDSARNQTGVIPKL |
| 380 | | I | IVTMWSLF |
| 381 | | D | AGEDNSHRQKPT |
| 382 | | T | TSAGC |
| 383 | | Y | YAFHLNWR |
| 384 | | D | NYDGKATRL |
| 385 | | D | D |
| 386 | * | H | H |
| 387 | | R | RG |
| 388 | | M | MINLV |
| 389 | | A | AGFCI |
| 390 | | M | MTLAH |
| 391 | | C | CAMSF |
| 392 | | F | FTLGNSA |
| 393 | | S | SAVGLFITQ |
| 394 | | L | LIVTAM |
| 395 | | V | VAITLM |
| 396 | | A | ASCGT |
| 397 | | L | LSIVMF |
| 398 | | S | SCAKITHRVMLGE |
| 399 | | D | DEANQKGSVRITMY |
| 400 | | T | TDGESALPKRHVQC |
| 401 | | P | PAGEQKSFDT |
| 402 | | V | VITEDLMHSN |
| 403 | | T | TAEDHVIKLSRQN |
| 404 | | I | IVLA |
| 405 | | N | LVNDEHRCYTKI |
| 406 | | D | DRKNHQGSE |
| 407 | | P | PAESTILFYVC |
| 408 | | K | KEQDGARHST |
| 409 | | C | CATGKHDNM |
| 410 | | T | TIVAYL |
| 411 | * | S | AGSNMRHEKTD |
| 412 | | K | KTVIDQC |
| 413 | | T | TSG |
| 414 | | F | FYLNICM |
| 415 | | P | P |
| 416 | | D | DGSTAEHYNQKR |
| 417 | | Y | YFS |
| 418 | | F | FGIPKHQLEAWVM |
| 419 | | D | EDKPQRTHLNSVAG |
| 420 | | K | QLKDHEIMWTRSV |
| 421 | | F | LCFRWIMA |
| 422 | | A | AENQDSKHRGLP |
| 423 | | Q | RGHQSNKTAELMI |
| 424 | | L | IMCLFDEPG |
| 425 | | S | SGITCMAVKEYNLQHR |
| 426 | | R | QTLARHDKNSGEVF |

P: Amino acid position in the reference sequence vcEPSPS; AS: binding amino acids in the EPSPS active site (*); vEPSPS: amino acids in the reference sequence; Alternative: other amino acids found in the ATGC

**SUPPLEMENTARY TABLE 10. Classification of the EPSPS in Lactobacillus sp.**

| | | | | | | | | | | | |
|---|---|---|---|---|---|---|---|---|---|---|---|
| Lactobacillus uli | Classlb | Lactobacillus fabifermentans T30PCM01 | Unclassified | Lactobacillus nodensis DSM 19682 = JCM 14932 = NBRC 1( | Unclassified | Lactobacillus ingluviei | Unclassified | Lactobacillus panis DSM 6035 | Unclassified | Lactobacillus farraginis DSM 18382 = JCM 141( | Unclassified |
| Lactobacillus fabifermentans T30PCM01 | ClassII | Lactobacillus fabifermentans T30PCM01 | Unclassified | Lactobacillus equigenerosi DSM 18793 = JCM 14505 | Unclassified | Lactobacillus secaliphilus | Unclassified | Lactobacillus paracasei (strain ATCC 334 / BCRC 17002 / | Unclassified | Lactobacillus suebicus DSM 5007 = KCTC 3549 | Unclassified |
| Lactobacillus mucosae LM1 | ClassII | Lactobacillus reuteri (strain DSM 20016) | Unclassified | Lactobacillus coryniformis subsp coryniformis CECT 5711 | Unclassified | Lactobacillus ozensis DSM 23829 = JCM 17196 | Unclassified | Lactobacillus pasteurii DSM 23907 = CRBIP 2476 | Unclassified | Lactobacillus tucceti DSM 20183 | Unclassified |
| Lactobacillus rennini DSM 20253 | ClassII | Lactobacillus mindensis DSM 14500 | Unclassified | Lactobacillus oris PB013-T2-3 | Unclassified | Lactobacillus shenzhenensis LY-73 | Unclassified | Lactobacillus ghanensis DSM 18630 | Unclassified | Lactobacillus coryniformis subsp coryniformis C | Unclassified |
| Lactobacillus gastricus PS3 | ClassII | Lactobacillus oligofermentans DSM 15707 = LMG 22743 | Unclassified | Lactobacillus harbinensis DSM 16991 | Unclassified | Lactobacillus parafarraginis F0439 | Unclassified | Lactobacillus selangorensis | Unclassified | Lactobacillus algidus DSM 15638 | Unclassified |
| Lactobacillus collinoides DSM 20515 = JCM 1123 | ClassII | Lactobacillus plantarum (strain ATCC BAA-793 / NCIMB 8826 / | Unclassified | Lactobacillus acetotolerans | Unclassified | Lactobacillus brevis subsp gravesensis ATCC 27305 | Unclassified | Lactobacillus odoratitofui DSM 19909 = JCM 15043 | Unclassified | Lactobacillus spicheri | Unclassified |
| Lactobacillus kunkeei | ClassII | Lactobacillus malefermentans DSM 5705 = KCTC 3548 | Unclassified | Lactobacillus hayakitensis DSM 18933 = JCM 14209 | Unclassified | Lactobacillus algidus DSM 15638 | Unclassified | Lactobacillus fuchuensis DSM 14340 = JCM 11249 | Unclassified | Lactobacillus mali KCTC 3596 = DSM 20444 | Unclassified |
| Lactobacillus odoratitofui DSM 19909 = JCM 15043 | ClassII | Lactobacillus oeni DSM 19972 | Unclassified | Lactobacillus ozensis DSM 23829 = JCM 17196 | Unclassified | Lactobacillus tucceti DSM 20183 | Unclassified | Lactobacillus pontis DSM 8475 | Unclassified | Lactobacillus cacaonum DSM 21116 | Unclassified |
| Lactobacillus ginsenosidimutans | ClassII | Lactobacillus camelliae DSM 22697 = JCM 13995 | Unclassified | Lactobacillus agilis DSM 20509 | Unclassified | Lactobacillus kalixensis DSM 16043 | Unclassified | Lactobacillus vaginalis DSM 5837 = ATCC 49540 | Unclassified | Lactobacillus pobuzihii | Unclassified |
| Lactobacillus coryniformis subsp coryniformis CECT 5711 | ClassII | Lactobacillus farciminis | Unclassified | Lactobacillus oryzae JCM 18671 | Unclassified | Lactobacillus capillatus DSM 19910 | Unclassified | Lactobacillus hamsteri DSM 5661 = JCM 6256 | Unclassified | Lactobacillus dextrinicus DSM 20335 | Unclassified |
| Lactobacillus lindneri DSM 20690 = JCM 11027 | ClassII | Lactobacillus farraginis DSM 18382 = JCM 14108 | Unclassified | Lactobacillus diolivorans DSM 14421 | Unclassified | Lactobacillus ghanensis DSM 18630 | Unclassified | Lactobacillus parabrevis ATCC 53295 | Unclassified | Lactobacillus oris PB013-T2-3 | Unclassified |
| Lactobacillus suebicus DSM 5007 = KCTC 3549 | ClassII | Lactobacillus nodensis DSM 19682 = JCM 14932 = NBRC 1071( | Unclassified | Lactobacillus coleohominis 101-4-CHN | Unclassified | Lactobacillus florum 8D | Unclassified | Lactobacillus vaccinostercus DSM 20634 | Unclassified | Lactobacillus ruminis (strain ATCC 27782 / RF3) | Unclassified |
| Lactobacillus vaccinostercus DSM 20634 | ClassII | Lactobacillus satsumensis DSM 16230 = JCM 12392 | Unclassified | Lactobacillus tucceti DSM 20183 | Unclassified | Lactobacillus nodensis DSM 19682 = JCM 14932 = NBI | Unclassified | Lactobacillus senioris DSM 24302 = JCM 17472 | Unclassified | Lactobacillus xiangfangensis | Unclassified |
| Lactobacillus ozensis DSM 23829 = JCM 17196 | ClassII | Lactobacillus oligofermentans DSM 15707 = LMG 22743 | Unclassified | Lactobacillus frumenti DSM 13145 | Unclassified | Lactobacillus lindneri DSM 20690 = JCM 11027 | Unclassified | Lactobacillus sakei subsp sakei (strain 23K) | Unclassified | Weissella viridescens (Lactobacillus viridescens | Unclassified |
| Lactobacillus bifermentans DSM 20003 | ClassII | Lactobacillus sp ASF360 | Unclassified | Lactobacillus nodensis DSM 19682 = JCM 14932 = NBRC 1( | Unclassified | Lactobacillus senioris DSM 24302 = JCM 17472 | Unclassified | Lactobacillus odoratitofui DSM 19909 = JCM 15043 | Unclassified | Lactobacillus vaginalis DSM 5837 = ATCC 4954 | Unclassified |
| Lactobacillus manihotivorans DSM 13343 = JCM 12514 | ClassII | Lactobacillus siliginis | Unclassified | Lactobacillus johnsonii (strain CNCM I-12250 / La1 / NCC 5: | Unclassified | Lactobacillus iners DSM 13335 | Unclassified | Lactobacillus animalis | Unclassified | Lactobacillus equi DPC 6820 | Unclassified |
| Lactobacillus mellis | ClassII | Lactobacillus lindneri DSM 20690 = JCM 11027 | Unclassified | Lactobacillus satsumensis DSM 16230 = JCM 12392 | Unclassified | Lactobacillus rennini DSM 20253 | Unclassified | Lactobacillus helveticus (strain DPC 4571) | Unclassified | Lactobacillus ingluviei | Unclassified |
| Lactobacillus satsumensis DSM 16230 = JCM 12392 | ClassII | Lactobacillus vini DSM 20605 | Unclassified | Lactobacillus buchneri CD034 | Unclassified | Lactobacillus paucivorans | Unclassified | Lactobacillus cacaonum DSM 21116 | Unclassified | Lactobacillus saerimneri 30a | Unclassified |
| Lactobacillus composti DSM 18527 = JCM 14202 | ClassII | Lactobacillus sanfranciscensis (strain TMW 11304) | Unclassified | Lactobacillus xiangfangensis | Unclassified | Lactobacillus capillatus DSM 19910 | Unclassified | Lactobacillus similis DSM 23365 = JCM 2765 | Unclassified | Lactobacillus fermentum (strain CECT 5716) | Unclassified |
| Lactobacillus capillatus DSM 19910 | ClassII | Lactobacillus fermentum (strain CECT 5716) | Unclassified | Lactobacillus bifermentans DSM 20003 | Unclassified | Lactobacillus sakei subsp sakei (strain 23K) | Unclassified | Lactobacillus crispatus (strain ST1) | Unclassified | Lactobacillus reuteri (strain DSM 20016) | Unclassified |
| Lactobacillus ghanensis DSM 18630 | ClassII | Lactobacillus mellifer | Unclassified | Lactobacillus salivarius (strain UCC118) | Unclassified | Lactobacillus buchneri CD034 | Unclassified | Lactobacillus dextrinicus DSM 20335 | Unclassified | Lactobacillus panis DSM 6035 | Unclassified |
| Lactobacillus kimchicus JCM 15530 | ClassII | Lactobacillus jensenii JV-V16 | Unclassified | Lactobacillus ceti DSM 22408 | Unclassified | Lactobacillus rossiae DSM 15814 | Unclassified | Lactobacillus farciminis | Unclassified | Lactobacillus farraginis DSM 18382 = JCM 141( | Unclassified |
| Lactobacillus hokkaidonensis JCM 18461 | ClassII | Lactobacillus ceti DSM 22408 | Unclassified | Olsenella uli (strain ATCC 49627 / DSM 7084 / CIP 109912 | Unclassified | Lactobacillus diolivorans DSM 14421 | Unclassified | Lactobacillus farciminis | Unclassified | Lactobacillus shenzhenensis LY-73 | Unclassified |
| Lactobacillus plantarum | ClassII | Lactobacillus thailandensis DSM 22698 = JCM 13996 | Unclassified | Lactobacillus mellis | Unclassified | Lactobacillus coleohominis DSM 14060 | Unclassified | Lactobacillus malefermentans DSM 5705 = KCTC 3548 | Unclassified | Lactobacillus saniviri JCM 17471 = DSM 24301 | Unclassified |
| Lactobacillus concavus DSM 17758 | ClassII | Lactobacillus amylophilus DSM 20533 = JCM 1125 | Unclassified | Lactobacillus bifermentans DSM 20003 | Unclassified | Lactobacillus collinoides DSM 20515 = JCM 1123 | Unclassified | Lactobacillus kisonensis F0435 | Unclassified | Lactobacillus farraginis DSM 18382 = JCM 141( | Unclassified |
| Lactobacillus ruminis (strain ATCC 27782 / RF3) | ClassII | Lactobacillus kimchicus JCM 15530 | Unclassified | Lactobacillus siliginis | Unclassified | Lactobacillus coleohominis 101-4-CHN | Unclassified | Lactobacillus farraginis DSM 18382 = JCM 14108 | Unclassified | Lactobacillus hokkaidonensis JCM 18461 | Unclassified |
| Lactobacillus wasatchensis | ClassII | Lactobacillus mindensis DSM 14500 | Unclassified | Lactobacillus brevis (strain ATCC 367 / JCM 1170) | Unclassified | Lactobacillus suebicus DSM 5007 = KCTC 3549 | Unclassified | Lactobacillus parafarraginis F0439 | Unclassified | Lactobacillus kunkeei | Unclassified |
| Lactobacillus parafarraginis F0439 | ClassII | Lactobacillus plantarum (strain ATCC BAA-793 / NCIMB 8826 / | Unclassified | Lactobacillus composti DSM 18527 = JCM 14202 | Unclassified | Lactobacillus saerimneri 30a | Unclassified | Lactobacillus gastricus PS3 | Unclassified | Lactobacillus agilis DSM 20509 | Unclassified |
| Lactobacillus similis DSM 23365 = JCM 2765 | ClassII | Lactobacillus hokkaidonensis JCM 18461 | Unclassified | Lactobacillus brantae DSM 23927 | Unclassified | Lactobacillus mindensis DSM 14500 | Unclassified | Lactobacillus nasuensis JCM 17158 | Unclassified | Lactobacillus concavus DSM 17758 | Unclassified |
| Lactobacillus kisonensis F0435 | ClassII | Lactobacillus fructivorans | Unclassified | Lactobacillus floricola DSM 23037 = JCM 16512 | Unclassified | Lactobacillus kisonensis F0435 | Unclassified | Lactobacillus manihotivorans DSM 13343 = JCM 12514 | Unclassified | Lactobacillus animalis | Unclassified |
| Lactobacillus buchneri CD034 | ClassII | Lactobacillus oeni DSM 19972 | Unclassified | Lactobacillus sp wkB8 | Unclassified | Lactobacillus fructivorans | Unclassified | Lactobacillus kunkeei | Unclassified | Lactobacillus spicheri | Unclassified |
| Lactobacillus fermentum (strain CECT 5716) | ClassII | Lactobacillus delbrueckii subsp bulgaricus (strain ATCC 11842 | Unclassified | Lactobacillus kimchicus JCM 15530 | Unclassified | Lactobacillus oryzae JCM 18671 | Unclassified | Lactobacillus wasatchensis | Unclassified | | |
| Lactobacillus dextrinicus DSM 20335 | ClassII | Lactobacillus rossiae DSM 15814 | Unclassified | Lactobacillus parabrevis ATCC 53295 | Unclassified | Lactobacillus paucivorans | Unclassified | Lactobacillus salivarius (strain UCC118) | Unclassified | | |
| Lactobacillus plantarum (strain ATCC BAA-793 / NCIMB 8826 / WCFS1) | ClassII | Lactobacillus sharpeae JCM 1186 = DSM 20505 | Unclassified | Lactobacillus ruminis (strain ATCC 27782 / RF3) | Unclassified | Lactobacillus vini DSM 20605 | Unclassified | Lactobacillus collinoides DSM 20515 = JCM 1123 | Unclassified | | |
| Lactobacillus fuchuensis DSM 14340 = JCM 11249 | ClassII | Lactobacillus aviarius subsp araffinosus DSM 20653 | Unclassified | Lactobacillus fuchuensis DSM 14340 = JCM 11249 | Unclassified | Lactobacillus wasatchensis | Unclassified | Lactobacillus acidophilus (strain ATCC 700396 / NCK56 / | Unclassified | | |
| Lactobacillus frumenti DSM 13145 | ClassII | Lactobacillus aviarius subsp araffinosus DSM 20653 | Unclassified | Lactobacillus ginsenosidimutans | Unclassified | Lactobacillus harbinensis DSM 16991 | Unclassified | Lactobacillus coleohominis DSM 14060 | Unclassified | | |
| Weissella viridescens (Lactobacillus viridescens) | ClassII | Lactobacillus floricola DSM 23037 = JCM 16512 | Unclassified | Lactobacillus perolens DSM 12744 | Unclassified | Lactobacillus ingluviei | Unclassified | Lactobacillus plantarum | Unclassified | | |
| Lactobacillus diolivorans DSM 14421 | ClassII | Lactobacillus brevis (strain ATCC 367 / JCM 1170) | Unclassified | Lactobacillus perolens DSM 12744 | Unclassified | Lactobacillus pobuzihii | Unclassified | Lactobacillus cacaonum DSM 21116 | Unclassified | | |
| Lactobacillus brevis subsp gravesensis ATCC 27305 | ClassII | Lactobacillus amylolyticus DSM 11664 | Unclassified | Lactobacillus frumenti DSM 13145 | Unclassified | Lactobacillus similis DSM 23365 = JCM 2765 | Unclassified | Lactobacillus vaccinostercus DSM 20634 | Unclassified | | |
| Lactobacillus malefermentans DSM 5705 = KCTC 3548 | ClassII | Lactobacillus concavus DSM 17758 | Unclassified | Lactobacillus mucosae LM1 | Unclassified | Lactobacillus hayakitensis DSM 18933 = JCM 14209 | Unclassified | Lactobacillus plantarum | Unclassified | | |
| Lactobacillus siliginis | ClassII | Lactobacillus ginsenosidimutans | Unclassified | Lactobacillus hominis DSM 23910 = CRBIP 24179 | Unclassified | Lactobacillus florum 8D | Unclassified | Lactobacillus equi DPC 6820 | Unclassified | | |
| Lactobacillus brevis subsp gravesensis ATCC 27305 | ClassIII | Lactobacillus secaliphilus | Unclassified | Lactobacillus sanfranciscensis (strain TMW 11304) | Unclassified | Lactobacillus pontis DSM 8475 | Unclassified | Lactobacillus rennini DSM 20253 | Unclassified | | |

SUPPLEMENTARY FIGURES

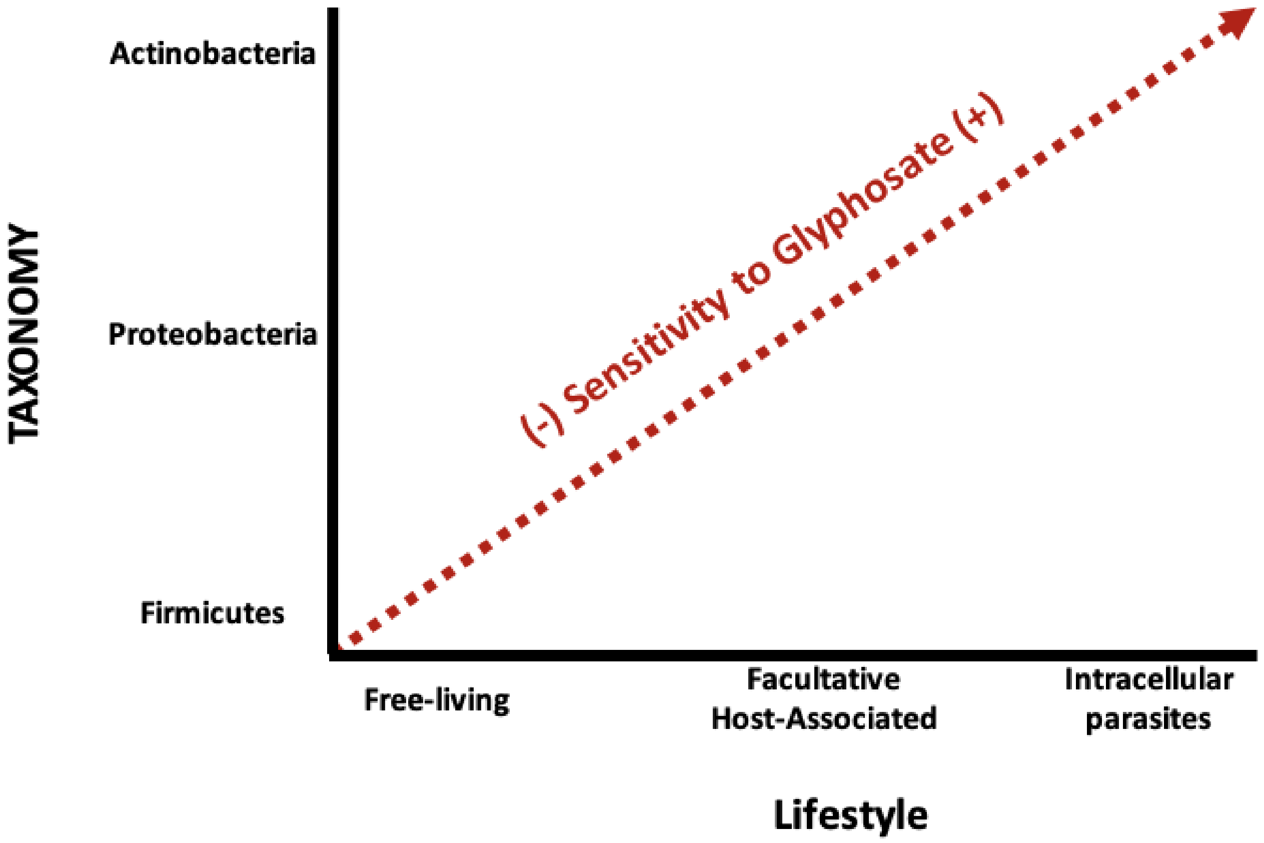

**Supplementary figure 1. Scheme of the differential sensitivity to glyphosate based on the three major taxonomic groups and lifestyle.**

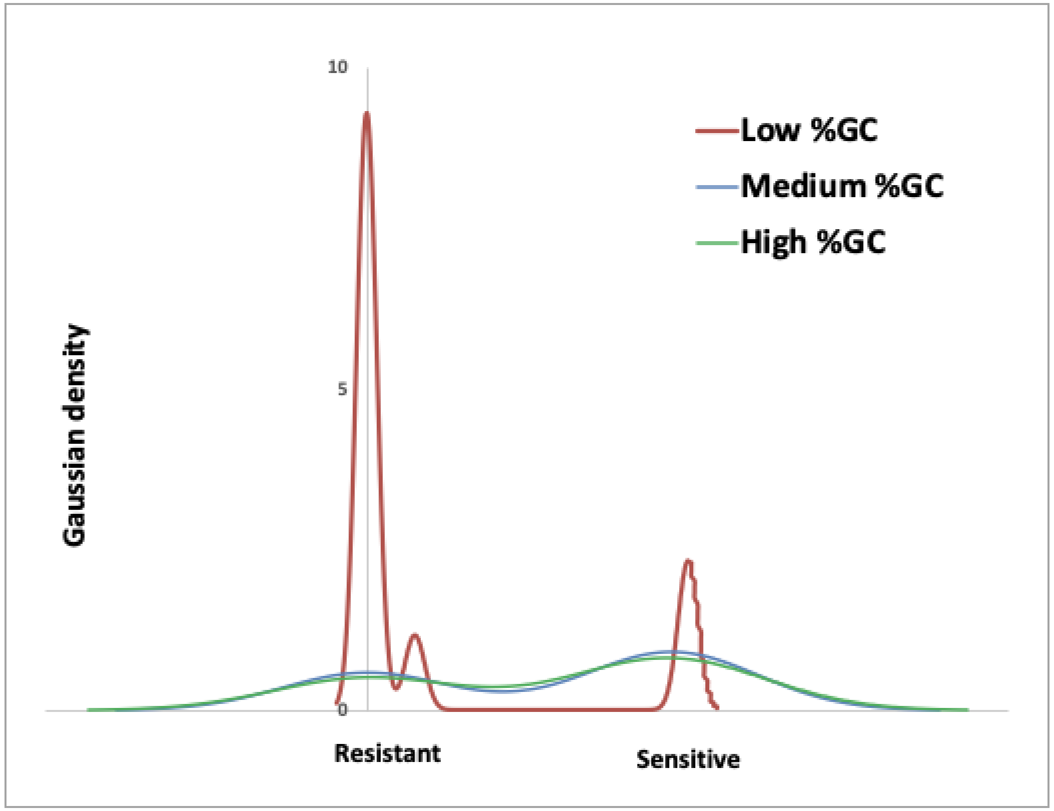

**Supplementary figure 2. Kernel density distribution of sensitivity in the ATGCs by %GC.** Genomes with low %GC (<40%) are more resistant to glyphosate (chi-test p-value < 0.001).

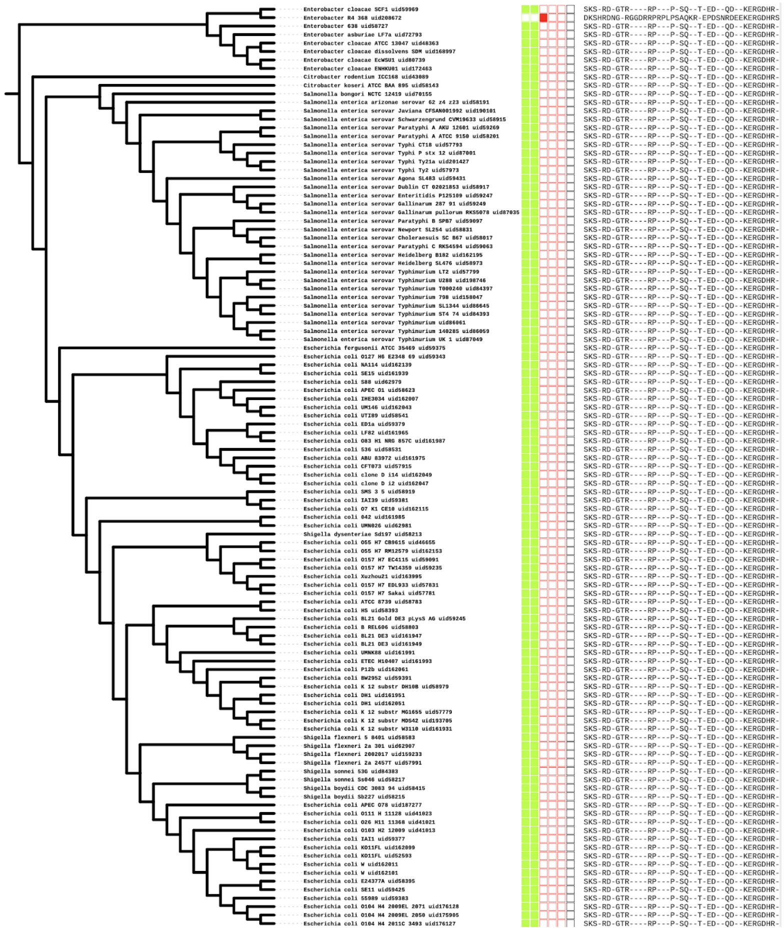

| | 21 | 22 | 23 | 26 | 27 | 49 | 94 | 96 | 97 | 100 | 104 | 117 | 118 | 123 | 124 | 125 | 128 | 132 | 133 | 150 | 170 |
|---|---|---|---|---|---|---|---|---|---|---|---|---|---|---|---|---|---|---|---|---|---|
| POS(vcEPSPS\|Class Iα) | S | K | S | X | R | D | X | G | T | R | X | X | X | X | R | P | X | X | X | P | X |
| POS(vcEPSPS\|Class Iβ) | S | K | S | X | R | D | X | G | T | R | X | X | X | X | R | P | X | X | X | P | X |
| POS(cbEPSPS\|Class II) | D | K | S | H | R | D | N | G | X | R | G | G | D | R | R | P | R | P | L | P | S |

| | 171 | 172 | 174 | 202 | 205 | 206 | 241 | 243 | 245 | 269 | 272 | 314 | 315 | 337 | 341 | 342 | 345 | 357 | 385 | 386 | 387 | 412 | 413 | 415 |
|---|---|---|---|---|---|---|---|---|---|---|---|---|---|---|---|---|---|---|---|---|---|---|---|---|
| POS(vcEPSPS\|Class Iα) | S | Q | X | X | T | X | E | D | X | X | Q | D | X | X | K | E | R | G | D | H | R | K | X | P |
| POS(vcEPSPS\|Class Iβ) | S | Q | X | X | T | X | E | D | X | X | Q | D | X | X | K | E | R | G | D | H | R | K | X | P |
| POS(cbEPSPS\|Class II) | A | Q | K | R | X | E | P | D | S | N | R | D | E | E | K | E | R | G | D | H | R | T | S | P |

**Supplementary figure 3. Example of change from sensitive to resistance in Enterobacter (ATGC001) with few amino acid mutations in the active site of the EPSPS.** Phylogenetic trees of the ATGC001. Filled boxes show bacteria sensitive (green) and resistant (red) to glyphosate. Boxed in order from left to right correspond to Class I alpha, Class I beta, Class II, Class III, Class IV and Unclassified. The amino acids in the active site are shown at right side of the figure.

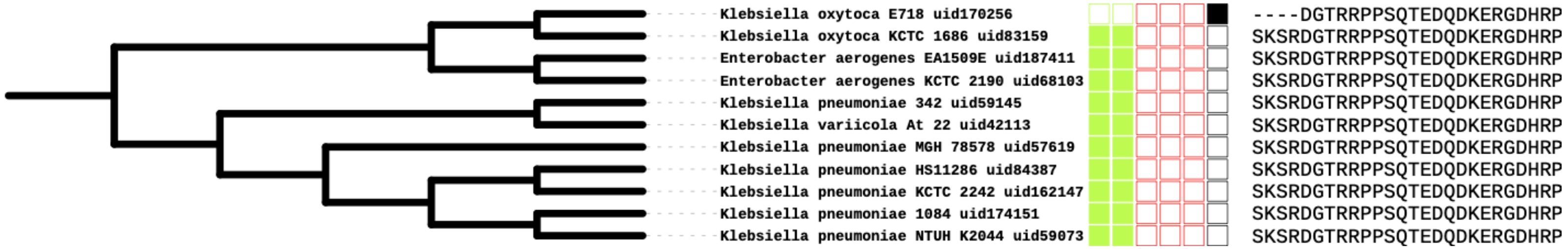

**Supplementary figure 4. Example of change from sensitive to unclassified in Enterobacter–Klebsiella (ATGC002) with four amino acid mutations deletions in the active site of the EPSPS.** Phylogenetic trees of the ATGC002. Filled boxes show bacteria sensitive (green) and resistant (red) to glyphosate. Boxed in order from left to right correspond to Class I alpha, Class I beta, Class II, Class III, Class IV and Unclassified. The amino acids in the active site are shown at right side of the figure.

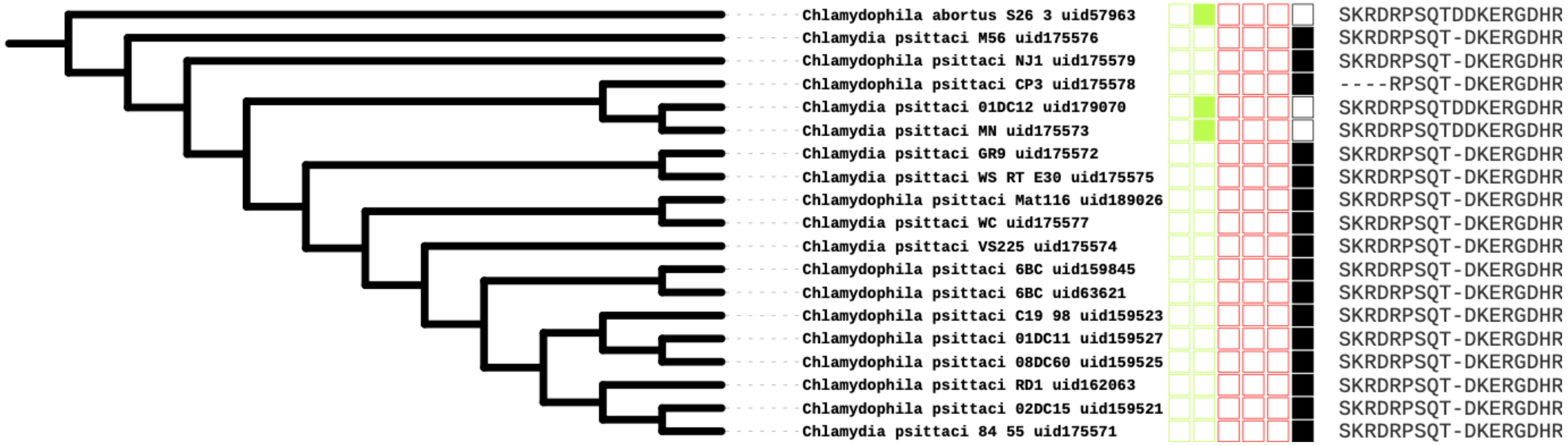

**Supplementary figure 5. Example of multiple changes from sensitive to unclassified in Chlamydia–Chlamydophila (ATGC022) with few amino acid deletions in the active site of the EPSPS.** Phylogenetic trees of the ATGC022. Filled boxes show bacteria sensitive (green) and resistant (red) to glyphosate. Boxed in order from left to right correspond to Class I alpha, Class I beta, Class II, Class III, Class IV and Unclassified. The amino acids in the active site are shown at right side of the figure.

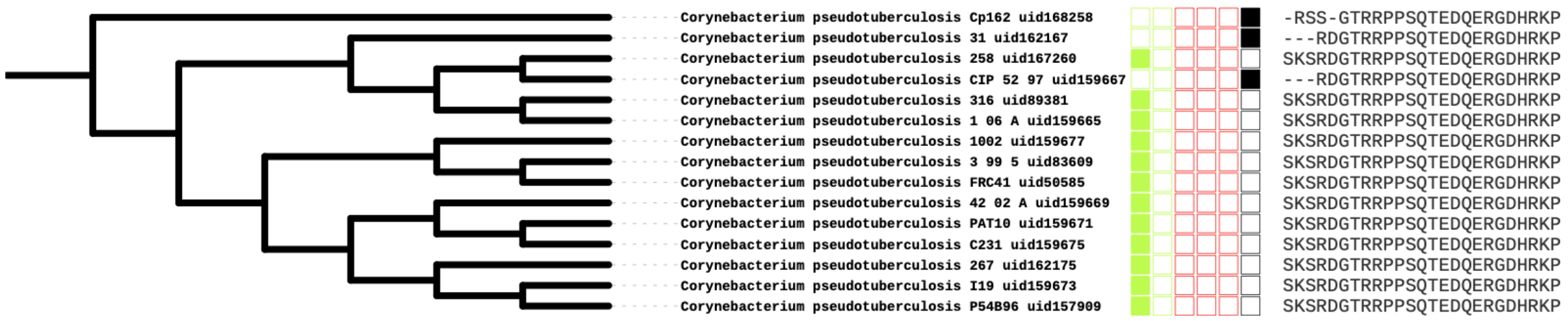

**Supplementary figure 6. Example of multiple changes from sensitive to unclassified in Corynebacterium (ATGC067) with few amino acid deletions in the active site of the EPSPS.** Phylogenetic trees of the ATGC067. Filled boxes show bacteria sensitive (green) and resistant (red) to glyphosate. Boxed in order from left to right correspond to Class I alpha, Class I beta, Class II, Class III, Class IV and Unclassified. The amino acids in the active site are shown at right side of the figure.

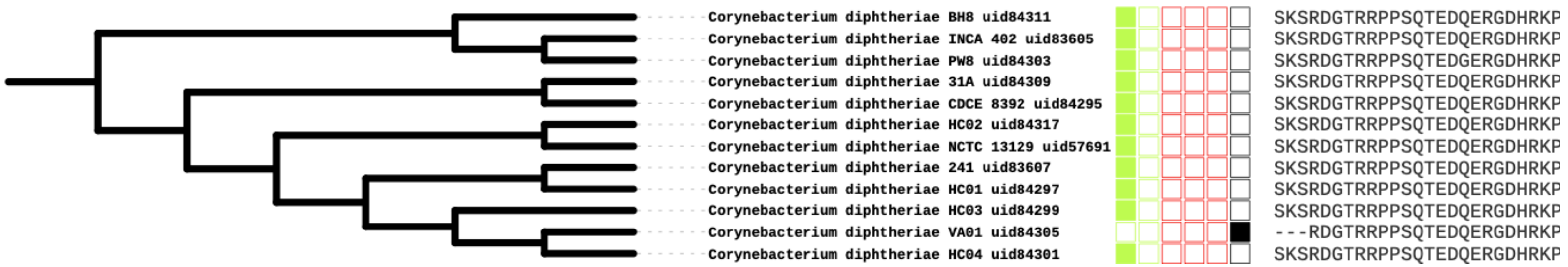

**Supplementary figure 7. Example of change from sensitive to unclassified in Corynebacterium (ATGC068) with few amino acid deletions in the active site of the EPSPS.** Phylogenetic trees of the ATGC068. Filled boxes show bacteria sensitive (green) and resistant (red) to glyphosate. Boxed in order from left to right correspond to Class I alpha, Class I beta, Class II, Class III, Class IV and Unclassified. The amino acids in the active site are shown at right side of the figure.

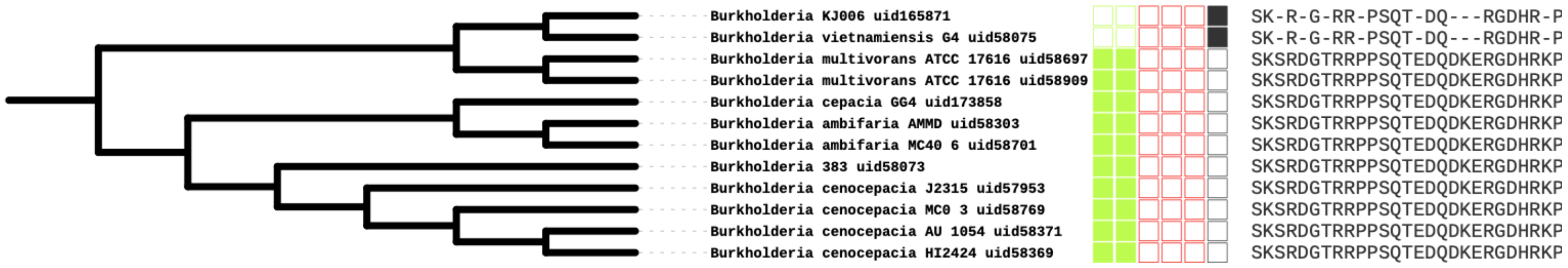

**Supplementary figure 8. Example of change from sensitive to unclassified in Burkholderia (ATGC090) with few amino acid deletions in the active site of the EPSPS.** Phylogenetic trees of the ATGC090. Filled boxes show bacteria sensitive (green) and resistant (red) to glyphosate. Boxed in order from left to right correspond to Class I alpha, Class I beta, Class II, Class III, Class IV and Unclassified. The amino acids in the active site are shown at right side of the figure.

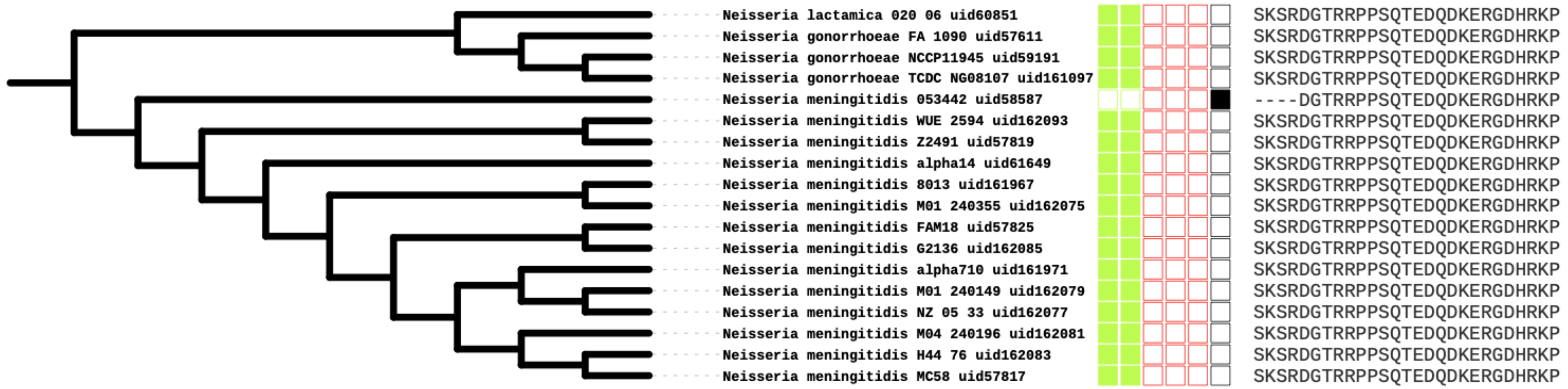

**Supplementary figure 9. Example of change from sensitive to unclassified in Neisseria (ATGC138) with few amino acid deletions in the active site of the EPSPS.** Phylogenetic trees of the ATGC138. Filled boxes show bacteria sensitive (green) and resistant (red) to glyphosate. Boxed in order from left to right correspond to Class I alpha, Class I beta, Class II, Class III, Class IV and Unclassified. The amino acids in the active site are shown at right side of the figure.

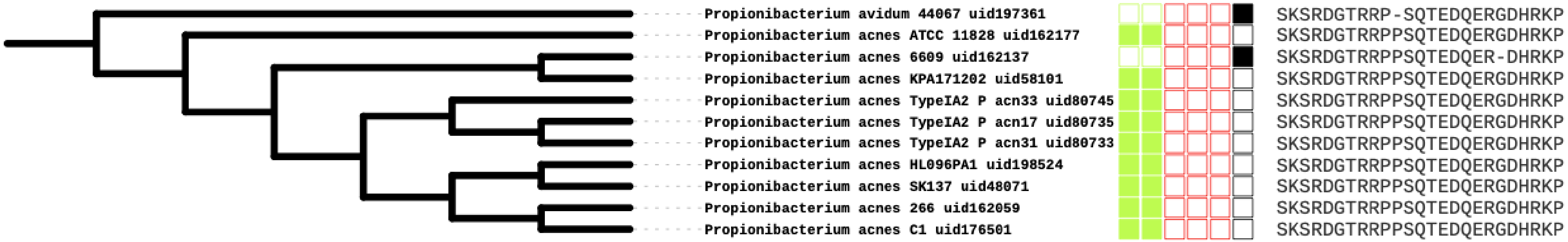

**Supplementary figure 10. Example of two independent changes from sensitive to unclassified in Propionibacterium (ATGC163) with a single amino acid deletion in the active site of the EPSPS.** Phylogenetic trees of the ATGC163. Filled boxes show bacteria sensitive (green) and resistant (red) to glyphosate. Boxed in order from left to right correspond to Class I alpha, Class I beta, Class II, Class III, Class IV and Unclassified. The amino acids in the active site are shown at right side of the figure.

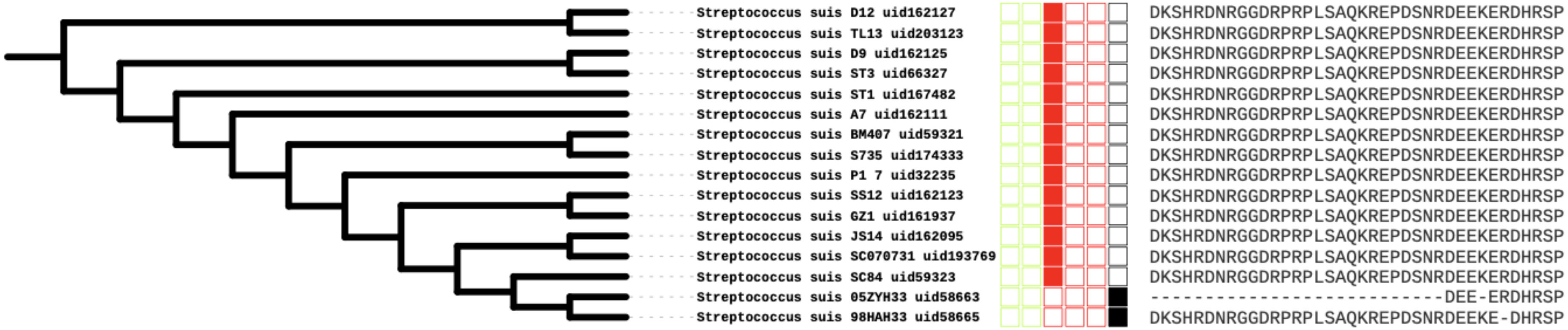

**Supplementary figure 11. Example of change from resistant (Class II) to unclassified in Streptococcus (ATGC005) with several amino acid deletions in the active site of the EPSPS.** Phylogenetic trees of the ATGC005. Filled boxes show bacteria sensitive (green) and resistant (red) to glyphosate. Boxed in order from left to right correspond to Class I alpha, Class I beta, Class II, Class III, Class IV and Unclassified. The amino acids in the active site are shown at right side of the figure.

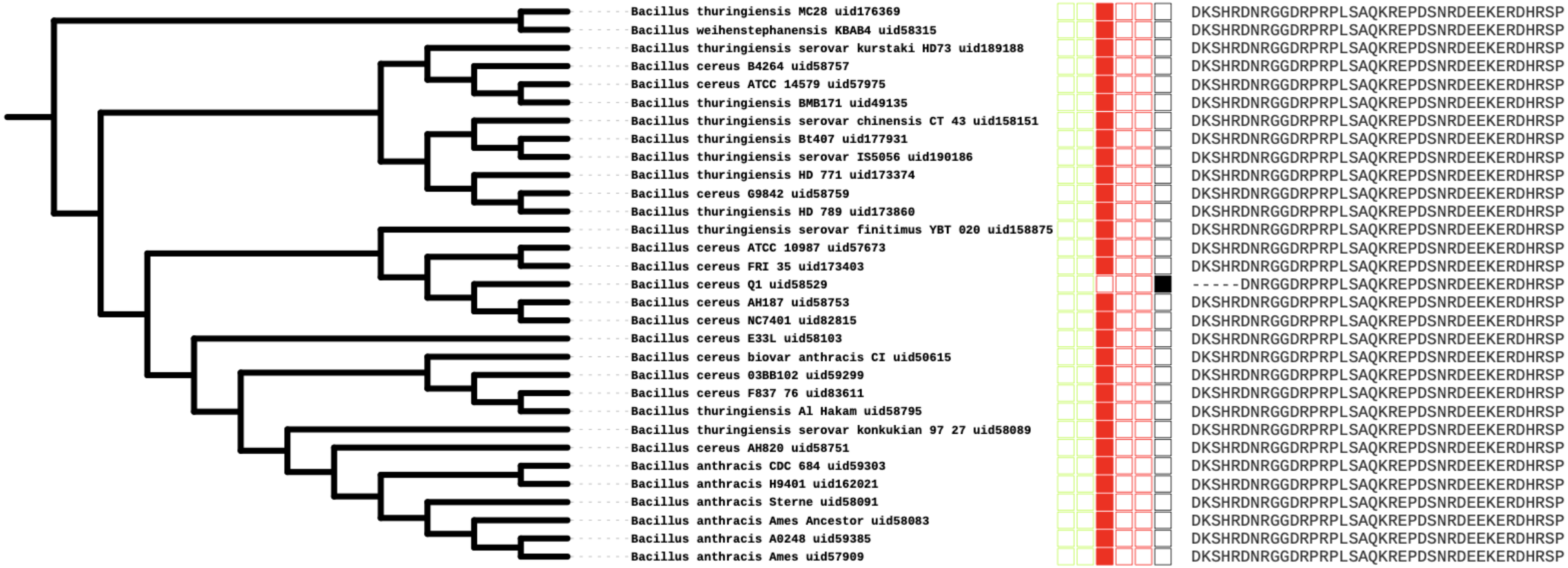

**Supplementary figure 12. Example of change from resistant (Class II) to unclassified in Bacillus (ATGC014) with few amino acid deletions in the active site of the EPSPS.** Phylogenetic trees of the ATGC014. Filled boxes show bacteria sensitive (green) and resistant (red) to glyphosate. Boxed in order from left to right correspond to Class I alpha, Class I beta, Class II, Class III, Class IV and Unclassified. The amino acids in the active site are shown at right side of the figure. Bacillus weihenstephanensis KBAB4 uid58315 has two copies of the sequence, one resistant to glyphosate (DKSHRDNRGGDRPRPLSAQKREPDSNRDEEKERDHRSP) and one unclassfied (DKSHR--RGGDRPRPLSAQKREPDSNRDEEKERDHRSP).

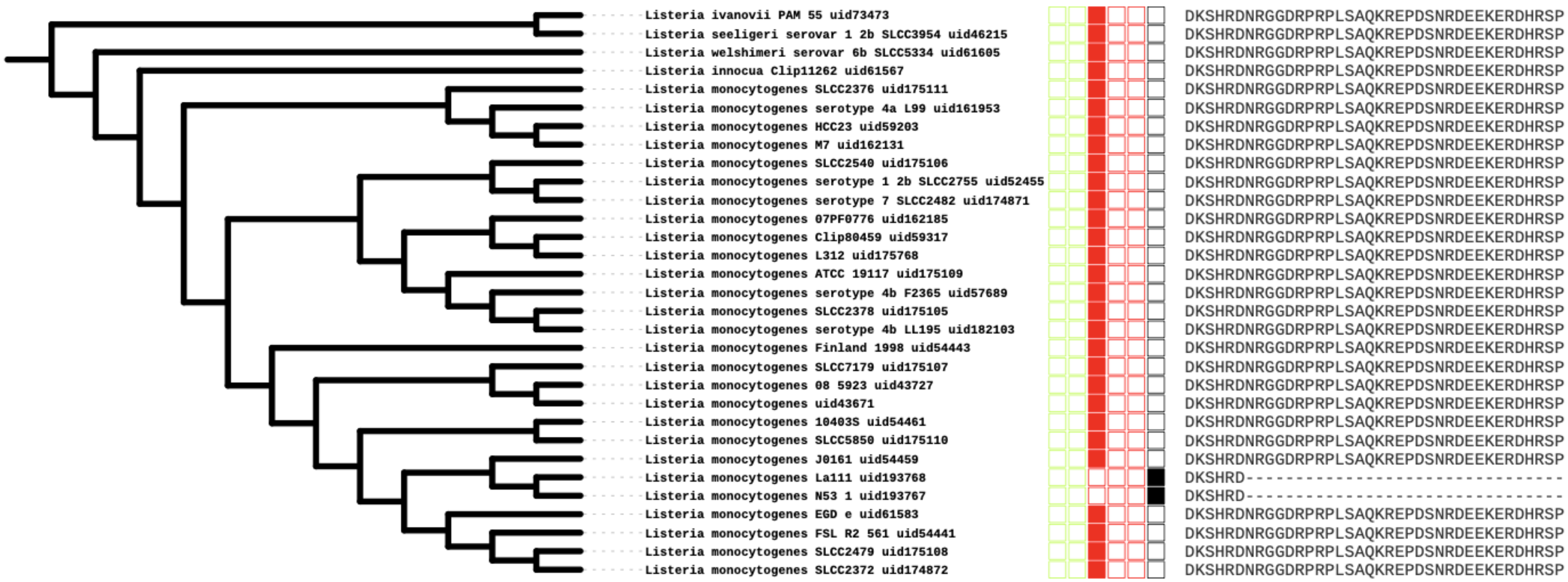

**Supplementary figure 13. Example of change from resistant (Class II) to unclassified in Listeria (ATGC109) with several amino acid deletions in the active site of the EPSPS.** Phylogenetic trees of the ATGC109. Filled boxes show bacteria sensitive (green) and resistant (red) to glyphosate. Boxed in order from left to right correspond to Class I alpha, Class I beta, Class II, Class III, Class IV and Unclassified. The amino acids in the active site are shown at right side of the figure.

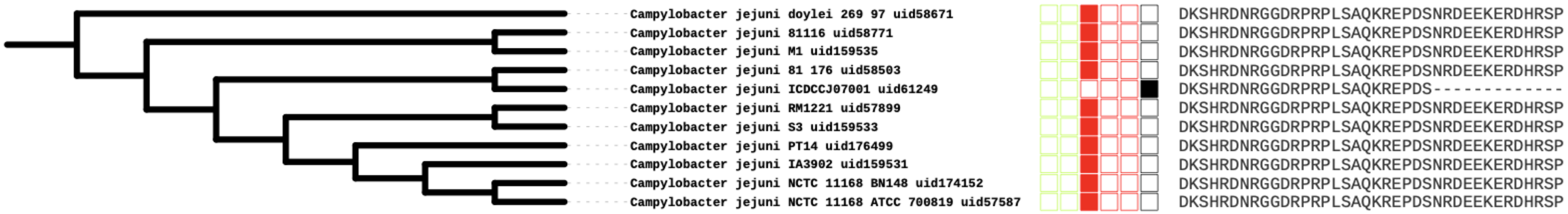

**Supplementary figure 14.Example of change from resistant (Class II) to unclassified in Campylobacter (ATGC144) with several amino acid deletions in the active site of the EPSPS.** Phylogenetic trees of the ATGC144. Filled boxes show bacteria sensitive (green) and resistant (red) to glyphosate. Boxed in order from left to right correspond to Class I alpha, Class I beta, Class II, Class III, Class IV and Unclassified. The amino acids in the active site are shown at right side of the figure.

a)

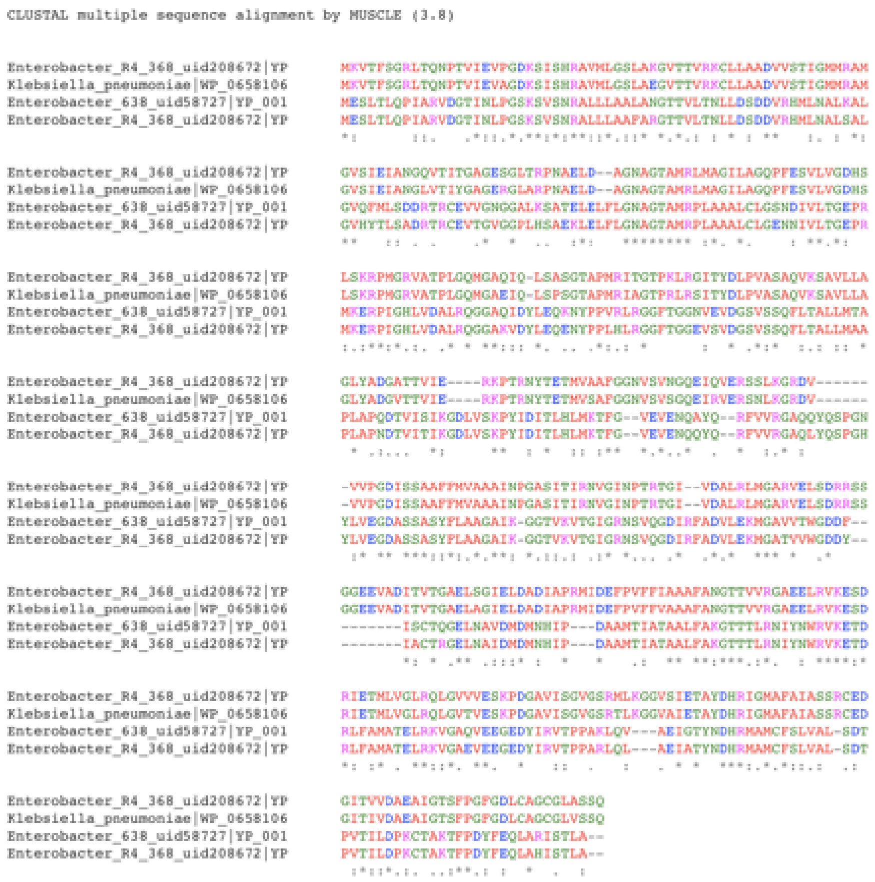

b)

| Species | Class I | Class II |
|---|---|---|
| Klebsiella pneumoniae (WP_065810684) | | X |
| Enterobacter_R4_368 (YP_008098272.1) | | X |
| Enterobacter_638 (YP_001176159.1) | X | |
| Enterobacter_R4_368_uid208672 (YP_008110400.1) | X | |

c)

Klebsiella pneumoniae (WP_065810684)

| CLASS | IDENTITY |
|---|---|
| Class I (alpha) | 0.800 |
| Class I (beta) | 0.889 |
| Class II | 1.000 |
| Class III | 0.346 |
| Class IV | 0.167 |

Enterobacter_R4_368 (YP_008098272.1)

| CLASS | IDENTITY |
|---|---|
| Class I (alpha) | 0.800 |
| Class I (beta) | 0.889 |
| Class II | 1.000 |
| Class III | 0.335 |
| Class IV | 0.167 |

Enterobacter_638 (YP_001176159.1)

| CLASS | IDENTITY |
|---|---|
| Class I (alpha) | 1.000 |
| Class I (beta) | 1.000 |
| Class II | 0.605 |
| Class III | 0.470 |
| Class IV | 0.233 |

Enterobacter_R4_368(YP_008110400.1)

| CLASS | IDENTITY |
|---|---|
| Class I (alpha) | 1.000 |
| Class I (beta) | 1.000 |
| Class II | 0.605 |
| Class III | 0.454 |
| Class IV | 0.233 |

**Supplementary figure 15. Putative acquisition of glyphosate resistance in *Enterobacter* R4 68 (ATGC001) *via* horizontal gene transfer from *Klebsiella pneumoniae*.**
a) Multiple sequence alignment of proteins. The sequence WP_065810684 from *K. pneumonia* was obtained as a result of the best BLAST hit (identity 100%).
b) Classification of the EPSPS enzyme into sensitive (class I) and resistant (Class II) to glyphosate
c) Identity values of the four proteins to the reference EPSPS sequences in *EPSPSClass* server (https://ppuigbo.me/programs/EPSPSClass/)

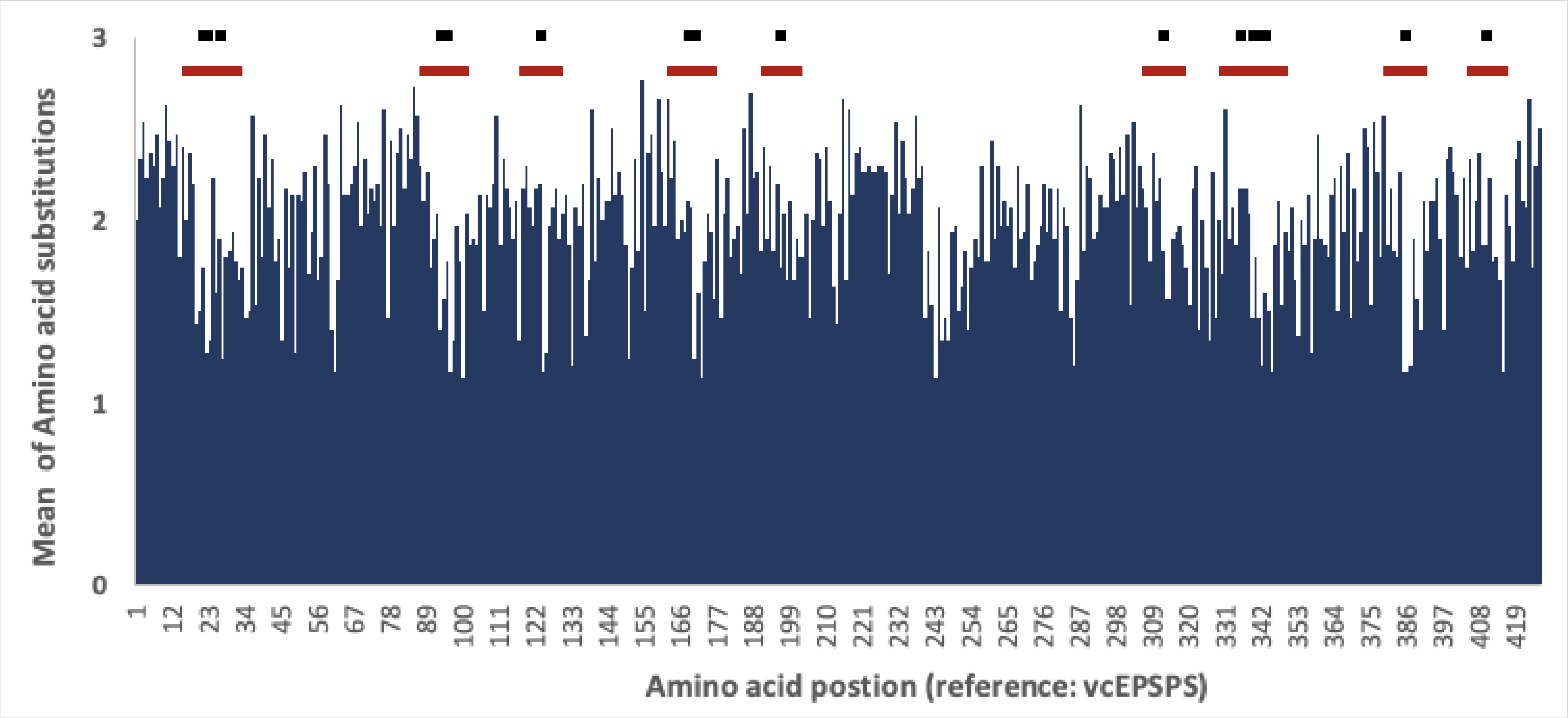

**Supplementary figure 16. Amino acids substitutions landscape.** Blue: Mean amino acid substitutions in the EPSPS sequences across ATGCs; Black: amino acids of the active site; red: active site region (active site +/-5 amino acids). Amino acids positions are based on the vcEPSPS sequence (supplementary table 6). Amino acids in the active site (black) of the EPSPS that bind are more conserved (Kruskal-Wallis test p-value $< 0.0001$) than the rest of amino acid residues. Also, amino acids around binding sites (+/- 5 residues) have higher levels of conservation (Kruskal-Wallis test p-value $= 0.005$).

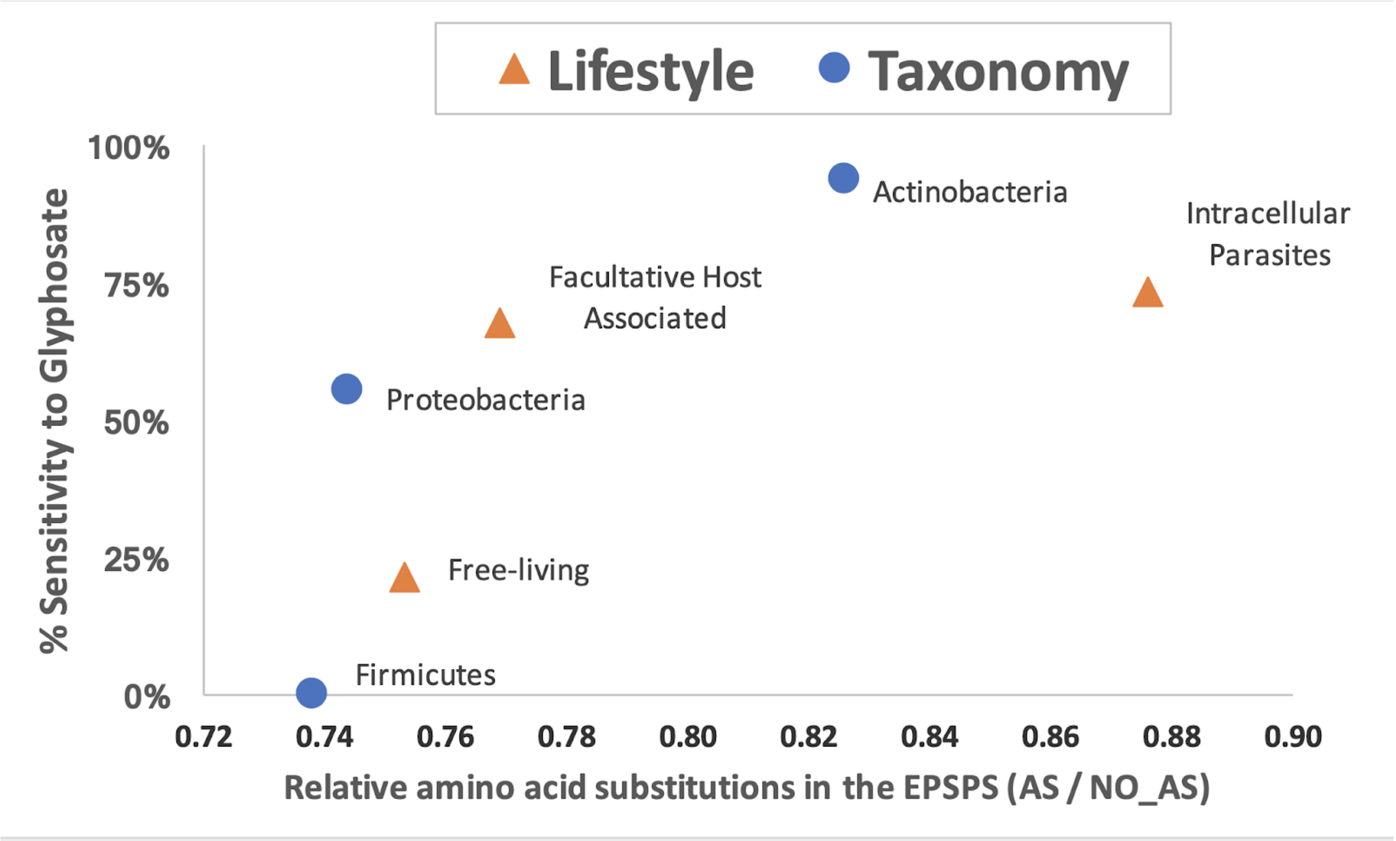

**Supplementary figure 17. Amino acid substitutions in the active site and sensitivity to glyphosate by taxonomy and lifestyle.** AS: amino acid substitution in the active site; No_AS: amino acid substitutions in other residues.
